# BI-TfR1 CapX rapidly transits the blood-brain barrier for efficient, low-dose gene delivery throughout the CNS

**DOI:** 10.64898/2026.09.23.753669

**Authors:** Ken Y. Chan, Chin-Yen Lin, Qin Huang, Simon Pacouret, Qingxia Zheng, John W. Harvey, Pam P. Brauer, Jencilin Johnston, Fiona Serack, Jessie R. Davis, Nikita G. Kamath, Saša Jereb, Daniel A. Sprague, Alexander Svanbergsson, Magalie Boucher, Eric V. Minikel, Sonia M. Vallabh, Benjamin E. Deverman

## Abstract

Safe, efficient gene delivery throughout the CNS remains a central obstacle to treating genetic diseases of the brain and spinal cord. We describe BI-TfR1 CapX, a human transferrin receptor (TfR1)-binding AAV capsid that, after a low intravenous dose in adult humanized *TFRC* mice, transduced more than 80% of cortical and spinal neurons and ∼60% of cortical astrocytes, with reduced distribution and expression in several peripheral tissues relative to AAV9. Unlike other engineered BBB-crossing capsids, CapX bound the CNS vasculature within minutes and a large fraction of its genomes reached parenchymal nuclei within 24 hours. Delivering a *Prnp*-targeted epigenetic silencer (*Prnp*-CHARM), CapX reduced brain *Prnp* mRNA by over 95% requiring 18-fold fewer vector genomes per brain cell than AAV-PHP.eB for 50% lowering. A CapX delivered dual-vector cytosine base editor installed a protein-truncating *PRNP* stop codon lowering brain prion protein by 80%. These properties make CapX an attractive vehicle for human CNS gene therapy.

## Introduction

Genetic and sporadic diseases of the CNS represent a vast area of unmet medical need. Gene therapy offers the prospect of durable, mechanistically targeted one-time treatments, but its broad application to CNS disease has been constrained by lack of human relevant vehicles for CNS-wide delivery. Adeno-associated viral (AAV) vectors remain the leading vehicles for gene supplementation, knockdown, and editing: they are small enough to spread through the CNS interstitium^1^, efficiently transduce neurons and glia^2–4^, and persist as episomes^5,6^, providing durable expression in long-lived cells. In addition, industrial manufacturing and clinical experience with these vectors has matured significantly over the past decade, resulting in greater production capacity and improved consistency and quality.

AAVs can be delivered to the CNS by several routes. Direct, MRI-guided intraparenchymal injection can target defined structures and underlies an approved gene therapy for AADC deficiency^7^, but most CNS diseases would benefit from broad CNS coverage^8–10^. Cerebrospinal fluid (CSF) routes of administration (intrathecal, intracerebroventricular, or intra-cisterna magna) offer wider distribution than direct injection and are in clinical testing, but diffusion-limited spread to CSF-adjacent parenchymal regions leaves structures unevenly transduced^11–15^. In contrast, because most neurons lie within 10-25 µm of the nearest capillary, intravenous administration offers whole-CNS access through a translationally scalable, noninvasive route. Clinically, this strategy has been realized through Zolgensma, a transformative AAV9 gene therapy for spinal muscular atrophy that has treated thousands of infants worldwide^16^. This therapy was enabled by the finding that AAV9 crosses the blood-brain barrier (BBB) in neonatal mice^2^ and other species^11–14,17^. Yet this hopeful precedent remains to be extended; intravenous AAV9 has proven less efficient in animals after the neonatal stage, and attempts to counter low efficiency with high doses can lead to serious adverse events.

Over the past decade, our group and others have shown that CNS tropism can be markedly improved through capsid engineering, where capsid variants that cross the BBB more efficiently in mice^3,18–21^ and non human primates (NHPs)^22,23^ are selected from highly diverse libraries. The tropism gains have generally been specific to the species used for selection, and translation to humans remains uncertain even for capsids selected in NHPs. One notable exception is the work of Moyer et al., who selected for CNS transduction in rodents and NHPs, and identified capsids that cross the BBB in both species through increased binding of ALPL, a highly conserved protein that was not previously known for its potential to aid in BBB crossing^23^. Capsids from this platform are now advancing to the clinic, with a first-in-human Phase I trial of one candidate (VY1706) actively enrolling patients with early Alzheimer’s disease (NCT07764146).

To circumvent the uncertainties of *de novo* selection in animals, we showed that AAVs can be directly reprogrammed to bind defined cell-surface receptors^24^, and we recently used this approach to create BI-hTFR1, a capsid that binds human transferrin receptor (TfR1)^24^. We chose TfR1 because it is highly and stably expressed on the human BBB across the lifespan^25^, mediates efficient transcytosis of iron-loaded transferrin into the brain, and is a clinically validated brain-delivery target: several TfR1-binding biologics reach the CNS, including AVLAYAH (tividenofusp alfa), an iduronate-2-sulfatase-Fc transport vehicle conjugate that was recently approved for Hunter syndrome^26^.

Although BI-hTFR1 was developed entirely with *in vitro* assays, it crosses the BBB and transduces the CNS of humanized TfR1 mice after intravenous delivery. In the present study, we matured BI-hTFR1 through mutagenesis and screening for TfR1-dependent functions *in vitro* and CNS transduction in humanized TfR1 gene knock-in (*hTFRC* KI) mice generating a second-generation variant, BI-TfR1 CapX, with substantially enhanced CNS transduction and reduced liver targeting. BI-TfR1 CapX transduces neurons and glia efficiently at low systemic doses, is manufacturable at yields comparable to AAV9 using standard methods, and rapidly binds and transits the brain endothelium, with less persistent vascular accumulation than other tested BBB crossing capsids. In a preliminary non-GLP study with a genome not encoding a transgene product, systemic BI-TfR1 CapX was well tolerated in *hTFRC* KI mice, with no discernible toxicity or adverse clinical pathology.

BI-TfR1 CapX retains the strict human-TfR1 specificity of BI-hTFR1. Using AlphaFold3^27^ and confirmatory receptor mutagenesis, we map its binding footprint on the TfR1 apical domain and identify a single human-specific residue that accounts for the species selectivity. In mice humanized for both *TFRC* and the prion protein gene (*PRNP*), we show that BI-TfR1 CapX can efficiently deliver a base editor^28^ split across two vectors to install a prion protein-truncating edit. Across a range of doses, we also compared the ability of BI-TfR1 CapX and a surrogate capsid AAV-PHP.eB to deliver the *Prnp*-CHARM epigenetic editor^29^, achieving near-complete brain-wide *Prnp* silencing and a ∼2-fold increase in potency over AAV-PHP.eB. Remarkably, BI-TfR1 CapX reaches 50% brain *Prnp* mRNA lowering with 18-fold fewer vector genomes per brain cell than AAV-PHP.eB, indicating a substantially higher potency per particle. Together with its rapid BBB crossing and low endothelial accumulation, these results establish BI-TfR1 CapX as a promising candidate vehicle for human CNS gene therapy.

## Results

### Maturation of BI-hTFR1 through SSM and *in vivo* screening

Our group’s initial reprogramming of AAV9 to bind human TfR1 yielded BI-hTFR1, a capsid that crosses the blood-brain barrier in *hTFRC* KI mice but that requires a moderately high systemic dose (∼2.5 × 10^13^ vg/kg) to transduce ∼50% of cortical neurons^24^. To mature BI-hTFR1 into a more efficient vehicle, we diversified the capsid and screened for variants with enhanced human-TfR1 binding and CNS transduction in a single round of combined *in vitro* and *in vivo* selection. We generated a focused library (∼7,900 variants) by single-site-saturation mutagenesis (SSM) of the BI-hTFR1 7-mer insertion (YSRIGPN) and dual-SSM of the flanking loop VIII residues (Figure 1A). As benchmarks we included the mouse BBB-crossing capsids AAV-PHP.eB^19^, 9P31^20^, AAV-BI28^18^ and AAV-BI30^30^, along with three sets of AAV9 sequences printed at 1x, 10x, and 100x copies, to improve the quantitative recovery of this benchmark capsid in assays where its performance is low. Each variant was encoded by two synonymous nucleotide sequences to provide internal replicates, and the library was cloned into two expression cassettes: a CMV-CBA-driven cassette for *in vitro* selection and a human synapsin (hSYN1)-driven cassette designed to add selective pressure *in vivo* for capsids that both cross the BBB and transduce neurons. *In vitro* pull-downs against purified hTfR1-Fc proteins demonstrated that 48% of variants retained selective receptor binding over the Fc-only control (mean of synonymous nucleotide log_2_ enrichment ≥ 1; Figure S1A). We then screened the library in *hTFRC* KI mice for brain transduction after intravenous delivery. We analyzed the AAV9 series to find the representation at which its brain enrichment estimate became reliable. Agreement between synonymous replicate nucleotide pairs tightened as copy number rose (enrichment-score differences of 4.76, 0.16, and 0.04 for 1x, 10x, and 100x), so we used the 10x AAV9 mean as the comparator throughout. Of the hTfR1 binding variants, 44% were enriched above AAV9 in the brain (Figure 1B), and brain enrichment was highly reproducible across synonymous pairs (r = 0.88, Log_2_ enrichment ≥ 1; Figure 1C). BI-hTFR1 outperformed AAV9 by 70x and was the single best-performing 7-mer SSM variant (Figure 1C), confirming that the previous *in vitro* screen had already reached a local fitness peak.

**Figure 1.**
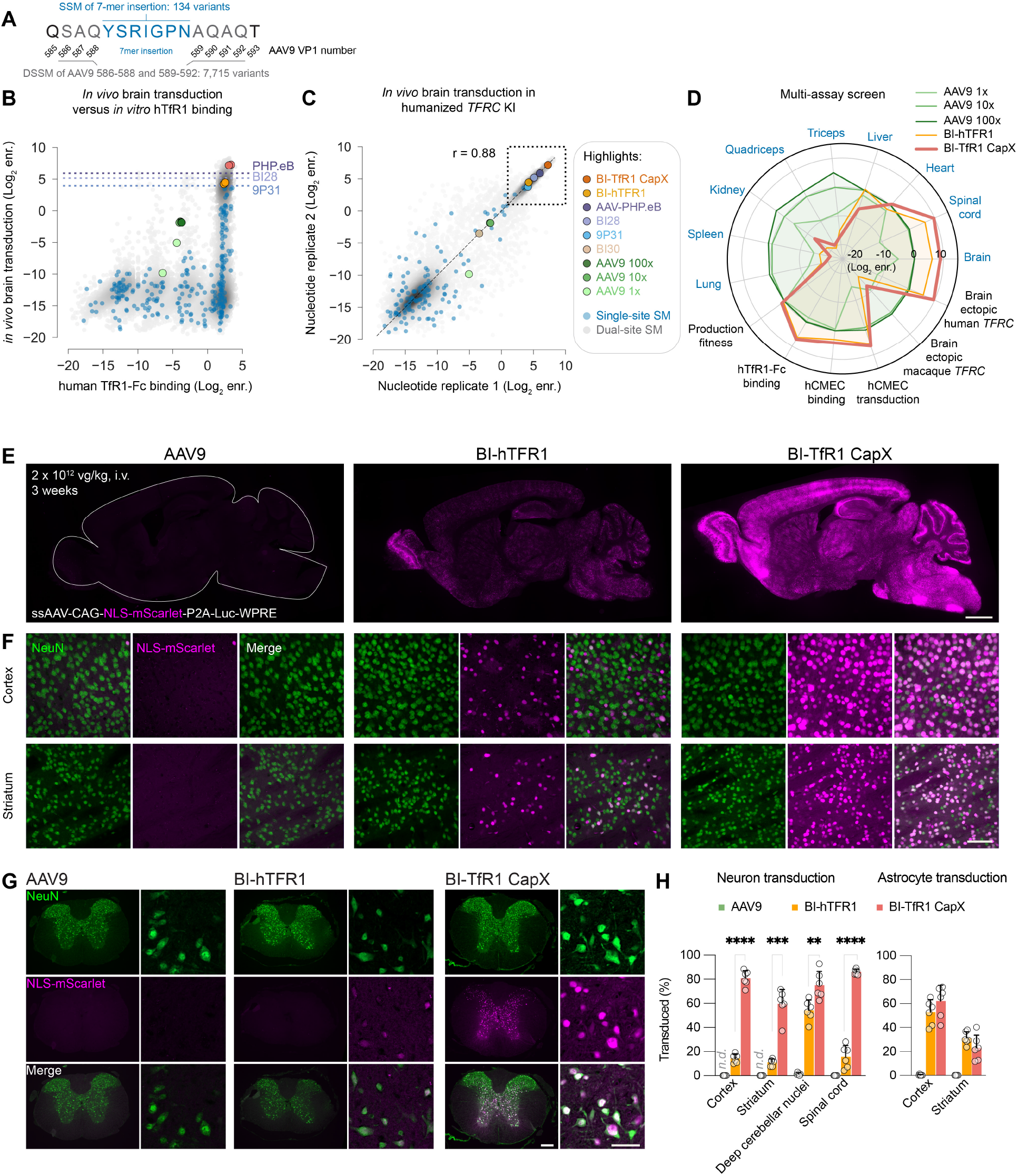
Identification of BI-TfR1 CapX, a second-generation capsid that enables efficient and widespread gene delivery to the CNS after low dose systemic administration. (**A**) Schematic representation of the amino acids diversified during single- and dual-site saturated mutagenesis (SSM, original BI-hTFR1 7-mer blue; DSSM, gray; and surrounding AAV9 amino acids, black). (**B**) CNS transduction in *hTFRC* KI mice versus hTfR1-Fc binding *in vitro* (log_2_ enrichment (enr.)). The dotted lines indicate AAV-PHP.eB, BI28, and 9P31 *in vivo* performance; the BBB crossing reference capsids were not included in the *in vitro* library (**C**) Reproducibility of brain transduction in *hTFRC* KI mice between the replicate synonymous nucleotide sequences encoding each capsid in the library. Pearson correlation is shown for variants with log_2_ enrichment ≥ 1. D) Mean log_2_ enrichment (across biological replicates and synonymous nucleotide encodings), normalized to AAV9 10x for production fitness, *in vitro* assays of TfR1-dependent function (hCMEC/D3 binding and transduction and hTfR1-Fc binding) and *in vivo* transduction measurements across multiple organs (blue) in *hTFRC* KI mice and in NSG mice made to express ectopic human or rhesus macaque *TFRC* with AAV-BI30:CAG-*hTFRC* or AAV-BI30:CAG-*rhTFRC. In vivo* screens were performed in mice administered 2 × 10^11^ library viral genomes (vg) intravenously (i.v.) and assessed at 3 weeks. (**E**-**H**) AAV9, BI-hTFR1, or CapX carrying a ssAAV-CAG-NLS-mScarlet-P2A-luc-WPRE-pA genome were i.v. administered to adult *hTFRC* KI mice at 2 × 10^12^ vg/kg (n = 6 mice per group). (**E**) Representative images show native mScarlet fluorescence in brain sagittal sections (magenta). (**F** and **G**) Confocal microscopy images show NeuN^+^ neurons (green) and native NLS-mScarlet fluorescence (magenta) 3 weeks post administration in the cortex and striatum (**F**) and the spinal cord (**G**). (**H**) The percentage of NeuN^+^ cells that expressed NLS-mScarlet in the cerebral cortex, striatum, deep cerebellar nuclei (DCN), or spinal cord (left). The percentage of SOX9^+^ astrocytes that expressed NLS-mScarlet (right). AAV9 served as a benchmark control and was not included in statistical testing; *n*.*d*. denotes no detection of NLS-mScarlet^+^ nuclei in any of the 6 animals. Each data point represents an individual animal and bars show the mean with SD (unpaired two-tailed Welch’s t test; *p < 0.05, **p < 0.01, ***p < 0.001, ****p < 0.0001). Scale bars: 1 mm (**E**), 50 µm (**F**), 500 µm (**G**, left panel) 50 µm (**G**, right panel).

A portion of the dual-SSM (DSSM) variants ascended above the BI-hTFR1 peak: multiple variants carrying flanking mutations were reproducibly more enriched for brain transduction than BI-hTFR1 and each of the benchmark BBB crossing capsids (Figure 1C). None engaged rhesus macaque TfR1 in an *in vivo* ectopic-expression assay (Figure S1B-D, see Methods for details), indicating that the human specificity of the 7-mer was not relaxed by this focused mutagenesis. With many strong candidates in hand, we prioritized additional properties needed for a clinical vehicle: production fitness, reduced off-target enrichment in liver and other organs, and binding and transduction of human brain endothelial cells (hCMEC/D3) (Figure 1D and S1E). On this basis we selected a lead, BI-TfR1 CapX (hereafter referred to as CapX for simplicity), which retains the YSRIGPN peptide and adds two flanking substitutions, S586E and A596N. CapX outperformed AAV9, 9P31, AAV-PHP.eB, BI28, and BI-hTFR1 for brain transduction in *hTFRC* KI mice by 514x, 9.6x, 2.4x, 4.2x, and 7.3x, respectively; maintained high production fitness; and showed reduced enrichment in liver and other non-CNS organs, with the exception of heart, relative to AAV9 (Figure 1D).

### CapX enables highly efficient transduction of the adult CNS at a low systemic dose

To characterize CapX individually, we packaged a single-stranded reporter genome expressing mScarlet and with a nuclear localization sequence (NLS) and firefly luciferase (luc) from a strong ubiquitous promoter (ssAAV-CAG-NLS-mScarlet-P2A-luc-WPRE) and, in parallel, packaged the same genome into AAV9 and BI-hTFR1. A low dose of each vector was administered intravenously at 2 × 10^12^ vg/ kg to adult *hTFRC* KI mice. Three weeks later, we assessed native mScarlet fluorescence throughout the brain and spinal cord and found that mScarlet expression was markedly higher in CapX-treated animals than in those that received BI-hTFR1 or AAV9 (Figure 1E-G; Figure S2).

Quantifying neuronal transduction as the fraction of NeuN^+^ nuclei co-labeled with mScarlet, CapX transduced significantly more neurons than BI-hTFR1 across every CNS region examined (cortex, 81% vs 14%, 5.7x; striatum, 60% vs 11%, 5.4x; deep cerebellar nuclei, 75% vs 54%, 1.3x; spinal cord, 86% vs 16%, 5.4x). At this low dose, transduction by AAV9 in the cortex, striatum and spinal cord was minimal or undetected (Figure 1H). Astrocyte transduction (mScarlet^+^/SOX9^+^) was comparable between CapX and BI-hTFR1 (cortex, 61% vs 52%; striatum, 23% vs 31%), whereas AAV9 transduced under 1% of astrocytes in both regions. A single round of maturation thus converted BI-hTFR1 into a capsid that transduces the majority of neurons across the brain and spinal cord at a low systemic dose.

### The enhanced tropism of CapX is CNS-specific

We next measured biodistribution as viral genomes per diploid genome (vg/dg) across the CNS and major organs (Figure 2A). CapX reached higher CNS biodistribution than either comparator (brain, 2.8 vs 1.4 vs 0.016 vg/dg; spinal cord, 5.2 vs 3.3 vs 0.048 vg/dg for CapX, BI-hTFR1, and AAV9), corresponding to a ∼2x increase over BI-hTFR1 in brain (with no significant difference in spinal cord) and ∼175x and ∼108x increases over AAV9 in brain and spinal cord, respectively. Liver biodistribution was lower for CapX (4.7 vg/dg) than for BI-hTFR1 (21.0 vg/dg) but not significantly different than AAV9 (14.4 vg/ dg). Because hepatic AAV transduction is influenced by sex through an androgen-dependent pathway^31^, we also analyzed liver biodistribution by sex (Figure S3A-B). CapX liver biodistribution was reduced relative to both BI-hTFR1 and AAV9, but this reduction was significant only in males. Neither CNS biodistribution nor transduction differed by sex for any capsid (Figure S3C-D). Notably, CapX distributed to the CNS and liver with similar efficiency, whereas AAV9 delivered ∼900-fold and ∼300-fold more genomes to the liver than to the brain and spinal cord, respectively (Figure 2A).

**Figure 2.**
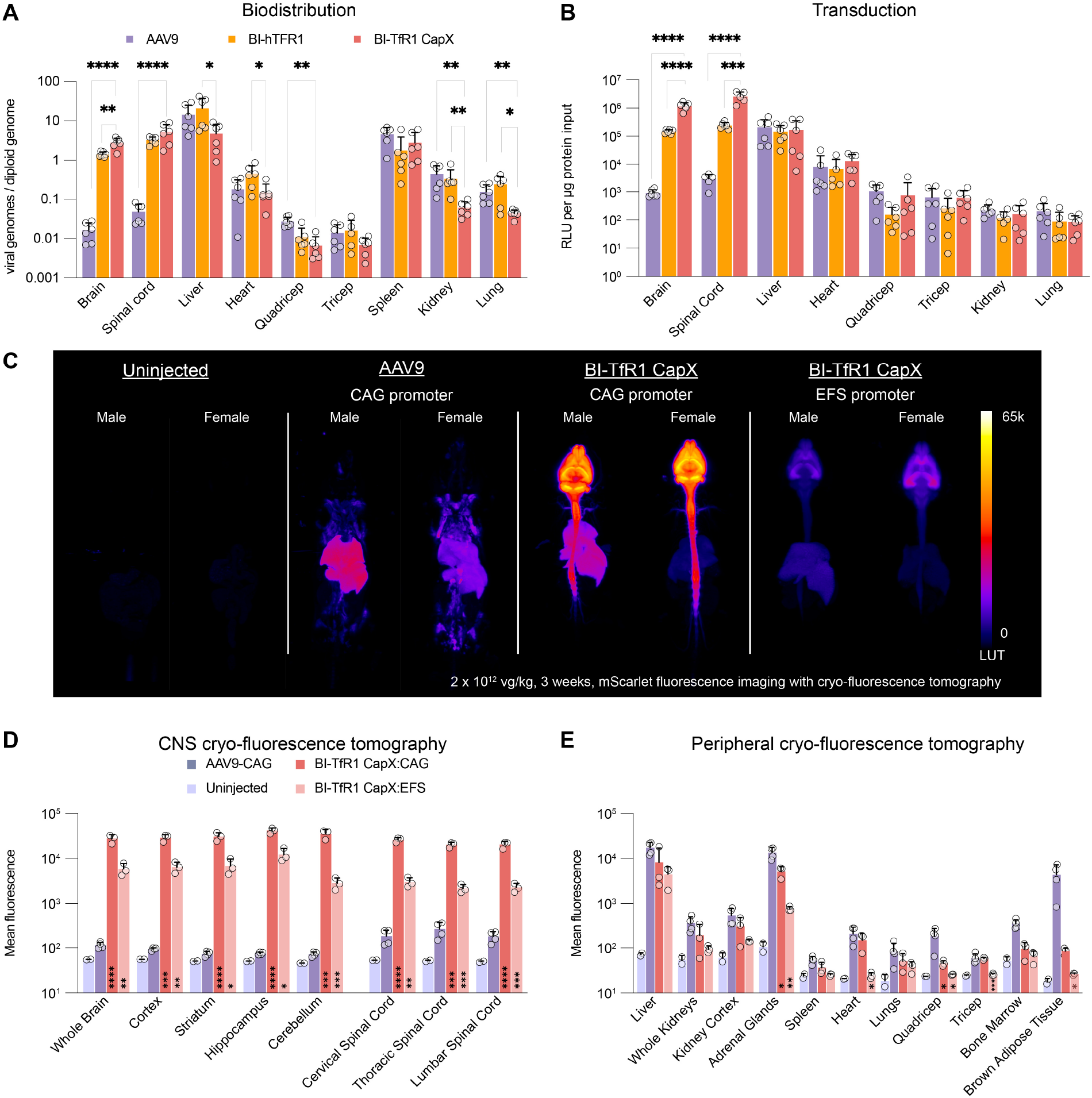
The enhanced tropism of CapX is CNS-specific and detargeted from multiple peripheral organs. AAV9, BI-hTFR1, or CapX carrying a ssAAV-CAG-NLS-mScarlet-P2A-luc-WPRE-pA genome were i.v. administered to adult *hTFRC* KI mice at 2 × 10^12^ vg/kg (n = 6 mice per group). (**A**) Biodistribution (viral genomes per diploid genome, vg/dg) was assessed by qPCR. (**B**) Transduction was assessed by luciferase activity (RLU) per µg protein in tissue homogenates. (**C**) Images show representative cryo-fluorescence tomography projections of native mScarlet fluorescence in *hTFRC* KI mice three weeks after administration of 2 × 10^12^ vg/kg of the indicated AAV (n = 3-4 per capsid and n = 2 uninjected controls). (**D** and **E**) The graphs show quantification of mean mScarlet fluorescence in segmented CNS regions (**D**) and non-CNS organs (**E**). In (**A** and **B**), capsids were compared within each organ. In (**D** and **E**), AAV9:CAG vs. CapX:CAG and CapX:CAG vs. CapX:EFS were compared within each brain region or organ. Each data point represents an individual mouse. Bars show the mean with SD. All comparisons were made using Brown-Forsythe and Welch ANOVA on log-transformed data, followed by Šídák’s multiple comparisons test (*p < 0.05, **p < 0.01, ***p < 0.001, ****p < 0.0001).

The enhanced CNS tropism seen by biodistribution of CapX as compared to BI-hTFR1 and AAV9 was even more evident when assessing expression of the luciferase reporter transgene. Transgene expression in the brain from CapX was 8x greater than that achieved by BI-hTFR1 and 1,300x greater than AAV9; in spinal cord the differences were 10.6x and ∼885x, respectively (Figure 2B). We observed no significant difference in transgene expression in the liver among the three capsids and no increase in biodistribution or transgene expression in any organ outside the CNS, establishing that the enhanced tropism of CapX is CNS-specific.

### Unbiased whole-body transduction assessment with cryo-fluorescence tomography

To assess whole-body transduction without regional bias, we performed cryo-fluorescence tomography (CFT)^32^ of mScarlet reporter expression in adult *hTFRC* KI mice that received ssAAV-CAG-mScarlet-2A-luc-WPRE-pA delivered by CapX or AAV9 at 2 × 10^12^ vg/kg. Native mScarlet fluorescence was imaged across the whole body three weeks later (Figure 2C). A parallel cohort that received CapX carrying the same reporter driven by a short fragment of the EF1a promoter (EFS) showed a similar distribution of mScarlet expression, though as expected, at a lower intensity.

Horizontal projections (Figure 2C) and three-dimensional reconstructions (Video S1-6) showed that CapX transduction was overwhelmingly CNS-directed, with whole-brain mScarlet fluorescence 242x that of AAV9 and regional increases of 306x, 416x, 547x, and 484x in the cortex, striatum, hippocampus, and cerebellum (Figure 2D), respectively, consistent with our luciferase reporter data. The enhanced tropism was CNS-restricted and no organ outside the CNS showed increased fluorescence relative to AAV9. Expression from CapX was significantly lower than from AAV9 in the adrenal glands, quadriceps, and brown adipose tissue (2.6x, 4.0x, and 51x, respectively). Trends toward reductions in the bone marrow and liver (3.7x and 2.0x, respectively) were observed but did not reach statistical significance (Figure 2E).

### The CNS tropism of CapX is consistent across humanized *TFRC* KI models

Multiple partially humanized *TFRC* knock-in mouse lines are now commercially available. Throughout our prior study and this work, we have used B-hTFR1 mice (referred to as *hTFRC* KI mice), which have the entire TfR1 extracellular domain humanized (Biocytogen, 110861). Because BI-hTFR1 binds the TfR1 apical domain^24^, we predicted that CapX would also be compatible with an additional mouse line that is humanized only within the apical domain (hAPI)^33^. Indeed, CapX transduced the two lines with similar efficiency (Figure S4), confirming that either line is suitable for preclinical evaluation of CapX-based gene therapies.

### TfR1 binding by CapX requires a human-unique apical domain residue

To define the basis of CapX’s human specificity, we used AlphaFold3 (AF3)^27^ to predict a docking pose of a CapX trimer with three chains of a human soluble TfR1 apical-domain fragment^34^ (TfR1sol-apical) (Figure 3A) and supported the model with confirmatory receptor mutagenesis. In the top-ranked model, AF3 predicted a pose in which CapX engages TfR1sol-apical with each residue of the engineered peptide (AA 586–596) within 5 Å of the receptor (Figure 3B). The top-ranked model had moderate confidence at the interface: an ipTM of 0.64 (interface predicted TM-score; 0-1, confidence increases with the score) and a minimum inter-chain predicted aligned error (PAE_min) of 6.13 Å (0-32 Å) (Figure 3C). A residue-level PAE heatmap (Figure S5A) confirmed that the lowest inter-chain PAE values for the docking interaction are concentrated at the engineered loop region (CapX residues 586-596) that was predicted to be in close contact with hTfR1sol-apical (Figure S5A). The docking poses were consistent across 11 seeds as assessed by low alpha carbon root mean square deviations (median RMSD = 3.3 Å) between each model’s TfR1sol-apical chain after capsid-subunit alignment to the top-ranked model (Figure 3C). We therefore treat the docking pose as a hypothesis-generating model.

**Figure 3.**
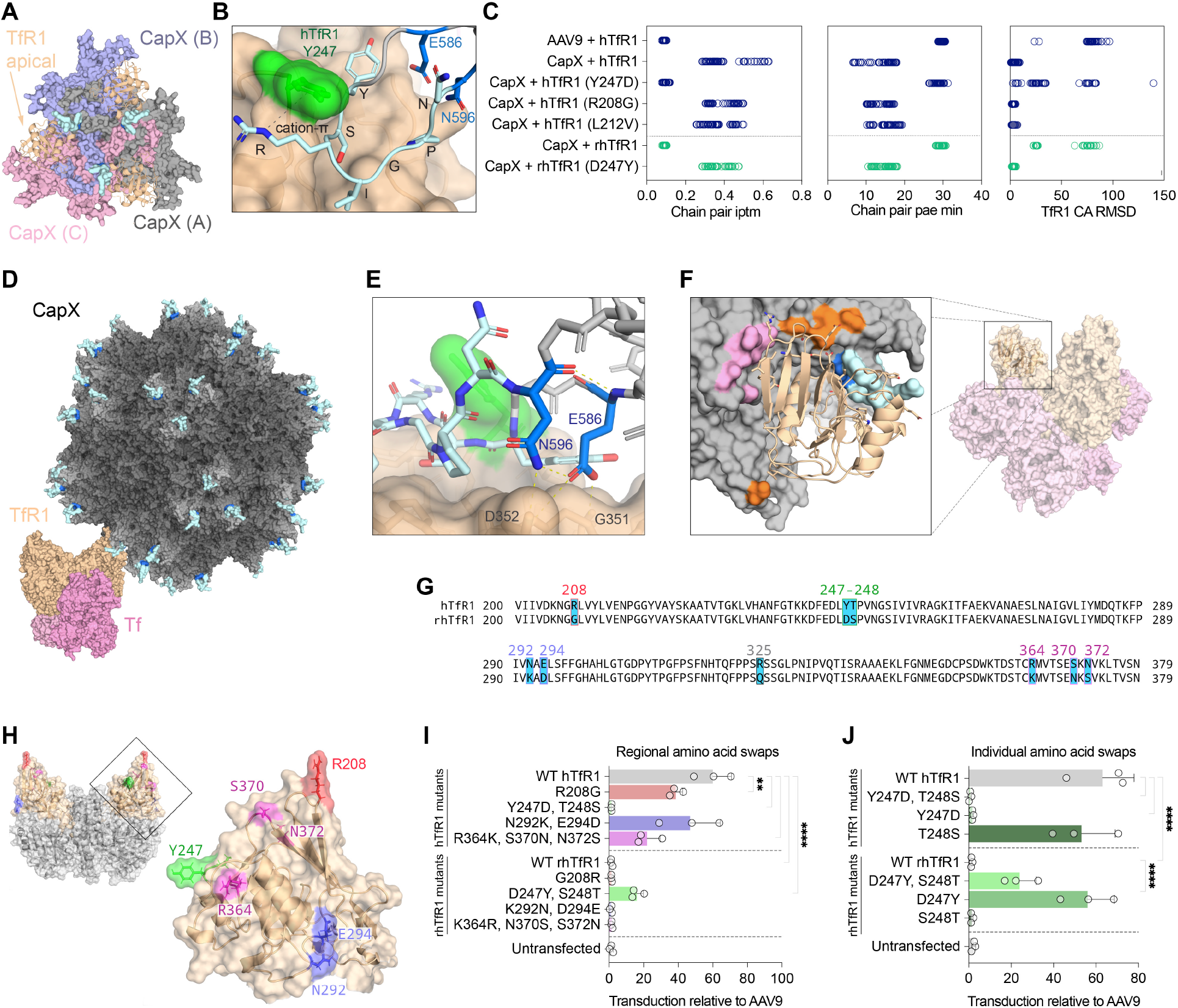
Structural insights into the human TfR1-specific binding of CapX. (**A**) AF3 predicted structural model of the docking of the AAV CapX trimer (gray, pink, blue) and three human TfR1 apical domain fragments (wheat). (**B**) The top ranked AF3 model predicts that the 7-mer peptide (cyan) and CapX-specific flanking substitutions (blue) are in close contact with the TfR1 apical domain. Arg591 (YSRIGPN) is predicted to make a cation-pi interaction with the human-specific TfR1 residue Tyr247 (green). (**C**) AF3 chain pair IPTM (left), predicted chain pair aligned error (pae min, middle) scores for the indicated capsid and TfR1 docking models are shown. The plot (right) shows model consistency across 55 models (11 seeds) based on the alpha carbon (Cα) RMSD of the TfR1 apical chain after alignment of CapX trimer. (**D**) An AF3 docking model was aligned to an assembled AAV9 model (PDB ID: 3UX1) and the soluble apical domain fragment was aligned with a TfR1-transferrin (Tf) complex (PDB ID: 1SUV). The model shows CapX in gray depth shading with the peptide insertions (cyan) along with the globular TfR1 ectodomain (wheat) and two Tf units (pink). (E) The two CapX specific substitutions (E586 and N596, blue) are predicted to form hydrogen bonds with TfR1 and each other (dotted lines). (**F**) All three subunits of the CapX trimer are predicted to contact the TfR1 apical domain (wheat). The space fill model highlights the targeting peptide (teal and blue) and predicted contact residues (within 4 angstroms) from the other two capsid subunits (orange and pink). (Right) The TfR1sol-apical fragment (cartoon) superimposed with the TfR1-Tf complex (spacefill; PDB ID: 1SUV). (**G**) Alignment of the human and rhesus macaque (rh)TfR1 apical domain AA sequences. (**H**) AF model of TfR1 highlighting residues that differ between human and rhesus macaque. (**I** and **J**) CHO cells transfected with the indicated TfR1 variant color coded to match (**G** and **H**) were transduced 2 days later with CapX and luciferase activity was measured 2 days later. (**J**) Transduction assay with individual human and rhesus macaque amino swaps. Bars show the mean with SD (representative experiment with three technical replicates, represented by data points). All comparisons were made using ordinary one-way ANOVA test followed by Šídák’s multiple comparisons test correction (**p < 0.01, ****p < 0.0001).

Aligning the docked receptor to a previously reported TfR1-transferrin (Tf) complex and the capsid subunits to a complete AAV9 capsid produced no clashes with the TfR1-Tf complex (Figure 3D), consistent with our prior observation that the parental BI-hTFR1 binds TfR1 while it is loaded with Tf^24^. The model predicts that the two substitutions that distinguish CapX from BI-hTFR1 (Ser586Glu and Ala596Asn) may form additional hydrogen bonds with TfR1 Gly351 and Asp352 and with each other (Figure 3E). The model also predicts that all three capsid subunits contribute potential contacts defined by proximity within 4 Å of the receptor (Figure 3F).

Interestingly, the model predicts that the engineered YSRIGPN peptide docks as an extended loop into a pocket on the apical domain, anchored by a cation-π interaction between CapX Arg591 and human TfR1 Tyr247 (4.9 Å from the Arg591 guanidinium proximal nitrogen (Nε) to the Tyr247 ring centroid). Because a Tyr at position 247 is unique to humans (Figure S5B), this interaction provides a structural rationale for the capsid’s species selectivity. Supporting the importance of Tyr247, AF3 returned low-confidence, inconsistent poses for the rhesus macaque apical domain (rhTfR1sol-apical) and for human TfR1sol-apical carrying a single Tyr247Asp substitution (Asp being the macaque residue). After capsid chain alignment between models, the median receptor-chain RMSD relative to the top human model was 62.8 Å and 26.3 Å for rhTfR1sol-apical and hTfR1sol-apical (Tyr247Asp), respectively (Figure 3C). Installing the reciprocal Asp247Tyr substitution onto rhTfR1sol-apical restored a consistent WT hTfR1-like pose (median RMSD = 2.6 Å), whereas the median RMSD for AAV9 with hTfR1 were inconsistent as expected (77.8 Å; Figure 3C).

To test the requirement for Tyr247 directly, we performed a receptor residue-swap experiment. Guided by an alignment of the human and macaque apical domains (Figure 3G), we generated human TfR1 constructs carrying one to three of the surface-exposed residues (Figure 3H) that differ from macaque. The WT or modified receptors were expressed in CHO cells and the cells were then transduced with CapX or AAV9 encoding a luciferase reporter. In cells transfected with WT human TfR1, the CapX transduction readout by the luciferase reporter was >50x higher than AAV9, whereas no enhancement of CapX transduction was observed with WT macaque TfR1 relative to untransfected cells (Figure 3I). The enhanced transduction from CapX was abolished in cells transfected with human TfR1 modified with both Tyr247Asp and Thr248Ser residue swaps, whereas the other residue sets caused at most partial reductions. Reciprocally, the humanizing Asp247Tyr and Ser248Thr swaps on macaque TfR1 raised relative CapX transduction ∼16-fold, whereas other humanizing sets did not. Testing the residues individually confirmed that Tyr247 alone is necessary and sufficient: Tyr247Asp on human TfR1 abolished the relative CapX transduction enhancement Asp247Tyr restored it to near WT human TfR1 levels, and swapping residues at position 248 in either human or macaque TfR1 had no effect (Figure 3J).

Although single amino-acid differences between species can have dramatic effects on binding, human polymorphism at this interface is minimal. Across the human TfR1 residues predicted to be within 5 Å of CapX, all missense variants are rare (maximum allele frequency 8.2 × 10^-4^; none exceed 1 × 10^-3^), indicating that common human *TFRC* variation is unlikely to affect the CapX binding interface (Table S1).

### CapX binds the CNS vasculature within minutes and efficiently reaches parenchymal nuclei

We next evaluated the kinetics of CNS targeting, BBB transit, and clearance from circulation exhibited by CapX relative to AAV9 and the engineered capsids AAV-PHP.eB^19^ and 9P31^20^, which use LY6A^35,36^ and carbonic anhydrase IV (Car4)^37^ for transport into the CNS, respectively. We packaged unique barcoded genomes into each capsid, enabling individual detection by either dPCR or by *in situ* hybridization sense probes designed to detect DNA, rather than mRNA, without crosstalk (Figure S6A-B). Each barcoded AAV was manufactured individually, pooled at equal viral genomes, re-titered to confirm equal concentration of each capsid within the injection cocktail (Figure S6C), and dosed at 2 × 10^12^ vg/kg per AAV (8 × 10^12^ vg/kg total AAV pool) into *hTFRC* KI mice, which are on a C57BL/6 background and express endogenous *Ly6a* and *Car4*. The brain, liver, serum, and blood were collected at 15 min, 2 hours, 24 hours, 3 days, 7 days, and 21 days post delivery. CapX rapidly accumulated in the brain, achieving a peak of 50 vg/dg by 15 min (Figure 4A), which was more than ∼100x greater than the peak AAV9 level (consistent with the biodistribution data, Figure 2A). AAV-PHP.eB and 9P31 peaked by 2 hours reaching ∼250 vg/dg and ∼20 vg/dg, respectively. In the liver, each of the engineered capsids reached peak levels by 2 hours while AAV9 continued to accumulate through 24 hours (Figure 4B).

**Figure 4.**
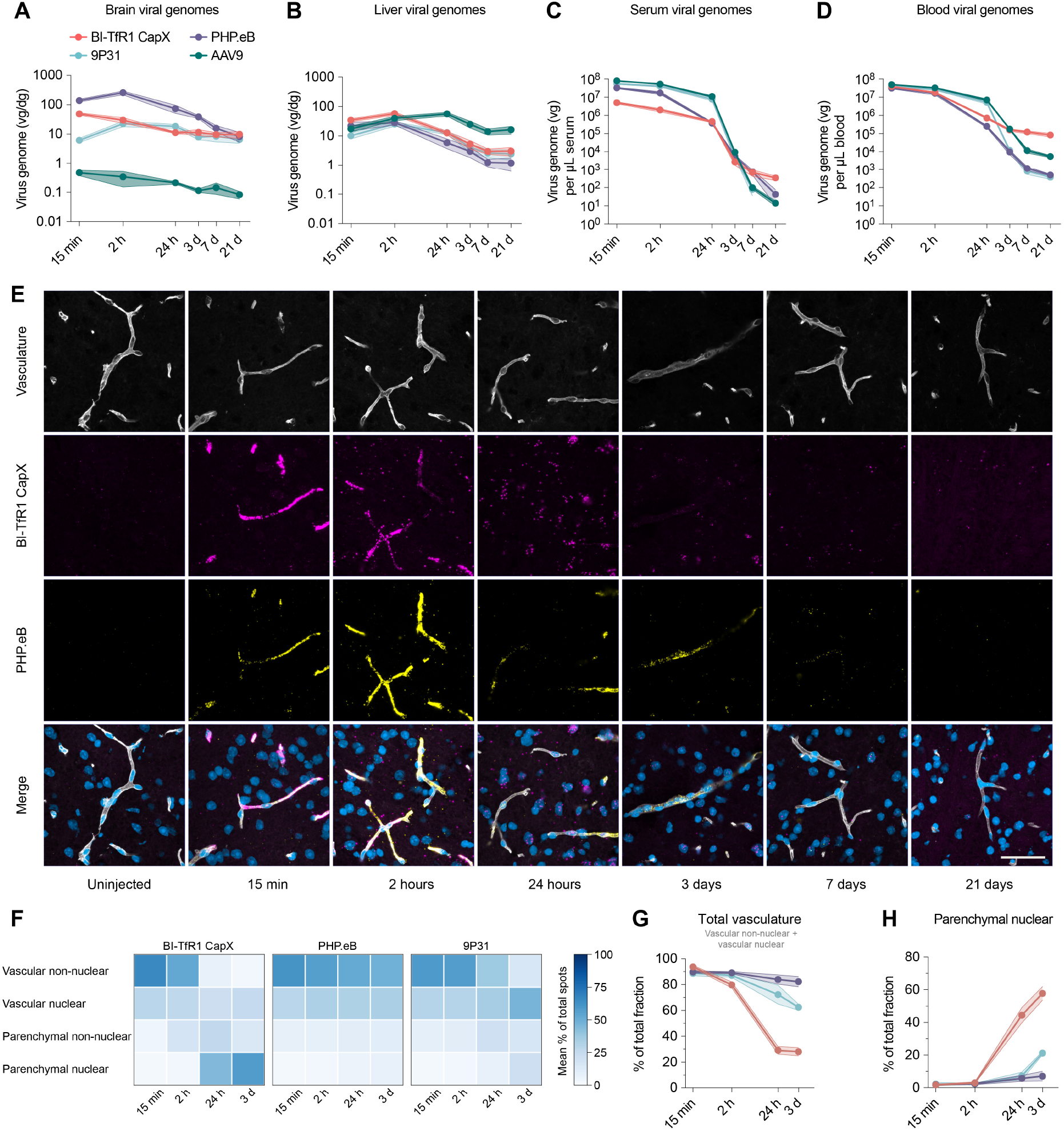
CapX rapidly crosses the BBB and enters parenchymal nuclei. A cocktail of four AAVs: CapX, AAV-PHP.eB, 9P31 and AAV9 each packaging a unique superfolder GFP expressing barcoded genome (ssCAG-H2B-sfGFP-WPRE-barcode-pA) were intravenously administered to *hTFRC* KI mice at a dose of 2 × 10^12^ vg/kg per barcoded AAV (8 × 10^12^ vg/kg total AAV pool). Tissues, serum, and blood were collected at the indicated timepoints. (**A**-**D**) dPCR assessments of AAV genomes are reported per diploid genome (vg/dg) for brain (**A**) and liver (**B**) and vg/µL for serum (**C**) and whole blood (**D**). (**E**) Representative RNAscope images from the brain at different time points post administration of the AAV pool. Images show the vasculature (lycopersicon esculentum lectin, white), CapX genomes (magenta), AAV-PHP.eB (yellow), and nuclei (DAPI, blue). Scale bar: 50 µm. (**F**) The heatmap shows the mean percentage of total detected spots per compartment (rows) across time (columns) for BI-TfR1 CapX, AAV-PHP.eB, and 9P31. Percentages are computed from 3 fields of view per animal across the four compartments and therefore sum to 100% within each column. (**G** and **H**) The plots show the percentage of total vascular (sum of the vascular non-nuclear and vascular nuclear) signal (**G**) and the parenchymal nuclear signal (**H**) over time. Data points represent mean with SEM; n = 6 animals per timepoint with 3 females and 3 males; unless otherwise stated (see Method details).

Compared to other capsids, CapX was rapidly cleared from sera. By 15 minutes, serum CapX levels were only ∼6% of AAV9 levels (Figure 4C), consistent with the known rapid brain accumulation and tissue uptake of TfR1-binding biologics^38,39^. Despite this rapid depletion from the serum fraction, the level of CapX in whole blood was similar to AAV9 at 15 min and higher than AAV9 at 7 and 21 days (Figure 4D). Further studies will be necessary to determine whether this observation is driven by CapX persistence in long-lived red blood cells derived from reticulocytes present at dosing, or in other long-circulating blood cell types.

We observed low variance in biodistribution across animals for CapX and 9P31, but AAV-PHP.eB was an outlier in a subset. Ranking animals by AAV-PHP.eB brain biodistribution showed 8 of 36 animals below their cohort-matched animal medians: 7.5-to 450-fold below in seven animals, and non-detectable in the eighth. In the liver, 7 of these 8 animals were 8-to 29-fold below their cohort medians (Figure S7). The deficit was specific to AAV-PHP.eB. In the same animals, the other three capsids were within ∼2.6-fold (brain) and ∼3.6-fold (liver) of their cohort medians; across all 36 animals and all four tissues, no value for CapX, 9P31 or AAV9 fell more than 6.5-fold below its cohort median. Because all four capsids were administered as a single mixture and quantified from the same samples, the deficit cannot be attributed to dosing, injection, or sample handling. These low responders were therefore excluded post hoc from AAV-PHP.eB-specific analyses.

We next looked at the location of AAV genomes at each timepoint post-dosing using RNAscope with a sense-strand probe designed to detect the AAV genome. We detected high levels of AAV-PHP.eB and CapX barcoded genomes on or within the brain vasculature by 15 minutes (Figure 4E). 9P31 was also detected, but at lower levels, whereas AAV9 was not reliably detected above background using this method (Figure S8). We next used a neural network-based tool (deepBlink)^40^ trained to convert diffraction-limited probe signal area into an estimate of the number of discrete probe spots per field of view per animal (Figure S9A-F), and then assigned each spot to one of four compartments: vascular non-nuclear, vascular nuclear, parenchymal non-nuclear, or parenchymal nuclear within each field of view analyzed per animal (Figure S9G-K). We focused the RNAscope quantification on the 15 min to 3 day timepoints, because the RNAscope signal-to-noise ratio declined at the 7 and 21 day timepoints. The low RNAscope signals at later time points are possibly due to a combination of genome clearance, which aligns with the reduction detected by dPCR bio-distribution, and/or conversion of the AAV genomes to double-stranded episomes that were less efficiently detected under the *in situ* probe hybridization conditions used in this study.

Over the course of minutes to days, the three engineered capsids diverged in how they distributed within the brain vasculature and parenchyma. The mean spots detected (spots per field of view per animal) peaked at 15 minutes post dose for CapX and at 2 hours post dose for AAV-PHP.eB and 9P31 (Figure S9G), in alignment with brain dPCR biodistribution data (Figure 4A). At 15 minutes, 88 to 94% of spots for all three capsids were located in the vascular compartments (Figure 4F-G). Between 15 minutes and 24 hours, two dramatic shifts occurred in tandem for CapX in particular: the vascular non-nuclear signal declined sharply from 65% to 7% of total spots (Figure 4F and Figure S9H), while the parenchymal-nuclear signal increased (from 2% to 44% of total spots (Figure 4H and Figure S9K). The result is a CapX-specific sequential shift in peak signal across compartments over time from vascular non-nuclear, to parenchymal non-nuclear, to parenchymal nuclear in both absolute counts (Figure S9H-K) (peaks at 15 minutes, 2 hours, and 24 hours) and fractional terms (Figure 4F) (spots in each compartment relative to total spots). By day 3, vascular nuclear spots accounted for 25% of the total remaining signal and 2% of the total signal detected at the initial 15 minute timepoint. Together, these data suggest that CapX particles rapidly transit through the vasculature and traffic to parenchymal nuclei by 24 hours.

Distinct profiles were observed for the two other engineered capsids. For both AAV-PHP.eB and 9P31 peak signal in the vascular non-nuclear compartment occurred later than for CapX, at 2 hours, and persisted longer (Figure 4F and Figure S9H). At 3 days, the vascular non-nuclear signal for AAV-PHP.eB and 9P31 accounted for 50% and 15% total signal respectively, compared to 3% for CapX (Figure 4F); the vascular-nuclear signal accounted for 34% and 47% of total signal, compared to 25% for CapX (Figure 4F). While a subset of AAV-PHP.eB particles were detected in the parenchyma at 2 hours, the persistence of a majority of particles in the vasculature through day 3 meant that we did not observe a sequential compartment peak shift as we did for CapX. 9P31 clears from the vasculature more rapidly than AAV-PHP.eB, but likewise traffics to the parenchymal nuclei more slowly and to a lesser degree than CapX.

### A Pre-clinical roadmap for the development of a gene therapy for prion disease

Advancing a gene therapy toward the clinic requires preclinical data establishing safety and efficacy before first-in-human dosing. Because an AAV therapy comprises a capsid and a transgene, we consider each in turn: first a capsid-specific safety assessment independent of any transgene, and then how the strict human specificity shapes the choice of models for the transgene-dependent efficacy and dose-finding studies needed to support clinical translation. As a preliminary and limited, non-GLP assessment of capsid-specific safety, we used CapX and AAV9 to package a CpG-depleted 4.4-kb ssAAV genome lacking a promoter or ORF, and dosed *hTFRC* KI mice at 1 × 10^13^ vg/kg (n = 3/sex/group), assessing tolerability at 3 and 21 days by hematology, organ weights, and gross and microscopic pathology. Both vectors were well tolerated, with no treatment-related effects on mortality, body or organ weights, clinical pathology, or gross and microscopic morphology relative to the PBS vehicle control (Data S1 and Table S2).

Because CapX is strictly human-specific, *hTFRC* KI animals are currently the only route to pharmacologically relevant *in vivo* biodistribution, potency, and safety data for dose-finding toward first-in-human studies. Crossing *hTFRC* KI mice to disease models or to lines carrying a humanized target gene is the most direct way to establish potency, efficacy, and safety, though it is time-consuming; alternatively, early proof-of-concept and dose range finding studies can use a well-matched surrogate capsid to assess transgene efficacy and safety that avoids the cross with *hTFRC* KI mice. Our two NIH Somatic Cell Genome Editing (SCGE)-supported prion disease programs illustrate both approaches, which we develop below.

### CapX enables efficient whole-brain stop-codon installation with a dual-vector cytosine base editor

We recently described a TadCBEd cytosine base editor that installs a premature stop codon (TGA) at Arg37 of *PRNP* in human *PRNP* transgenic mice^28^. Delivered systemically with AAV-PHP.eB, this dual-vector, split-intein editor achieved up to 44% editing and a 63% reduction in prion protein (PrP) throughout the brain. To evaluate the efficiency of base editing-mediated PrP reduction using CapX, we generated mice humanized for both *PRNP* and *TFRC* on a *Prnp*-null background (see Methods) and delivered the hSyn-driven TadCBEd R37X editor and R37X sgRNA (Figure 5A) with CapX at two doses, 2.5 × 10^12^ and 1 × 10^13^ vg/kg per AAV (5 × 10^12^ and 2 × 10^13^ vg/kg total, combined dose). Bulk installation of R37X (C7 editing) reached 23% at the low dose and 40% at the high dose (Figure 5B) and lowered brain PrP by 65% and 80%, respectively (Figure 5C). Synonymous bystander edits within the editing window were also observed (Figure S10). The greater reduction in PrP as compared to the editing frequency may reflect that while editing was measured in bulk tissue and reflects all cell CNS types, neurons are both the CNS cell type most efficiently transduced by CapX and the predominant producers of PrP^41,42^. Alternatively, this disconnect could reflect non-uniform expression from the tandem *PRNP* copies in this transgenic line^43^. These data highlight the capability of CapX to mediate efficient dual-vector delivery for whole-brain genome editing in a pre-clinical disease model.

**Figure 5.**
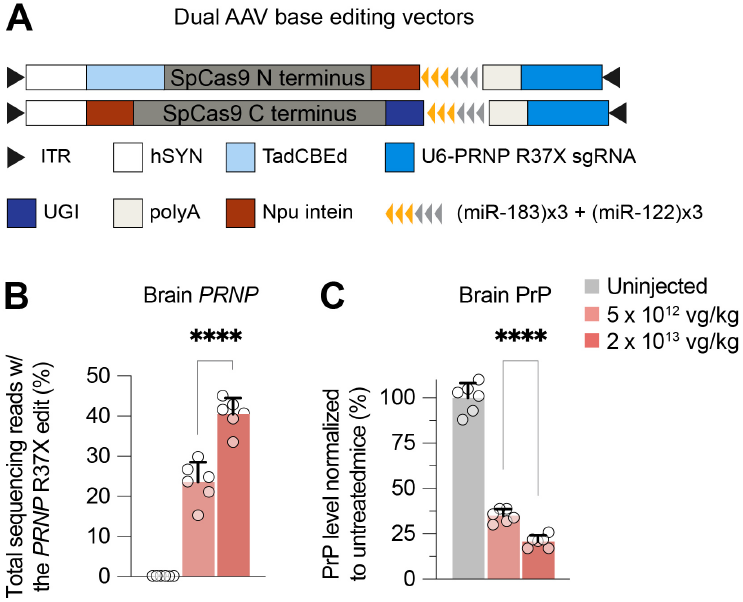
CapX enables brain-wide installation of a protein truncating mutation in *PRNP* with a dual-vector base editor. (**A**) Schematics of the split vector CBE (TadCBEd). A transgenic line containing *hTFRC* and human *PRNP* (see methods) were injected with 2.5 × 10^12^ vg/kg each AAV (5 × 10^12^ vg/kg total) or 1 × 10^13^ vg/kg of each AAV (2 × 10^13^ vg/kg total). The plots report the frequency of R37X editing (**B**) and prion protein (PrP) levels (**C**) in whole-brain hemisphere homogenates 5 weeks post-administration. Data points represent individual mice (n = 6 animals per dose) and bars show the mean with SD (unpaired two-tailed Welch’s t test; only p values < 0.05 are shown,****p < 0.0001, no statistical tests were made to uninjected animals).

### AAV-PHP.eB is less potent than CapX but can be used as a suitable surrogate in early preclinical studies

As described in our FDA INTERACT submission for a CHARM-based *PRNP* epigenetic-silencing program (Data S2)^44^, we proposed reserving *hTFRC* KI mice for GLP toxicology and on-target biodistribution while using AAV-PHP.eB as a surrogate capsid for non-GLP efficacy and proof-of-concept studies in humanized *PRNP* mice. We compared CapX and AAV-PHP.eB head-to-head. Each capsid was used to package Prnp-ZF-CHARM Kv1, which epigenetically silences mouse *Prnp*^29^, and was administered to *hTFRC* KI mice across four doses from 6.6 × 10^11^ to 1.8 × 10^13^ vg/kg. Both capsids showed similar dose-dependent accumulation in brain, spinal cord, liver, heart, and muscle, with AAV-PHP.eB reaching modestly higher brain vg/dg at the lower doses (Figure 6A). At the highest dose (1.8 × 10^13^ vg/kg), CapX achieved 96% brain-wide silencing of *Prnp* mRNA relative to untreated controls, versus 72% for AAV-PHP.eB (Figure 6B), corresponding to 84% and 63% reductions in brain PrP, respectively (Figure 6C). CapX achieved 50% silencing at 2.3x lower dose than AAV-PHP.eB for *Prnp* mRNA (absolute IC50 1.35 × 10^12^ vs 3.12 × 10^12^ vg/kg) and 3.0x lower dose for PrP protein (1.13 × 10^12^ vs 3.35 × 10^12^ vg/kg), indicating greater potency of CapX on a per-vg/kg basis. Liver *Prnp* silencing was similar for the two capsids (Figure 6D), and silencing in the heart was detectable at high doses (Figure S11). *In situ* hybridization detecting the CHARM transcript (WPRE antisense probe) and *Prnp* transcripts confirmed brain-wide expression and silencing in both treatment groups at the high dose (Figure 6E).

We next assessed silencing as a function of biodistribution to the brain. Remarkably, relative to AAV-PHP.eB, CapX achieved 50% *Prnp* mRNA and PrP lowering with ∼18-fold (absolute IC50 0.48 vs 8.45 vg/dg) and ∼20-fold (0.36 vs 7.30 vg/dg) fewer vector genomes delivered to the brain, indicating that CapX has a substantially greater per AAV particle potency (Figure 6F-G). In the liver, the two capsids had similar per particle potencies (0.42 vs 0.44 vg/ dg) for 50% *Prnp* lowering. CapX thus achieves ∼50% brain *Prnp* lowering at roughly one genome per two cells, indicating that a large fraction of the genomes it delivers to the brain reach *Prnp* expressing cells, in contrast to AAV-PHP.eB, which is less efficient at vascular transit (Figure 4), and needs an order of magnitude more genomes per cell in the brain for the same effect.

**Figure 6.**
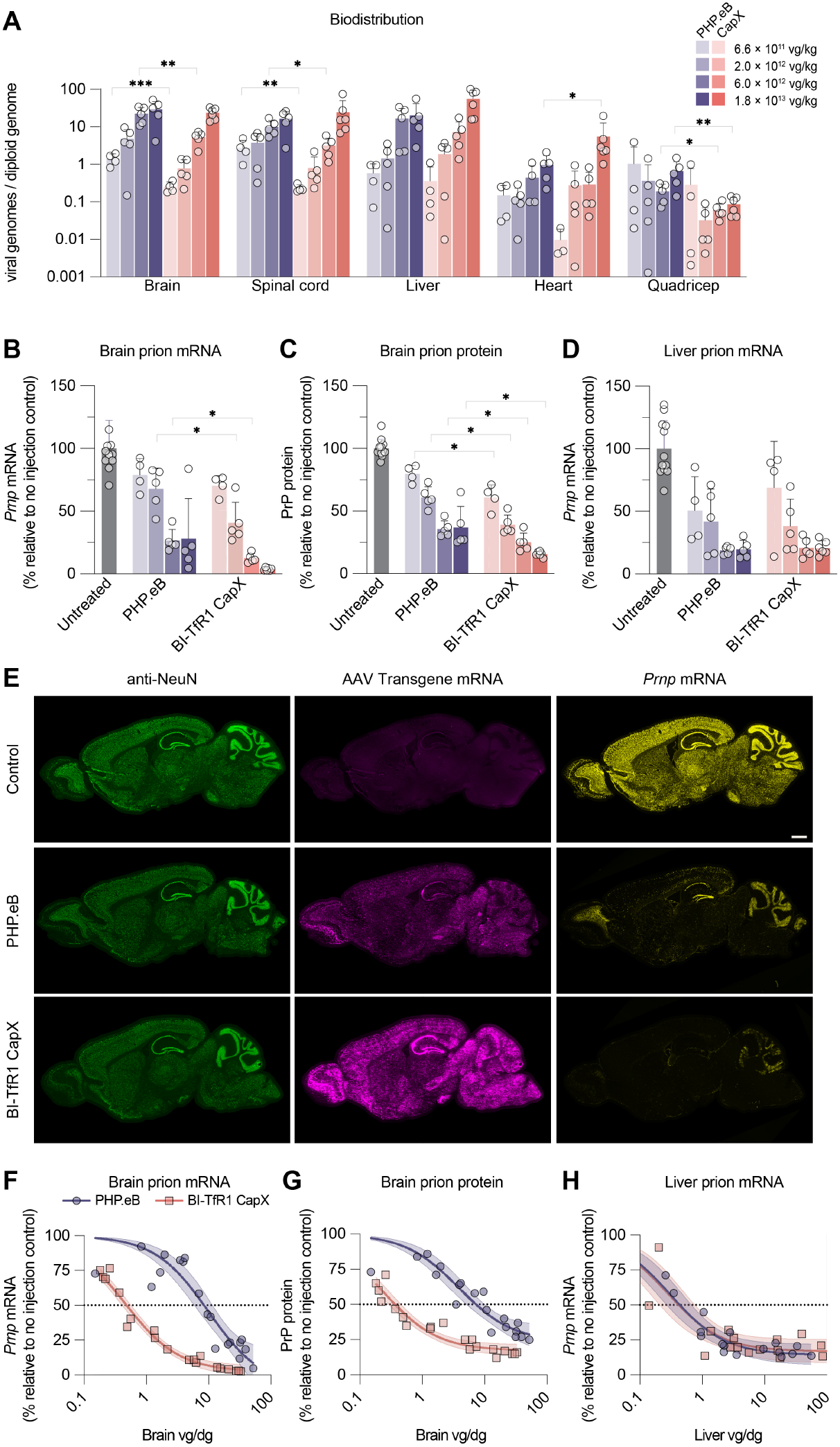
CapX:Prnp-CHARM mediates efficient epigenetic silencing of *Prnp*; AAV-PHP.eB is a suitable pre-clinical surrogate. CapX or AAV-PHP.eB were i.v. administered a mouse-*Prnp* CHARM vector across 4 different doses that range from 6.6 × 10^11^ vg/kg to 1.8 × 10^13^ vg/kg into adult *hTFRC* KI mice and tissues were collected 6 weeks later. (**A**) Vector genome biodistribution assessed by qPCR. (**B**-**D**) Mouse brain *Prnp* transcripts (**B**), brain PrP, or liver *Prnp* transcripts relative to uninjected controls. (**E**) Representative whole sagittal brain sections showing NeuN antibody staining (green) with codetection of AAV (WPRE, magenta) and *Prnp* transcripts (yellow) using RNAscope from an untreated animal or animals treated with 1.8 × 10^13^ vg/kg. (**F**-**H**) The plots show the percent *Prnp* transcripts in the brain relative to uninjected controls (**F**), PrP protein in the brain (**G**), or *Prnp* transcripts in the liver (**H**) versus biodistribution (vg/dg) in the indicated organ. Each symbol represents an individual animal (n = 6 mice per group, unless other noted in the methods). Bars show the mean with SD (**A**-**D**); curves show 95% CI as shaded (filled) bands around each fitted curve (**F**-**H**). Lognormal Welch’s t test within organ within similar doses (**A**); normal Welch’s t test within organ within similar doses (**B-D**); (*p < 0.05, **p < 0.01, ***p < 0.001, ****p < 0.0001).

### CapX can be produced and purified with similar yields as AAV9 and is compatible with commercial affinity resins

Building on these promising preclinical results we are developing a human *PRNP*-targeted CHARM epigenetic silencing gene therapy for first-in-human studies. As CapX moves toward the clinic for this and other indications, scalable operationalization of this technology will become load-bearing. To this end, we have gathered data indicating that CapX is readily manufacturable. Across multiple genomes and batches produced by triple transfection of suspension HEK293T cultures, CapX yields have proven comparable to AAV9, and post-iodixanol purification yields have routinely exceeded ∼0.5x those of AAV9 (Table S3). CapX is compatible with standard affinity resins (AAVX, AVIPure, and AAV9 resin) (Figure S12).

In support of the prion program we have generated a 50 L lot of CapX-packaged *PRNP*-CHARM at a contract drug manufacturing organization. In this run, we obtained 3 × 10^14^ vg/L at bulk lysate and a final product yield of 8.1 × 10^13^ vg/L (27% overall recovery yield) through their platform process, which included affinity chromatography with POROS AAV9, iodixanol density gradient ultracentrifugation, AEX chromatography, and tangential flow filtration (Table S4). The final product was formulated at 2.3 × 10^13^ vg/mL, with 100% of the particle volume with a predominant size between 21-23 nm (dynamic light scattering) and 85.6% full (mass photometry). Together these outcomes suggest that high-quality CapX material will be producible at clinical scale using industry standard practices.

## Discussion

From initial reprogramming of AAV9 to bind human TfR1, followed by a single round of focused maturation, we have developed CapX, a capsid that transduces the majority of neurons and a large fraction of astrocytes throughout the brain and spinal cord with a low systemic dose (2 × 10^12^ vg/kg), while reducing biodistribution in the liver and expression in several other peripheral organs relative to AAV9. While *in vitro* screening was highly sensitive at identifying BI-hTFR1 and other TfR1 binding candidates, ranking candidates across multiple metrics critical for translational use was best served by screening directly for production fitness, human brain endothelial cell transduction, and CNS transduction in humanized *TFRC* expressing mice.

We find that CapX behavior reflects the native transporter activity of its binding partner, TfR. Its performance is most clearly distinguished from capsids that co-opt non-transporter proteins by the greater speed and completeness of its BBB transit. Within minutes, CapX achieves peak brain vasculature association; within 24 hours, the majority of detected particles have moved into the parenchyma, with a large fraction localized to nuclei. In contrast, AAV-PHP.eB and 9P31, which engage the GPI-anchored proteins LY6A and Car4, respectively, remain largely vasculature-associated for days. This rapid, efficient transit likely underlies the exceptional per-genome potency of CapX, which may also be enhanced by direct engagement with neuronally expressed TfR. Strikingly, when paired with the epigenetic editor *Prnp*-CHARM, the capsid achieves roughly 50% brain-wide *Prnp* mRNA lowering at approximately one vector genome per two cells, requiring 18-fold fewer genomes per brain cell than AAV-PHP.eB.

Consistent with its rapid tissue uptake, we find that CapX is largely depleted from serum within minutes. The divergence between serum clearance and prolonged whole-blood persistence suggests that CapX is rapidly cleared from free plasma circulation, but that a fraction of the administered dose may remain blood cell-associated. Further studies are needed to determine whether this blood distribution persists at later time points and whether it relates to binding or uptake into reticulocytes or other circulating cells that express TfR1. Notably, no hematologic changes were observed in our limited scope, capsid-focused tolerability study (Table S2).

CapX binding depends on the human-unique Tyr247 residue of TfR1 and is thereby strictly human-specific. As such, it presents translational pros and cons distinct from other capsids, including the ALPL-binding capsid family, that mediate BBB crossing in rodents and primates but do not leverage the use of a clinically validated transport protein. The human TfR1 specificity of CapX places significant weight on humanized models throughout preclinical development, as animals expressing human *TFRC* are the only pharmacologically relevant models for biodistribution, potency, and safety studies. Several mouse models with different regions of *TFRC* humanized (extracellular or apical domain only) are available and respond comparably to CapX in terms of tropism and CNS transduction efficiency; note that because TfR1 homodimerizes, these mice must be used as homozygotes. To test editors or assess disease models these mice can be crossed to lines bearing other humanized genes or variants. Alternatively AAV-PHP.eB can be employed as a surrogate for initial proof-of-concept studies and, where appropriate, non-GLP efficacy studies. Our comparison of CapX and AAV-PHP.eB across a range of doses (Figure 6) aim to support these efforts or guide program-specific bridging, as necessary.

Our results from human TfR1 protein, human receptor studies, and *hTFRC* KI mice support the ability of CapX to bind human TfR1 and efficiently cross the human BBB to reach neurons and glia throughout the brain. But as we consider the clinical potential of this capsid, we acknowledge that which may not be predictable. The first test of an engineered BBB-crossing capsid changed the AAV safety landscape. An investigational gene therapy for STXBP1-related developmental and epileptic encephalopathy was associated with fatal rapid-onset cerebral edema in the first treated patient. This adverse event had not previously been observed with AAV gene therapy and was not predicted from the program’s non-human primate toxicology study^45^. Information about the underlying cause remains limited, although the identification of the metalloprotease ADAM15 as the capsid’s receptor was recently made public^45^. This tragic event brings to the forefront the fact that redirecting AAV tropism toward the CNS can shift the risk profile of systemic gene therapy from more precedented liver and cardiac toxicities toward new events such as cerebral edema. Whether this event reflects a toxicity specific to engaging ADAM15; to the capsid, which was optimized for crossing the BBB by engaging the NHP receptor; to transgene overexpression in the endothelium^46^; or the disease, rather than a broader concern, is not yet known. No data has been reported to support or rule out the possibility that this may represent a class effect that would extend to all engineered BBB-crossing AAVs or only to the subset that dwell longer on or in the endothelium due to their binding of cell surface proteins not known to mediate transcytosis.

It is critical that we develop a more detailed mechanistic understanding of how each capsid’s route of CNS entry shapes risk. Our kinetic and vascular-accumulation studies are a first step, and we will continue to define the BBB-crossing characteristics of CapX and other BBB-crossing capsids. We acknowledge that safety and efficacy in humans can only be established through cautious clinical testing, starting with low doses anticipated to be effective that maximize safety margins in patients with severe, debilitating or life-threatening disease for whom the potential benefit justifies the risk. While the outcome of next steps cannot be known, the potential of this capsid motivates continued, circumspect advancement. Ultimately, thoughtful clinical studies must balance and contextualize the risks of new technologies while also not allowing their promise to go unexplored.

## Methods

### Animals

All procedures were performed as approved by the Broad Institute or MIT Division of Comparative Medicine IACUC. The following animals were used in this work: *hTFRC* KI mice with a fully humanized extracellular domain (Biocytogen, C57BL/6-Tfr1tm1TFR1/Bcgen, B-hTFR1, strain#: 110861), humanized apical domain knock-in (hAPI) KI mice^33^ (JAX, C57BL/6-Tfrctm1(*TFRC*)Bdes/ J, strain#: 038212), immunodeficient NOD *scid* gamma (NSG) mice from (JAX, strain #, 005557) and the Prnp^ZH3^; Tg(PRNP)26372 double mutant mouse line which harbors mouse *Prnp* knock-out as well as full-length human *PRNP* (JAX, strain, #075939)^47,48^. Mice were randomly assigned to groups based on predetermined sample sizes to achieve similar sex and age distribution. Experimenters were not blinded to the sample groups. AAVs were administered intravenously (i.v.) via the retro-orbital sinus. Mice were euthanized with either cervical dislocation, decapitation or transcardial perfusion under isoflurane anesthesia or CO_2_ inhalation.

### Plasmids

Capsid library assembly was performed as previously described^49^. Briefly, we designed an oligo pool comprising single- and dual-site saturated mutagenesis (SSM and DSSM) to the BI-hTFR1^24^ sequence or surrounding AAV9 sequence and reference sequences and had the pool printed by Twist Bioscience. 5 pM of the oligo pool was used as an initial reverse primer along with 0.5 µM Assembly-XbaI-F oligo as the forward primer to amplify and extend 10 ng pUC19-AAV9-347-585 for 10 cycles. Then, the reaction was run with 0.5 µM of primer Assembly-AgeI-R and amplified for an additional 25 cycles (Table S5).

The PCR product was then inserted into an hSYN-mRNA or CMV-CBA mRNA selection backbone via NEBuilder HiFi DNA Assembly Master Mix (NEB, E2621L) at a 3:1 Molar ratio of insert:vector in a 80 µL reaction volume incubated at 50 °C for 1 h, followed by incubation at 72 °C for 5 minutes. Afterwards, 4 µL of Quick CIP (NEB, M0508S) was added to the reaction and incubated at 37 °C for 30 mins to dephosphorylate unincorporated dNTPs that may inhibit T5 Exonuclease. Finally, 4 µL of T5 Exonuclease (NEB M0663S) was added to the reaction mixture and incubated at 37 °C for 30 mins to remove unassembled products. The final assembled products were cleaned using AM-Pure XP beads (Beckman, A63881) following the manufacturer’s protocol and the final product concentrations were quantified with a Qubit dsDNA HS Assay Kit (Thermo Fisher Scientific, Q32851) using a Qubit fluorometer. The final purified and quantified product was used for AAV library production (see AAV production section). Note all capsid sequences within the library, including AAV9 and 9P31 incorporate a K449R mutation introduced to facilitate cloning^3^.

The following plasmids were synthesized by GenScript: an AAV transgene plasmid lacking CpGs and ORFs outside of the ITRs; the dual reporter vectors ssAAV-CAG-mScarlet-P2A-luc-WPRE and ssAAV-EFS-mScarlet-P2A-luc-WPRE; Rep-Cap plasmids for 9P31 and CapX; and four barcoded transgene plasmids (Barcodes 03, 08, 09, and 13). The Rep-Cap plasmids CapX, BI-hTFR1, and AAV-PHP.eB contain a K449R substitution (AAV9, VP1 numbering). AAV9 and 9P31 were generated without the K449R mutation. For the barcoded plasmids, a unique 300 bp barcode was inserted between the sfGFP and WPRE sequences of a CAG-H2B-sfGFP-WPRE-hGH-U6 backbone. Finally, a series of plasmids was synthesized for the ectopic expression of receptors *in vitro* in CHO cells and *in vivo* in NSG animals using the two-step assay. These transgenes, encoding human TfR1, rhesus macaque TfR1, mouse Ly6A, or chimeric human/rhesus TfR1 variants, were cloned into an ITR-flanked backbone containing a CAG promoter, three tandem miR122 binding sites, a shortened WPRE, and an hGH polyA signal. All plasmid synthesized plasmid maps will be available at Addgene upon publication.

The Rep-Cap AAV9, BI-hTFR1 (Addgene, 218796), ssAAV-CAG-NLS-mScarlet-P2A-luc-WPRE (Addgene, 218793) and Human TfR1 C-terminal Fc-fusion plasmids were described^24^. The mouse-*Prnp* CHARM vector (Addgene, 220842) was previously described^29^.

### AAV production

For library AAV production, HEK293T/17 cells (ATCC, CRL-11268) were seeded at 2.2 × 10^7^ cells per 15 cm dish the day before transfection and maintained in DMEM with GlutaMAX (Gibco, 10569010) supplemented with 5% FBS and 1x non-essential amino acids (NEAA; Gibco, 11140050). Each plate was triple-transfected with 39.93 µg of pHelper, RepStop (encoding the AAV2 Rep genes), and pUC19 at a 2:1:1 mass ratio together with 10 ng of assembled library DNA using PEI Max at (Polysciences, 24765). Medium was exchanged for fresh DMEM with 5% FBS and 1x NEAA at 20 h post-transfection. The medium and cell lysate were harvested at 60 h post-transfection and purified as previously described^49^.

For individual recombinant AAVs, HEK293T/17 cells were adapted for growth in suspension in FreeStyle F17 medium (Thermo Fisher Scientific, A1383501) at 37 °C, 8% CO2, and 125 rpm. Cells were seeded at approximately 1 × 10^6^ cells/mL. 24 h later approximately 2 × 10^6^ cells/mL cells were transfected with pHelper, Rep-Cap, and transgene plasmids at a 2:1:1 mass ratio (2 µg total DNA per 106 cells) using Transporter 5 transfection reagent (Polysciences, 26008) at a 2:1 PEI:DNA mass ratio. Three days post-transfection, cells were pelleted at 300-2,000 x *g* for 15 min in 250 mL conical bottles, the supernatant discarded, and pellets stored at −20 °C until purification. Each pellet, corresponding to 200 mL of culture, was resuspended in 7 mL of 500 mM NaCl, 40 mM Tris base, and 10 mM MgCl_2_ containing 100 U/mL Salt Active Nuclease (ArcticZymes, 70920-202) and incubated at 37 °C for 1.5-2 h. Lysate was clarified at 4,000 x *g* for 30 min and loaded onto a discontinuous iodixanol gradient prepared from OptiPrep (Cosmo Bio, AXS-1114542) at 60%, 40%, 25%, and 15% (5, 5, 6, and 6 mL, respectively) in OptiSeal tubes (Beckman Coulter, 361625). Gradients were centrifuged in a Beckman Type 70 Ti rotor at 340,252 x *g* for 1 to 2.25 h at 18 °C. Approximately 4.5 mL was drawn from the 40-60% interface with a 16-gauge needle, filtered through a 0.22 µm PES filter, buffer-exchanged into PBS containing 0.001% Pluronic F-68 using 100K MWCO protein concentrators (Thermo Fisher Scientific, 88532), and concentrated to 200-500 µL. The concentrated vector was filtered through a 0.22 µm PES filter, aliquoted, and stored at −80 °C.

### AAV titering

To determine AAV titers or assess library variant production fitness, 5 µL of each purified AAV library along with a known internally generated AAV standard and a non-template control sample were incubated in technical duplicates with 100 µL of an endonuclease cocktail consisting of 1000U/mL Turbonuclease (Sigma T4330-50KU) with 1x DNase I reaction buffer (NEB B0303S) in UltraPure DNase/ RNase-Free distilled water at 37 °C for 1 h. Next, the endonuclease solution was inactivated by adding 5 µL of 0.5M, pH 8.0 EDTA (Thermo Fisher Scientific, 15575020) and incubated at room temperature for 5 minutes and then at 70 °C for 10 minutes. To release encapsidated AAV genomes, 120 µL of a Proteinase K cocktail consisting of 1M NaCl, 1% N-lauroylsarcosine, 100 µg/mL Proteinase K (Qiagen, 19131) in UltraPure DNase/RNase-Free distilled water was added to the mixture and incubated at 56 °C for 2 to 16 hours. The Proteinase K treated samples were then heat inactivated at 95 °C for 10 minutes. The released AAV genomes were then serially diluted between 460-460,000x in dilution buffer consisting of 1x PCR Buffer (Thermo Fisher Scientific, N8080129), 2 µg/mL sheared salmon sperm DNA (Thermo Fisher Scientific, AM9680), and 0.05% Pluronic F68 (Thermo Fisher Scientific, 24040032) in UltraPure Water (Thermo Fisher Scientific, 10977015). Unless stated otherwise, two sets of primer probes were used (ITR_F/R/Probe) and (WPRE_F/R/Probe) (Table S5) to quantify viral genomes either through ddPCR or dPCR.

For titering via ddPCR, following sample dilution, 2 µL of the diluted samples were used as input in a ddPCR supermix for probes (Bio-Rad, 1863023) following the manufacturer’s protocol. Droplets were generated using a QX100 Droplet Generator following the manufacturer’s protocol. The droplets were then transferred to thermocycler and cycled according to the manufacturer’s protocol with an annealing/extension of 58°C for 1 minute. Finally, droplets were read on a QX100 Droplet Digital System to determine titers.

### AAV administration

#### Library screens

The BI-hTFR1 mutagenesis library was screened in *hTFRC* KI mice. Three adult female mice 7-8 weeks old were each given 2.0 × 10^11^ vg by i.v. injection and tissues were harvested and fresh frozen 3 weeks post injection without transcardial perfusion.

#### In *vivo* ectopic receptor expression assay

A “two-step” *in vivo* receptor expression assay was performed to screen the BI-hTFR1 mutagenesis library in NSG mice ectopically expressing defined receptors in the brain vasculature. In the first step, AAV-BI30^30^ was used to deliver human *TFRC*, rhesus macaque *TFRC, or* mouse *Ly6a* expression under the CAG promoter in brain vasculature at 1 × 10^11^ vg/ mouse i.v. (n = 3 females/group). In the second step, three weeks later, the BI-hTFR1 mutagenesis library was administered i.v. at 2 × 10^11^ vg/mouse. After an additional 3 weeks tissues were collected and fresh frozen without transcardial perfusion.

#### Reporter expression studies

CapX, BI-hTFR1, and AAV9 were each packaged with a dual reporter transgene (ssCAG-NLS-mScarlet-P2A-luc-WPRE-pA) and separately administered i.v. to *hTFRC* KI mice at 2.0 × 10^12^ vg/kg (n = 3/sex/group). Tissues were harvested 3 weeks post-injection following transcardial perfusion with ice-cold PBS. PFA perfusion was omitted so that tissues remained suitable for immunohistochemistry (IHC), biodistribution, and luciferase-based transduction analysis. Portions of the brain and spinal cord were drop-fixed in 4% PFA for IHC. All other tissues were frozen and stored for analysis.

#### For cryo-fluorescence tomography (CFT)

mice were injected i.v. at 2 × 10^12^ vg/kg with either AAV9 packaging ssCAG-mScarlet-P2A-luc-WPRE-pA, CapX packaging the same CAG transgene, or CapX packaging ssEFS-mScarlet-P2A-luc-WPRE-pA (n = 2/sex/group). Two uninjected animals were also processed and imaged to judge tissue and diet-based autofluorescence. Tissues were harvested 3 weeks post-injection. Animals were placed on an irradiated Teklad global 14% rodent diet (Inotiv, 2914) 10 days prior to euthanasia to reduce autofluorescence for CFT imaging. CFT cohort animals were then euthanized with transcardial perfusion under isoflurane anesthesia with ice-cold PBS and the thoracic cavity was sutured close. The whole bodies were then submerged into isopentane in dry ice for 15 minutes and stored in −80 °C. Samples were then packed and shipped to EMIT Imaging to perform CFT.

#### Transduction comparison of CapX in *hTFRC* KI and hAPI KI mice

a ssCAG-NLS-mScarlet-P2A-Luciferase-WPRE-pA packaged reporter that was delivered at 2 × 10^12^ vg/kg. Three weeks later, tissues were harvested without perfusion.

#### To assess tolerability of AAV9 or CapX using histopathology

*hTFRC* KI mice were injected with a CpG-depleted 4.4-kb ssAAV genome lacking a promoter or ORF at 1 × 10^13^ vg/kg and were terminally assessed at 3 or 21 days (n = 3/sex/group) post administration. The AAV was titered using ITR and CpG_depleted primer and probe sets (Table S5). Tissues were collected without perfusion and reviewed under a board-certified comparative pathologist at MIT Division of Comparative Medicine.

#### Pooled barcoded biodistribution and BBB crossing study

To assess AAV9, AAV-PHP.eB, 9P31, and CapX together within the same animal, an AAV pool containing the capsids were i.v. administered to adult *hTFRC* KI mice at a total dose of 8.0 × 10^12^ vg/kg total (2.0 × 10^12^ vg/kg for each capsid). Mice were returned to their home cages. Blood and tissues were collected at 15 min, 2 h, 24 h, 3 d, 7 d, or 21 d post-injection, (n = 3/sex/group). Under isoflurane anesthesia, whole blood was collected via retro-orbital sinus using blood collection capillaries (Fisherbrand, 22-362566) and separated into lithium heparin or serum separator tubes (BD Diagnostic Systems, Cat# 365965 and 365967, respectively). Mice were then perfused with ice-cold PBS at 2 mL/min for 4 min to clear blood from tissues. Tissues were collected for ISH and dPCR (see below for details).

#### Dual base editing

To evaluate dual base editors designed to install a stop codon in human *PRNP*, we crossed Tg26372 mice, which carry the full human *PRNP* gene^48^ and its noncoding sequence on a mouse *Prnp* (−/-) background^47^ to *hTFRC* KI mice until animals were homozygous for both humanized alleles and null for both mouse *Prnp* alleles. Dual AAV editors were delivered by CapX i.v. at either 2.5 × 10^12^ vg/kg per AAV or 1 × 10^13^ vg/kg per AAV, (n = 6 animals/dose). Tissues were harvested for NGS and ELISA at 5 weeks post injection (see below section for details).

#### PRNP-ZF-CHARM

The mouse-*Prnp* targeting CHARM Kv1 vector (Addgene, Plasmid #220842) was packaged into CapX or AAV-PHP.eB and administered i.v. at either 6.6 × 10^11^, 2.0 × 10^12^, 6.0 × 10^12^ or 1.8 × 10^13^ vg/kg in *hTFRC* KI animals (n = 3/ sex/group). A second cohort of animals (n =3/sex/ group) were i.v. injected at the 1.8 × 10^13^ vg/kg dose in order to collect brains for ISH. Animals were harvested at 6 weeks post-administration without perfusion and tissues were collected and frozen for DNA, RNA, and protein or ISH analysis.

### Fc pull-down assay

Fc construct DNA was transfected into HEK293T/17 cells (ATCC, CRL-11268) with PEI (Polysciences, 23966) at 40 µg DNA per 15 cm dish in DMEM containing 5% FBS. At 12-16 h post-transfection, cells were rinsed with PBS and the medium replaced with serum-free medium (Lonza, BEBP12-764Q). Conditioned medium containing secreted Fc-fusion protein was collected 48 and 96 h after the medium change and pooled, filtered through a 0.22 µm filter (Millipore, SE1M003M00), and stored at 4 °C. To purify the Fc-fusion proteins, protein A-conjugated magnetic beads (35 µL; Thermo Fisher Scientific, 10001D) and Tween-20 (0.05% final) were added to 30 mL of conditioned medium and incubated overnight at 4 °C with end-over-end rotation. Beads were then washed three times in DPBS with 0.05% Tween-20. Expression was assessed by running a 5 µL aliquot of protein-bound beads on a 4-12% polyacrylamide gel; the remainder was carried into the pull-down assay bead-bound, without elution.

Fc-fusion protein-bound beads (10 µL) were mixed with 1 × 10^10^ vg of the AAV capsid library in DPBS containing 0.05% Tween-20 and 1% BSA and incubated overnight at 4 °C. AAV-bound beads were washed three times in PBS with 0.05% Tween-20, then treated with proteinase K to release encapsidated genomes. Released genomes were purified with AMPure XP beads (Beckman Coulter, A63881) per the manufacturer’s protocol and carried into PCR recovery and NGS library preparation as described in the NGS section.

### NGS assessment of mutagenesis library screen

Library screening with the BI-hTFR1 mutagenesis library in hCMEC/D3 binding and transductions were performed. For binding, cells were seeded at 30,000 cells per well, at 24 hours post-seeding, medium was exchanged for fresh cold media that contained the library. The plate was then maintained at 4 °C with gentle rocking for an hour, followed by five PBS washes. Total DNA was extracted using a DNeasy Kit (Qiagen, 69504). For transduction, cells were seeded at 10,000 per well, at 24 hours post-seeding media was exchanged with fresh medium that contained the library. Cells were incubated for an additional 24 hours and total RNA was recovered using an RNeasy Kit (Qiagen, 74104).

Viral DNA was recovered from Fc screens by treating samples in a Proteinase K cocktail consisting of 1M NaCl, 1% N-lauroylsarcosine, 100 µg/mL Proteinase K (Qiagen, 19131) in UltraPure DNase/RNase-Free distilled water and incubated at 56°C for 2 to 16 hours. The Proteinase K treated samples were then heat inactivated at 95 °C for 10 minutes.

Total RNA was recovered from tissues with TRIzol (Invitrogen, 15596026) according to the manufacturer’s instructions and further cleaned up using an RNeasy kit (Qiagen, 74106) with on-column DNA digestion. RNA was converted to cDNA with the Maxima H Minus Reverse Transcriptase (Thermo Fisher Scientific, EP0751) kit according to the manufacturer’s protocol, using an anchored Oligo-dT(20) primer (Table S5).

Libraries were then prepared for sequencing as previously described^18^. Briefly, qPCR was first performed on extracted AAV genomes or transcripts to determine sample-specific cycle thresholds and prevent overamplification. A first-round PCR using equal primer pairs (AAV9_588_NGS, primers 1-8; Table S5) and Q5 Hot Start High-Fidelity 2X Master Mix (NEB, M0494L) was used to recover library-specific sequences and attach partial Illumina Read 1 and Read 2 sequences, with annealing at 65 °C for 20 s and a 60 s extension. PCR products were purified with AMPure XP beads per the manufacturer’s protocol, eluted in 25 µL UltraPure water (Thermo Fisher Scientific, 10977015), and 2 µL was used as input for a second-round PCR that added unique Illumina adaptors and dual index primers (NEB, E7600S) over 5-7 cycles under the same Q5, annealing, and extension conditions. Second-round PCR products were bead-purified and eluted in 25 µL UltraPure water, then quantified and sized using an Agilent High Sensitivity DNA Kit (Agilent, 5067-4626) on an Agilent 2100 Bioanalyzer. Libraries were pooled, diluted to 2-4 nM in 10 mM Tris-HCl, pH 8.5, and sequenced per the manufacturer’s instructions on an Illumina NextSeq 550 using a NextSeq 500/550 Mid or High Output Kit (Illumina, 20024904 or 20024907), or on an Illumina NextSeq 1000 using a NextSeq P2 v3 kit (Illumina, 20046812). Reads were allocated as I1: 8, I2: 8, R1: 150-200, R2: 0. NGS screening data was viewed using a custom built, open-source drag and drop plotting, data selection, and filtering software (https://github.com/vector-engineering/plotplot).

### NGS assessment of base editing

Whole brain hemispheres were pulverized and stored as 10 mg aliquots at −80 °C. Tissue was lysed in 500 µL QuickExtract DNA Extraction Solution (Lucigen, QE09050) for 1 h at 55 °C with shaking at 800 rpm, homogenized manually by pipetting, and incubated for a further 5 min. 100 µL of lysate was transferred to a heat block and incubated at 65 °C for 6 min followed by 98 °C for 10 min. For NGS library preparation, the *PRNP* locus was amplified with primers PRNP_R37_F/R (Table S5) and NEB-Next Ultra II Q5 Master Mix (NEB, M0544X) using the following thermocycling protocol: 98 °C for 30 s; 24 cycles of 98 °C for 10 s, 68 °C for 20 s, and 72 °C for 30 s; and a final extension at 72 °C for 5 min. Unique Illumina-compatible barcodes were then attached using the following protocol: 98 °C for 30 s; 8 cycles of 98 °C for 10 s, 64 °C for 20 s, and 72 °C for 30 s; and a final extension at 72 °C for 30 s. Libraries were pooled and purified with AMPure XP beads (Beckman Coulter, A63880) at a 0.9x ratio.

Sequencing was performed on an Illumina MiSeq according to the manufacturer’s instructions using a MiSeq v2 300 kit (Illumina, MS-102-2002). Reads were demultiplexed with MiSeq Reporter software v2.6 (Illumina) and analyzed with CRISPResso2 (v2.3.3), with the minimum average read quality set to 30, ‘discard_indel_reads’ set to TRUE, and the quantification window set to 20. Indel percentages were quantified as (reads discarded)/(reads aligned to all amplicons) x 100, and base editing percentages as (indel-free reads containing the specified edit)/ (reads aligned to all amplicons) x 100.

### Immunohistochemistry (IHC)

Hemispheres from brains were sectioned using a vibratome to 80 µm. Spinal cords were sectioned using a cryostat to 14 µm and mounted on slides. Brain IHC was performed on floating sections. Tissues were incubated with antibodies diluted in PBS containing 5% donkey serum, 0.1% Triton X-100, and 0.05% sodium azide. Primary antibodies to NeuN 1:500 (Invitrogen, MA5-33103) and SOX9 1:250 (abcam, ab185966) were incubated at room temperature overnight. The sections were then washed and stained with secondary Alexa-conjugated antibodies Alexa Fluor Plus 488 (Invitrogen, A32723), or Alexa Fluor 647 (Invitrogen, A-31573) at 1:500-1:1000 for four hours or overnight (brain), or 2 hours (spinal cord). The sections were incubated for 10 minutes with DAPI (BioLegend, 422801, 1:1000 dilution) for nucleic acid staining. All sections were mounted in ProLong Diamond Antifade Mountant with DAPI (Invitrogen, P36971).

### *In situ* hybridization (ISH)

Fresh-frozen whole brain hemispheres were cryosectioned to 14 µm and mounted on slides. Sections were fixed in 4% PFA for 30 min at 4 °C, dehydrated through a graded ethanol series (50%, 70%, 100%; 5 min each), and treated with RNAscope Hydrogen Peroxide for 10 min to quench endogenous peroxidase activity. Target retrieval was performed in 90 °C deionized water for 10 s, followed by 1x Target Retrieval Buffer (ACD, 322001) for 8 min at 90 °C, then RNAscope PretreatPro (ACD, 323770) for 30 min at 40 °C. Hybridization and signal development was performed with the RNAscope Multiplex Fluorescent Reagent Kit v2 (ACD, 323110) according to the manufacturer’s protocol (ACD, Doc. No. 323100-USM) using the probes and fluorophores described below.

#### Pooled *in vivo* kinetics barcode detection

Custom probes against the sense strand of each unique AAV transgene barcode were ordered from ACD: Barcode 03 (1815111-C1), Barcode 08 (1816351-C1), Barcode 09 (1814771-C1), Barcode 13 (1814781-C1), and WPRE (1690601-C3). Signal was developed with TSA Vivid 570 (ACD, 323272) for C1 probes and TSA Vivid 650 (ACD, 323273) for the C3 probe, both at 1:2000. Slides were mounted with ProLong Diamond Antifade Mountant with DAPI (Invitrogen, P36971).

Specificity of these custom barcode probes were confirmed in HEK293T/17 cells (ATCC, CRL-11268). Cells were seeded at 1 × 106 cells/well and transfected 24 h later with the corresponding barcode plasmid using TransIT-LT1 (Mirus Bio, MIR2306) per the manufacturer’s instructions. After a further 24 h at 37 °C and 5% CO_2_, cells were prepared as described in the ACD technical note (Doc. No. MK-50-010) and assayed as above, with each probe tested against both its matched and non-matched barcodes (Figure S6).

#### AAV transgene and *Prnp* mRNA detection

Sections were hybridized with a cocktail of WPRE-C1 (ACD, 410051) and Mm-*Prnp*-C2 (ACD, 476611-C2) to detect AAV vector transcripts and mouse *Prnp* transcripts, respectively. The signal was developed with HRP-C1 and TSA Vivid 650 (1:2000; ACD, 323273) for WPRE, and HRP-C2 with TSA Vivid 570 (1:2000; ACD, 323272) for *Prnp*.

A subset of these sections were processed for immunohistochemistry after the RNAscope assay to co-visualize neuronal markers. Sections were blocked in 1.5% normal donkey serum (Sigma, 566460-5ML) with 0.3% Triton X-100 (Thermo Fisher Scientific, 85111) for 30 min at room temperature and incubated overnight at 4 °C with recombinant monoclonal anti-NeuN (Invitrogen, 702022; 5 µg/mL). After washes in 1x PBS, sections were incubated with Alexa Fluor Plus 488 secondary antibody (Invitrogen, A32731; 1:500) for 2 h at room temperature. Nuclei were stained with DAPI (BioLegend, 422801; 1:1000) for 10 min, and sections were washed and mounted with ProLong Diamond Antifade Mountant (Invitrogen, P36971).

### Pooled AAV *in vivo* kinetic study

Flash-frozen brain hemispheres were cryopulverized using a Covaris CP02 cryoPREP Automated Dry Pulverizer. DNA was isolated from cryopulverized brain and flash-frozen liver using the DNeasy 96 Blood & Tissue Kit (Qiagen, Cat# 69582) according to the manufacturer’s instructions. Serum and whole blood were digested with thermolabile proteinase K (New England Biolabs, Cat# P8111S) prior to analysis to release encapsidated genomes. To resolve concatemerized AAV genomes, samples were digested with EcoRV-HF and SphI-HF (New England Biolabs, Cat# R3195L and R3182L) per the manufacturer’s instructions and heat inactivated.

dPCR was used to quantify viral genomes in tissues using a QIAcuity Eight Platform System. Tissue DNA was diluted 1:16 to 1:16,000 and blood and serum samples were diluted 1:84 to 1:84,000 so that target concentrations were within the working range of the QIAcuity 8.5k 96-well Nanoplate (Qiagen, 250011). dPCR was performed on the diluted samples following the manufacturer’s protocol in the QIAcuity Probe PCR Kit (Qiagen, 250103). Customized primers and multiplexing probes targeting barcodes and *Gapdh* (Table S5) were used. The dPCR was performed with conditions of 95 °C incubation (2 minutes for brain DNA, and 5 minutes for blood and serum sample), followed by 50 cycles of 95 °C for 15 secs and combined 60 °C for 1 min. Viral genomes in brain and liver tissues were quantified based on the positive counts of barcoded genomes packaged into CapX, AAV-PHP.eB, 9P31, or AAV9, and normalized to *Gapdh* gene copy numbers. Blood and serum were quantified as per µL of undiluted sample input. Following sample dilution, 2 µL of diluted sample was combined with 14 µL of master mix (Qiagen, 250101) following the manufacturer’s protocol. Samples were transferred to each well of a 8.5K 96 well QIAcuity Nanoplate (Qiagen, 250021). The nanoplate was sealed according to the manufacturer’s instructions and analyzed using a QIAcuity digital PCR system.

To visualize the unique AAV transgene barcodes *in situ*, whole brain hemispheres were cryosectioned to 14 µm. ISH was performed as described above (see Pooled *in vivo* kinetics: barcode detection). Capsids were detected in pairs, one per fluorescence channel of the same image: CapX (barcode 09, TSA Vivid 570) was paired with AAV-PHP.eB (barcode 03, TSA Vivid 650) and AAV9 (barcode 08, TSA Vivid 570) was paired 9P31 (barcode 13, TSA Vivid 650). After ISH, tissues were stained with a lectin conjugated to FITC (Invitrogen, L32478) in order to mark the vasculature. Tissue samples were then mounted with ProLong Diamond Antifade with DAPI (Invitrogen, P36971). Samples were then imaged on a Plan-Apochromat 63x/1.4 Oil DIC M27 (Zeiss, 420782-9900-799) on a Zeiss LSM 900 confocal. DAPI, FITC (lectin), TSA Vivid 570 and TSA Vivid 650 were imaged across three brain regions corresponding to the cortex, thalamus, and striatum per animal in 16-bit and with a z thickness of 5 µm and then collapsed down to MIPs.

Segmentation was performed on the image for vessels and nuclei to generate masks. For vessel segmentation scikit-image 0.25.0 (see code for detail algorithm and parameters) was used. Nuclear segmentation was performed via Cellpose 3.1.0 using the nuclear model. Nuclear masks ≥ 50% inside a vessel mask were considered vascular nuclei.

To resolve the diffraction-limited ISH probe signal into discrete puncta, we used deepBlink^40^ v0.1.4 with the pre-trained smfish1 model to the raw 16-bit images, which returns sub-pixel spot centroids. Because deepBlink reports no intensity value, each detected spot was assigned a brightness equal to the maximum pixel value within a 3-px-radius disk centered on its rounded coordinates in the raw channel. deepBlink also returns puncta in background-only images collected from uninjected animals, so an intensity cutoff was calibrated separately for each bar-code-probe and the cutoff set to the 99th percentile of the resulting intensity distribution, excluding 99% of background detections. Each cutoff was then applied unchanged to all injected samples and time-points for that barcode-probe. Retained spots were classified into one of the mutually exclusive compartments: vascular non-nuclear, vascular nuclear, parenchymal non-nuclear and parenchymal nuclear. Three fields of view were acquired per animal and averaged to yield a single value per animal; data are reported as the mean number of thresholded spots per fields of view.

### Microscopy

Images of native mScarlet or *in situ* hybridization of the whole brain or spinal cord were taken on a Keyence BZ-X810. Single plane tiled images were taken with autofocus mode on and stitched using the Keyence Analysis Software. For subregion images of the brain - cortex, striatum, hippocampus, thalamus, deep cerebellar nuclei, or cells from *in vitro* assays were taken on a Zeiss LSM 900 confocal using either a 20x air objective Plan-Apochromat 20x/0.8 Air M27 (Zeiss, 420650-9902-000) or a Plan-Apochromat 63x/1.4 Oil DIC M27 (Zeiss, 420782-9900-799). Within each experiment, images were taken with predetermined, optimized, and fixed exposure times, percent laser power and/or gain to allow image comparisons.

### Cryo-Fluorescence Tomography

Images were collected and analyzed by EMIT imaging. Fluorescence biodistribution across CNS and peripheral organs was quantified using ROI analysis of paired RGB white light and 50 ms mScarlet fluorescence images. ROIs were drawn on the white light image and fluorescence measured at corresponding coordinates on the mScarlet image; values below system linearity were substituted with linearly rescaled longer-exposure measurements. Five ROIs were sampled per tissue (whole brain excepted, which used a pseudo-3D volumetric ROI interpolated from ROIs spanning the structure) and averaged to a per-animal tissue mean, then averaged across animals for group-level values. Brain regions sampled (5 ROIs each): hippocampus, cortex, cerebellum (avoiding white fibrous tracts), and striatum. Spinal cord: 5 ROIs across white/gray matter at cervical, thoracic, and lumbar levels, plus separate dorsal and ventral horn sampling (thoracic, gray matter only, mutually exclusive). Peripheral organs (5 ROIs each, avoiding large vessels where applicable): liver, whole kidney, kidney cortex, adrenal glands, spleen, heart, lungs, right quadriceps, right triceps, right femoral/tibial bone marrow, and inter-scapular brown adipose tissue.

### Cell counting

#### Tissue processing

Brains were sliced sagittally to a thickness of 80 µm on a vibratome and immunostained for NeuN (neuron marker) or SOX9 (astrocytic marker) and detected with AlexaFluor-647 (AF647) secondary antibody (see IHC section for details). NeuN and SOX9 were assayed on separate sections from the same animals. Spinal cords were sucrose (15% then to 30%) embedded and then cryosectioned at a thickness of 14 µm and stained with NeuN (see IHC section for details).

#### Image acquisition

For brains, confocal 10 µm z-stacks were acquired on a Zeiss LSM 900 confocal using a 20x air objective Plan-Apochromat 20x/0.8 Air M27 (Zeiss, 420650-9902-000), with mScarlet and AlexaFluor-647 collected as separate tracks of the same field, and reduced to maximum-intensity projections for analysis. All imaging settings were matched across capsids. Cortical fields were sampled from four fields of view per animal and striatal fields were sampled from two fields of view per animal. SOX9 sampling was four fields per animal per region. No fields of view were excluded from analysis post hoc. For deep cerebellar nuclei and spinal cords, two or three sections per animal were imaged.

#### Automated segmentation

For brain cortex and striatum NeuN and SOX9, channels were segmented independently with Cellpose v3.1.0 (pretrained cyto3 model, denoising disabled) after conversion of maximum-intensity projections to single-channel grayscale. Segmentation parameters were identical between BI-hTFR1 and BI-TfR1 CapX for every marker, channel, and region. AAV9 transduction at the dose evaluated (2 × 10^12^ vg/kg) was at or below the assay’s detection floor and required different segmentation parameters.

#### Co-localization

Transduced cells were classified by cell type using a reciprocal area-overlap criterion. Candidate pairs of mScarlet^+^ and NeuN^+^ or SOX9^+^ objects were pre-selected by bounding-box intersection, then evaluated pixelwise. An mScarlet^+^ object and a marker^+^ object were scored as co-localized when their intersection covered at least 50% of the mScarlet^+^ object’s area and at least 50% of the marker^+^ object’s area. Requiring 50% in both directions prevents incidental contact between adjacent cells, or a small object lying wholly inside a much larger one, from being scored as co-localized. Transduction efficiency was calculated per cell type as the number of co-localized mScarlet^+^ objects divided by the total number of marker^+^ objects of that type. In the deep cerebellar nuclei and spinal cord cells were counted manually.

### Biodistribution

For biodistribution analysis (Figure 2), fresh frozen samples were cryopulverized for homogenization as above. DNA was isolated from cryopulverized tissues using a DNeasy 96 Blood & Tissue Kit (Qiagen, 69582) following the manufacturer’s protocol with the exception of quadriceps and triceps, which was extracted using a KAPA Express Extract Kit (Roche, KK7100) following the manufacturer’s protocol.

For biodistribution analysis (Figure 6), DNA was extracted from all tissues using the KAPA Express Extract kit; brain samples were first cryopulverized (Covaris CP02) according to the manufacturer’s protocol, whereas all other tissues were processed without cryopulverization.

qPCR standard curves were generated for the AAV WPRE transgene and *Gapdh*, then used to quantify copies of each in the isolated DNA samples. Reactions used TaqMan Universal PCR Master Mix (Applied Biosystems, 4304437) with the WPRE_F/ R/Probe, and the *Gapdh*_F/R/Probe (Table S5).

### Luciferase assays

Fresh-frozen organs were pulverized on a Covaris CP02 cryoPREP Automated Dry Pulverizer per the manufacturer’s protocol. Tissue powder was homogenized with a disposable plastic pestle in T-PER Tissue Protein Extraction Reagent containing Halt protease inhibitor cocktail (Thermo Fisher Scientific, 78425), incubated on ice for 30 min, subjected to 1-4 freeze thaw cycles, and centrifuged at 21,000 x *g* for 15 min at 4 °C. Supernatants were aliquoted and stored at −80 °C. Total protein was quantified with the Pierce BCA Protein Assay Kit (Thermo Fisher Scientific, 23225). The linear range of the luminescence assay was established for each organ by serial dilution in T-PER; lysates were normalized in T-PER to an organ-specific concentration within that range and plated at 50 µL/well in white 96-well plates (Thermo Fisher Scientific, 136101). Britelite Plus (Revvity, 6066761) was added at 50 µL/well and mixed, and each sample was transferred in technical triplicate at 25 µL/well to a white 384-well OptiPlate (Revvity, 6007290). Luminescence was read on a PerkinElmer EnVision plate reader using the Britelite program.

### *Prnp* RT-qPCR

Whole brain hemispheres were pulverized on a Covaris CP02 cryoPREP Automated Dry Pulverizer per the manufacturer’s protocol; spinal cord, liver, heart, and quadriceps were dissected into 20-50 mg pieces. Total RNA was extracted from these samples using a RNeasy 96 Universal Tissue Kit (Qiagen, 74881) with on-column DNase I digestion. Quantitative analysis was performed using the Luna Universal Probe One-Step RT-qPCR Kit (New England Biolabs, E3006X). Customized TaqMan Primers and Probes were used to detect *Prnp* (Invitrogen, Mm00448389_m1; 4351370) and an internal control *Tbp* (Invitrogen, Mm00446971_m1; 4448484) transcript. *Prnp* transcript levels were normalized to *Tbp* as an internal control and expressed as a percentage of the mean of the untreated group using the delta-delta Ct method.

### PrP ELISA

Cryopulverized brain tissue was transferred to pre-weighed 2 mL Precellys soft-tissue homogenization tubes pre-loaded with zirconium oxide beads (Pre-cellys P000912-LYSK0-A). Cold lysis buffer (0.2% CHAPS in PBS with protease inhibitor (Sigma-Aldrich 04693159001), one tablet per 10 mL) was added to 10% w/v, and samples were homogenized on a Bertin Precellys Evolution homogenizer. Homogenates were divided into 40 µL aliquots, flash-frozen, and stored at −80 °C until analysis.

PrP was quantified in these 10% homogenates using a previously described in-house cross-species ELISA^47,50^. Briefly, EP1802Y (Abcam, ab52604) was used as the capture antibody at 2.0 µg/mL and biotinylated 8H4 (Abcam, ab61409) as the detection antibody at 0.25 µg/mL, followed by streptavidin-HRP (Thermo Fisher Scientific, 21130), TMB substrate (Cell Signaling, 7004P4), and stop solution (Cell Signaling, 7002P4). Standard curves were prepared with recombinant mouse PrP (MoPrP23-231) produced in-house as described^51^; the assay has equivalent reactivity for mouse and human PrP. Values were normalized to the mean of the untreated controls and are expressed as percentage residual PrP.

### Alphafold structural modeling

A local Alphafold3 installation was used to generate docking models of the CapX-TfR1 interactions. To generate a fully formed structure around the engineered region of the capsid (3-fold axis), we included three CapX VP3 subunits (residues 226-743, chains A-C) and three chains of a previously described soluble apical domain fragment from human TfR1 (TfR1sol-apical, chains D-F)^34^. Each set of predictions was run with 11 seeds. Structural model figures were generated using Pymol 3.1.6.1.

The top ranked model of the complex was renumbered: capsid chains were offset to AAV9 VP1 numbering (UniProt Q6JC40) by adding 225 to the model position, placing the modeled VP3 fragment at residues 226–743. TfR1 chains were mapped to human TfR1 numbering (UniProt P02786) by pair-wise alignment to the full-length receptor: because the soluble apical-domain fragment carries a 7-residue linker (TGKGKSG) in place of an extended loop, model positions 1–108 were assigned TfR1 residues 197–304 (model position + 196; the linker occupies 298–304) and positions 109–156 were assigned TfR1 residues 332–379 (model position + 223).

Interface chain pair confidence scores were provided after resolving the arbitrary labeling of the three structurally equivalent TfR1 apical domains. For each model, the nine pairwise interface metrics were arranged into 3 by 3 matrices of chain-pair ipTM and chain-pair minimum predicted aligned error (PAE_min), with capsid chains (A, B, C) as rows and TfR1 chains (D, E, F) as columns. Each matrix was min–max normalized across its nine values to [0, 1]; the PAE_min matrix was inverted (1 − normalized value) so that lower PAE corresponded to higher confidence, and the two normalized matrices were averaged to give a combined score matrix. The one-to-one assignment of TfR1 chains to capsid chains that maximized the summed combined score across the three subunits was identified by evaluation of all permutations. Under this optimal assignment, the per-model mean ipTM and mean PAE_min were computed as the averages of the three assigned (raw) chain-pair ipTM and PAE_min values, respectively. This procedure was applied to all 55 models of each prediction.

Docking reproducibility across models (Figure 3C) was quantified as the Cα root-mean-square deviation (RMSD) of the TfR1 apical domain after capsid chain superposition. For each prediction, all 55 models were superposed onto the top-ranked (rank-1) CapX-human TfR1 apical-domain model by Kabsch least-squares fitting of the Cα atoms of capsid chain A, placing every model in a common capsid subunit reference frame. Holding the capsid chain fixed (no additional fitting of the receptor), the Cα RMSD between the top ranked TfR1 apical domain (chain D) and each of the model’s three TfR1 chains (D, E, and F) was computed, and the minimum for each model was reported (the best-matching copy), so that the value reflects the docked receptor pose deviation after capsid chain alignment. Small RMSDs indicate a reproducible docking pose across seeds, whereas large RMSDs indicate that AlphaFold3 did not converge on a consistent pose. Superpositions and RMSD calculations were performed with a custom Python/NumPy script implementing the Kabsch algorithm.

Predicted interface contact and human-variation analysis was defined from the top-ranked AlphaFold3 model of the CapX capsid trimer bound to the human TfR1 apical domain. A capsid residue was scored as a contact if any of its atoms lay within 4 Å (Figure 3F) or 5 Å (Table S1) of any atom of the TfR1 apical-domain chain D. Each contacting capsid residue and the human TfR1 residue(s) it approached were tabulated together with the minimum inter-atomic distance. Every human TfR1 contact position was then cross-referenced to gnomAD missense variants for *TFRC* (Ensembl transcript ENST00000360110).

### *In vitro* exogenous TfR1 expression in CHO cells

CHO cell lines were seeded at a density of 3 × 10^3^ cells. At 24 hours post-seeding cells were transiently transfected with plasmid DNA containing wild-type or mutant versions of human or macaque *TFRC* cDNA using TransIT-LT1 Transfection Reagent (Mirus, MIR2306) following manufacturer’s instruments. Transfected CHO cells were allowed to express plasmids for 24 hours and then were transduced with AAV9 or CapX encoding ssAAV-CAG-NLS-mScarlet-P2A-luc-WPRE at an MOI of 5 × 10^3^ vg/cell. Transduced cells were maintained at 37 °C with 5% CO2 for an additional 24 hours. Vector transduction was measured using the Britelite Plus (Revvity, 6066761) Reporter Gene Assay System (revvity, 6066761).

### Quantification and statistical analysis

Unless stated otherwise, no animals or sections were excluded, and statistical analyses were performed in GraphPad Prism 11.0.2. Data and code will be made publicly available upon publication.

Statistics (applies throughout unless noted). n = 6 animals per capsid (3 males, 3 females), representing biological replicates; points show individual animals and bars mean with SD. Group comparisons were performed with Brown-Forsythe and Welch one-way ANOVA followed by Šídák’s multiple comparisons test against CapX within each organ. All raw data will be included as a supplementary file with the published manuscript.

#### Figure 1H

Transduction efficiency was quantified as the percentage of marker^+^ nuclei (NeuN, neurons; SOX9, astrocytes) co-labeled with mScarlet using automated segmentation in cortex and striatum and manual counting in deep cerebellar nuclei and spinal cord. CapX was compared with BI-hTFR1 within each region by unpaired two-tailed Welch’s t test; AAV9 was not tested statistically and is reported descriptively.

#### Figure 2A

Biodistribution was quantified by qPCR with a standard curve using probes against the AAV WPRE transgene and mouse *Gapdh* DNA, as viral genome copies per diploid genome. Comparisons were made between CapX, AAV9 and BI-hTFR1. Insufficient material was recovered from one tricep sample from the BI-hTFR1 group for biodistribution analysis.

#### Figure 2B

Transduction was quantified by luciferase activity from tissue lysates normalized to total protein (µg) within each organ, within the linear detection range. Insufficient material was recovered from spinal cords from one AAV9 and one CapX animal and a heart sample from one BI-hTFR1 animal.

#### Figure 2D and 2E

Cryo-fluorescence tomography was quantified as ROI-grouped tissue mean fluorescence (see methods detail), with comparisons within each organ or region sampled. n = 2 uninjected (1 male, 1 female), n = 4 AAV9:CAG (2 male, 2 females), n= 3 CapX:CAG (1 male, 2 females), n = 3 CapX:EFS (2 males, 1 female). One CapX:CAG male animal was excluded because CFT showed reduced systemic transduction and high subcutaneous mScarlet around the injection site, and one CapX:EFS animal, which showed no minimal to no expression in any organ based on CFT.

#### Figure S3A-D

Biodistribution and transduction data from Figure 2A and B, separated by sex; comparisons were against AAV9 and BI-hTFR1. Bars show the mean with SD (two-way ANOVA) data followed by Šídák’s multiple comparison test correction

#### Figure S4C

Transduction data between *hTFRC* and hAPI using unpaired two-tailed Welch’s t test on log-transformed data; none reached statistical significance.

#### Figure 4A-D

Viral genomes were quantified by dPCR using an WPRE probe normalized to diploid genome measured by a mouse *Gapdh* probe. In 8 of 36 animals we observed AAV-PHP.eB brain biodistribution having 7.5- to 450-fold below their cohort-matched animal medians (7 out of 8 animals) and non-detectable in the brain biodistribution in the eighth. In the liver, 7 out of these 8 animals were 8- to 29-fold below their cohort medians (Figure S7). As noted in the results section, the deficit was specific to AAV-PHP.eB. In the same animals, the other three capsids were within ∼2.6-fold (brain) and ∼3.6-fold (liver) of their cohort medians; across all 36 animals and all four tissues. All four capsids were co-administered from the same mixture and quantified from the same samples and probe-primer stocks, therefore, the deficit is unlikely attributable to experimental conditions. These low responders were excluded post hoc from AAV-PHP.eB-specific analyses.

#### Figure 5B and C

Dual AAV base editing to install R37X into human *PRNP* was assessed by NGS, and prion protein levels were measured by ELISA and compared across the two doses relative to untreated controls. n = 6 uninjected animals (all males); n = 6 animals (3 males and 3 females) per treated dose. Unpaired two-tailed Welch’s t test was used to compare the two doses. No comparisons were made to uninjected animals.

#### Figure 6A

Biodistribution across the 4 doses of CapX and AAV-PHP.eB was quantified as in Figure 2A. At each matched dose within each organ, an unpaired Welch’s t test was performed between the two capsids on lognormal data. Four PHP.eB and four CapX animals were excluded post hoc as they showed no viral genomes across the 5 organs tested using qPCR, suggesting that they weren’t dosed properly. One AAV-PHP.eB animal was found dead 4 days after being injected from a 6 × 10^12^ vg/kg injection, the cause was undetermined. A heatmap showing all animals used in the study with the *Prnp* silencing levels can be found in (Figure S11).

#### Figure 6B-D

Remaining mouse *Prnp* transcripts in brain and liver were measured by a qPCR probe and remaining mouse PrP in brain by ELISA (see Method details). At each matched dose within each organ, an unpaired Welch’s t test was performed between the two capsids. Animals that were excluded post hoc from Figure 6A were also excluded from analysis in these panels.

## Acknowledgements

Andrew J. Barry for building an open-source drag and drop plotting, data selection, and filtering software for the lab that was used to visualize NGS data. Jarrett Rios with assistance with the NGS data processing pipelines. Jason Wu, Charles Vanderburg, and Nuria R. Botticello-Romero for assistance with tissue collection. Jenna Hurley in help with tissue processing and laboratory operations. Fred Roberts, Rachel Friedland, Matt Silva and the EMIT Imaging team who performed CFT tissue imaging and post imaging analysis. Andrew Steinsapir, Chris Davis, Jorge Santiago-Ortiz, Von Wiltman, and Diego Garzón, Yujia A. Chan, Yuan Yuan, and Albert T. Chen for helpful discussions and comments on the manuscript. Tyler Caron, Nathan Chan, and the Comparative Medicine staff at the Broad Institute for their support and discussions related to animal experiments. Samantha Moore, Amanda Clark, Mina Yamini, Caroline Atkinson, and Ed Clark at The Division of Comparative Medicine Pathology Core at MIT for their technical support with necropsy, tissue processing, and slide preparation. Garam Kim for assistance with imaging processing pipelines. Meiru An and David Liu for providing dual base editor AAV plasmids. Elvira Dzhura and Abigail Desrochers in the Stanley Center for Psychiatric Research, and Brian Buckley, Simone Lord, and Lizzie Schaeffer in Chemical Biology and Therapeutics Science at the Broad Institute, for laboratory operations support.

## Author contributions

Conceptualization: K.Y.C., Q.H., and B.E.D. Methodology: K.Y.C., C.-Y.L., and B.E.D. Software: K.Y.C., D.A.S., and A.S. Validation: K.Y.C. and D.A.S. Formal analysis: K.Y.C., D.A.S., M.B., and B.E.D. Investigation: K.Y.C., C.-Y.L., S.P., Q.H., Q.Z., J.W.H., J.J., F.S., J.R.D., N.G.K., S.J., and B.E.D. Resources: J.W.H., P.P.B., F.S., E.V.M., and S.M.V. Data curation: K.Y.C., C.-Y.L., and B.E.D. Writing original draft: K.Y.C. and B.E.D. Writing - review & editing: all authors. Visualization: K.Y.C. and B.E.D. Project administration: K.Y.C. and B.E.D. Supervision: K.Y.C., S.P., and B.E.D. Funding acquisition: E.V.M, S.M.V., and B.E.D.

## Declaration of generative AI and AI-assisted technologies in the writing process

During the preparation of this work, the authors used Claude (4.6, 4.8, and 5.0) for error checking and to improve clarity. After using these tools, the authors reviewed the edited text and take full responsibility for the content of the publication.

## Lead contacts

Information requests should be directed to K.Y.C. and B.E.D.

## Materials availability

Plasmids used in this study will be deposited to Addgene.

## Data and code availability

Upon publication, all data will be available in the manuscript and its supplementary material, and all code related to the cell-counting analysis (Figure 1), RMSD analysis (Figure 3), and kinetics analysis (Figure 4) will be made publicly available.

