## Supplementary Data for "BI-TfR1 CapX rapidly transits the blood-brain barrier for efficient, low-dose gene delivery throughout the CNS"

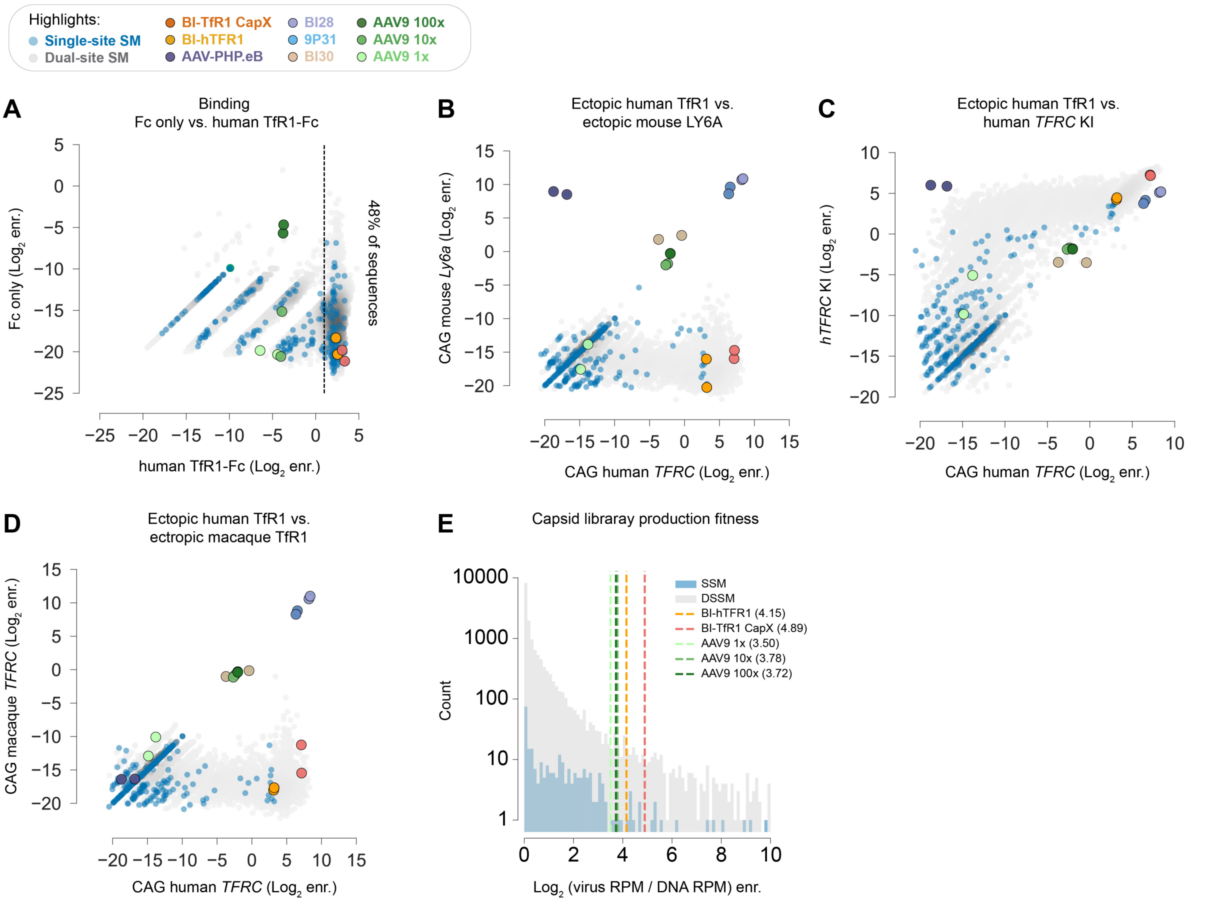


**Figure S1. Development of CapX through a single round of *in vitro* and *in vivo* screens for TfR1-dependent functions and production fitness related to Figure 1.** (**A**) $\text{Log}_2$ enrichment (enr.) of Fc-control vs hTfR1-Fc binding of the library variants, dotted vertical line demarcates variants defined as TfR1 binding ($\text{Log}_2$ enrichment ≥ 1). (**B**-**D**) The plots show $\log_2$ $\log_2$ enrichment of capsid library mRNA expression in the brains of mice transduced with AAV-BI30 packaging the indicated receptor expressed from a CAG promoter ("two-step" ectopic receptor assay, see Method details): (**B**) CAG-mouse *Ly6a* vs. CAG-human *TFRC*, (**C**) *hTFRC* KI mice vs CAG-human *TFRC*, and (**D**) CAG-rhesus macaque *TFRC* vs CAG-human *TFRC*. (**E**) $\text{Log}_2$ enrichment of production fitness for the indicated capsids with AAV9 reference sequences.

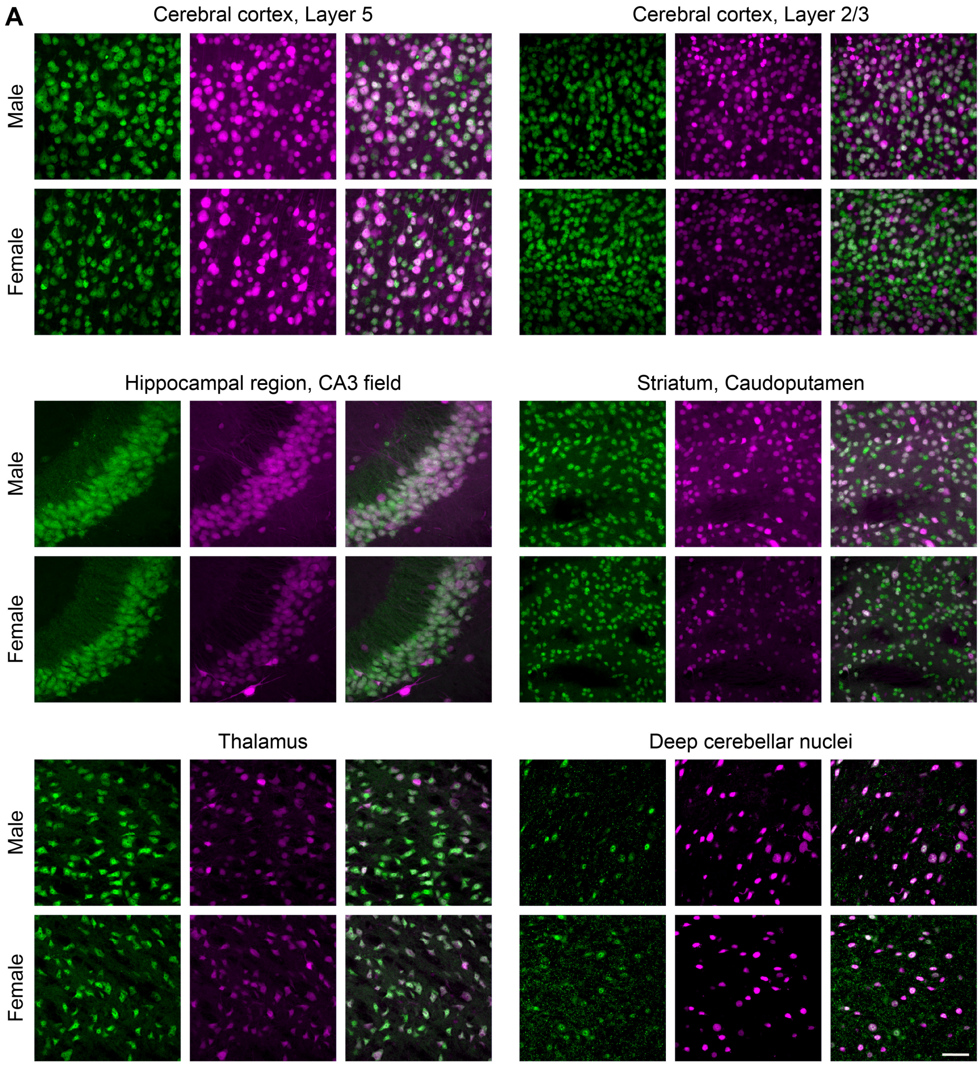


**Figure S2. CapX transduces neurons across multiple brain regions.** (**A**) Mice from the same cohort shown in Figure 1E-1H. Images show neurons (NeuN, green), transduced cells (NLS-mScarlet, magenta), and the overlay in the indicated brain regions. Scale bar: 20 µm.

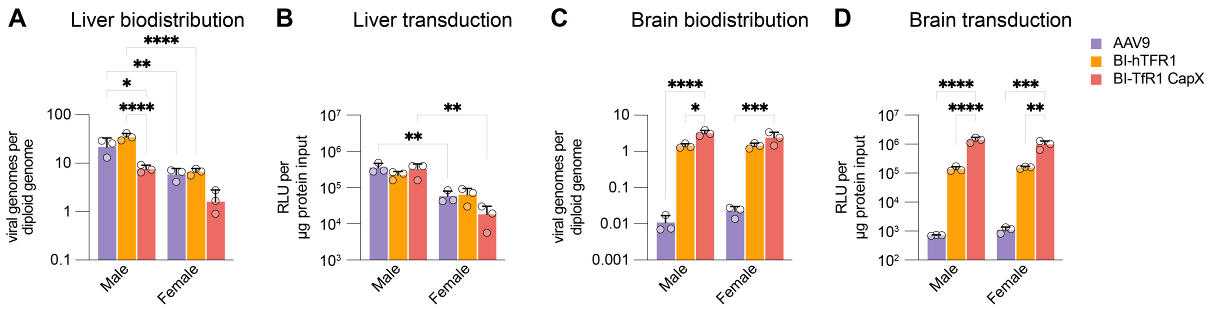


**Figure S3. Sex influences liver but not brain biodistribution and transduction.** Mice from the Figure 1 and Figure 2 cohort were assessed for liver biodistribution (**A**) and transduction (**B**) and brain biodistribution (**C**) and transduction (**D**), with the data segregated by sex. We observed a significant sex x capsid interaction for liver biodistribution (p = 0.003), but not for liver transduction or for brain biodistribution or brain transduction. Data points represent individual mice (n = 6 mice per group) and bars show the mean with SD (two-way ANOVA) data followed by Šídák's multiple comparison test correction; *p <0.05, **p < 0.01, ****p < 0.0001).

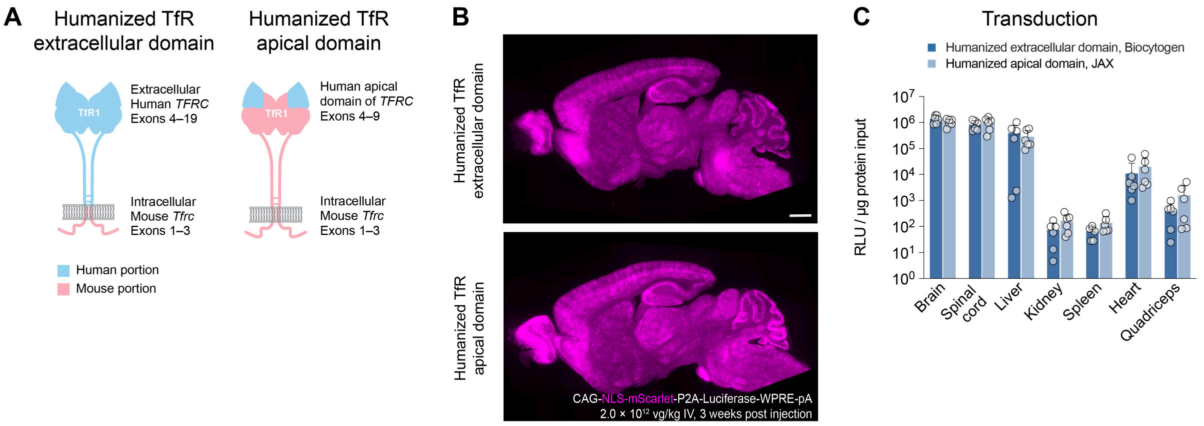


**Figure S4. CapX is compatible with multiple *hTFRC* KI mice.** (**A**) The schematics show representations of the humanization of two available *hTFRC* KI mice with the humanized areas shown in blue and the regions of the proteins that remain murine in pink. The indicated *hTFRC* KI mouse lines were injected with CapX at 2 x $10^{12}$ vg/kg (n = 6 mice per group) and assessed for mScarlet expression in sagittal brain sections (**B**) and for luciferase expression in the indicated organ (**C**) at three weeks post intravenous administration. Data points represent individual mice; bars show the mean with SD. Comparisons within each organ were made using unpaired two-tailed Welch's t test on log-transformed data; none reached statistical significance.

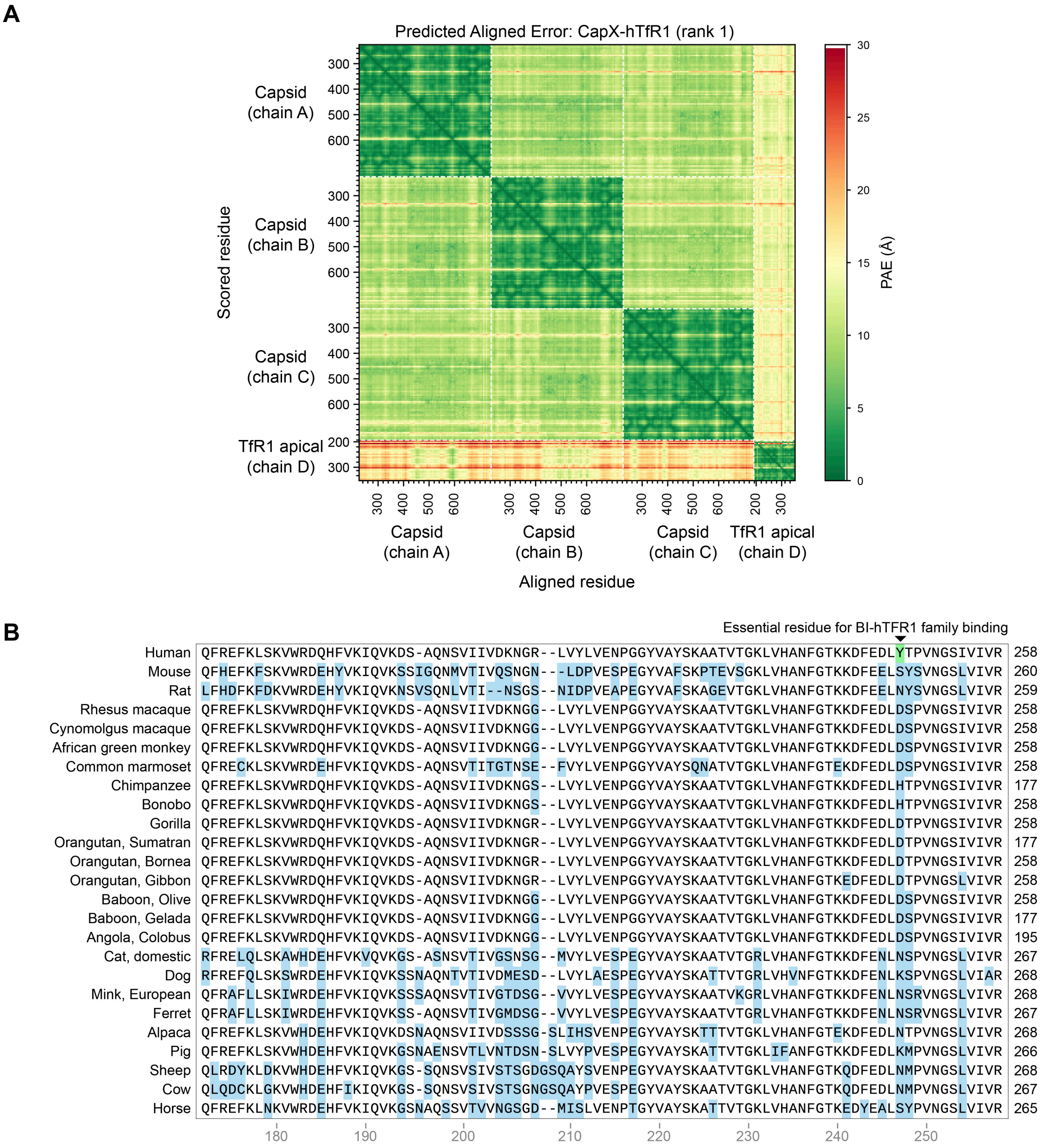


**Figure S5. Tyr247 is unique to humans and docking figures supporting Figure 3.** (**A**) Predicted aligned error (PAE) for the top-ranked AF3 model of the BI-TfR1 CapX trimer (capsid chains A, B, C) bound to one human TfR1 apical domain (chain D). Color indicates the expected position error (Å) for each scored residue (rows) when the model is aligned on each aligned residue (columns), from low (green) to high (red). Low-PAE blocks along the diagonal indicate confidently predicted intrachain folds, whereas off-diagonal blocks report the confidence in the relative positioning of each chain pair. In the capsid–TfR1 (A/B/C vs D) blocks, the lowest inter-chain PAE is concentrated at the engineered loop region (CapX Chain A residues 586–596). (**B**) The figure shows an alignment (MUSCLE, Snapgene) of the human TfR1 apical domain amino acid sequence surrounding position 247 with the sequences from other species.

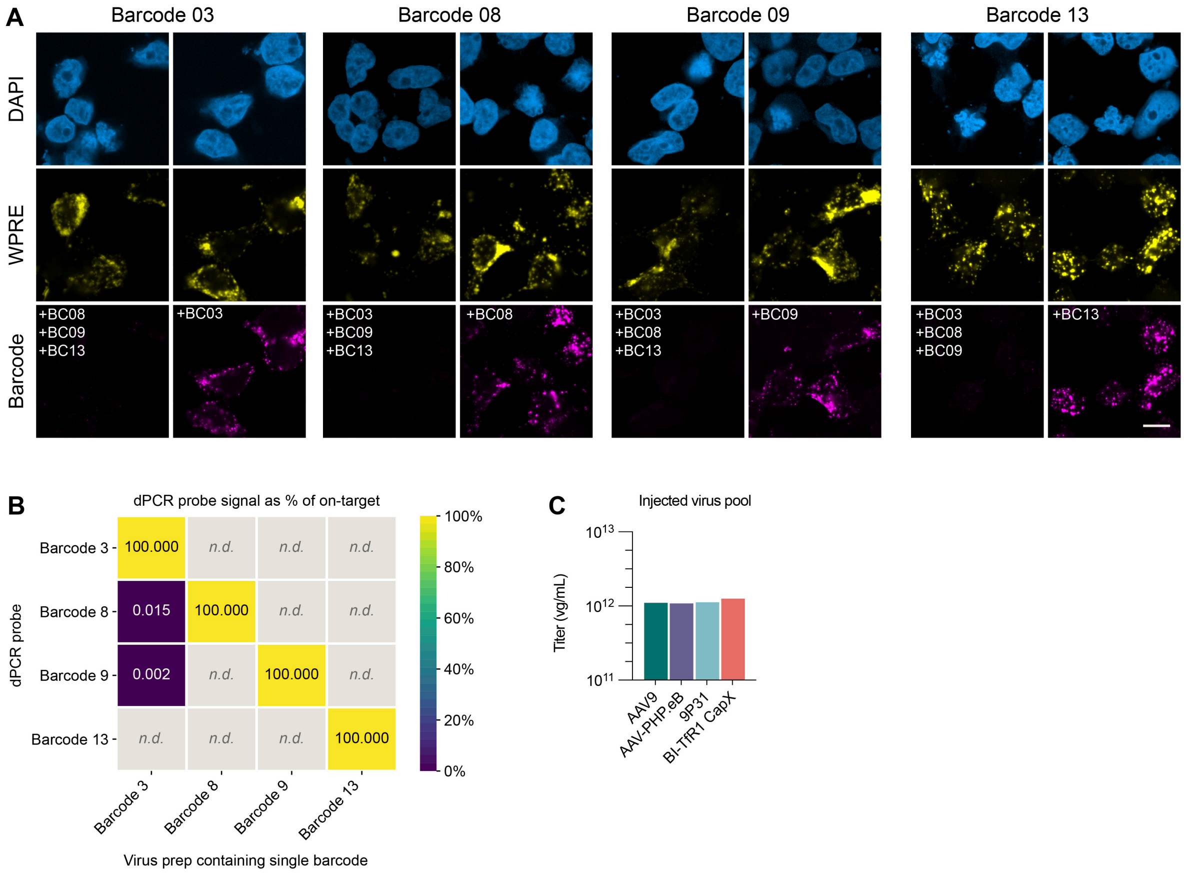


**Figure S6. The barcode-specific RNAscope and dPCR probes used in Figure 4.** (**A**) HEK293T cells were transfected with a single barcoded superfolder GFP (sfGFP) expressing reporter (ssCAG-H2B-sfGFP-WPRE-barcode-pA) or with a mix of the other three barcodes used in Figure 4. Each condition was then probed with the RNAscope sense probe against the indicated barcode (magenta) and against the WPRE element common to all four constructs (yellow). (**B**) Each uniquely barcoded AAV prep was assayed with all four barcode-detection dPCR probes. Values show the mean off-target signal as a percentage of the on-target probe concentration; *n.d.* denotes zero positive partitions in dPCR. (**C**) Titers of the injected AAV pool after pooling, as used in Figure 4. Scale bar: 10 μm.

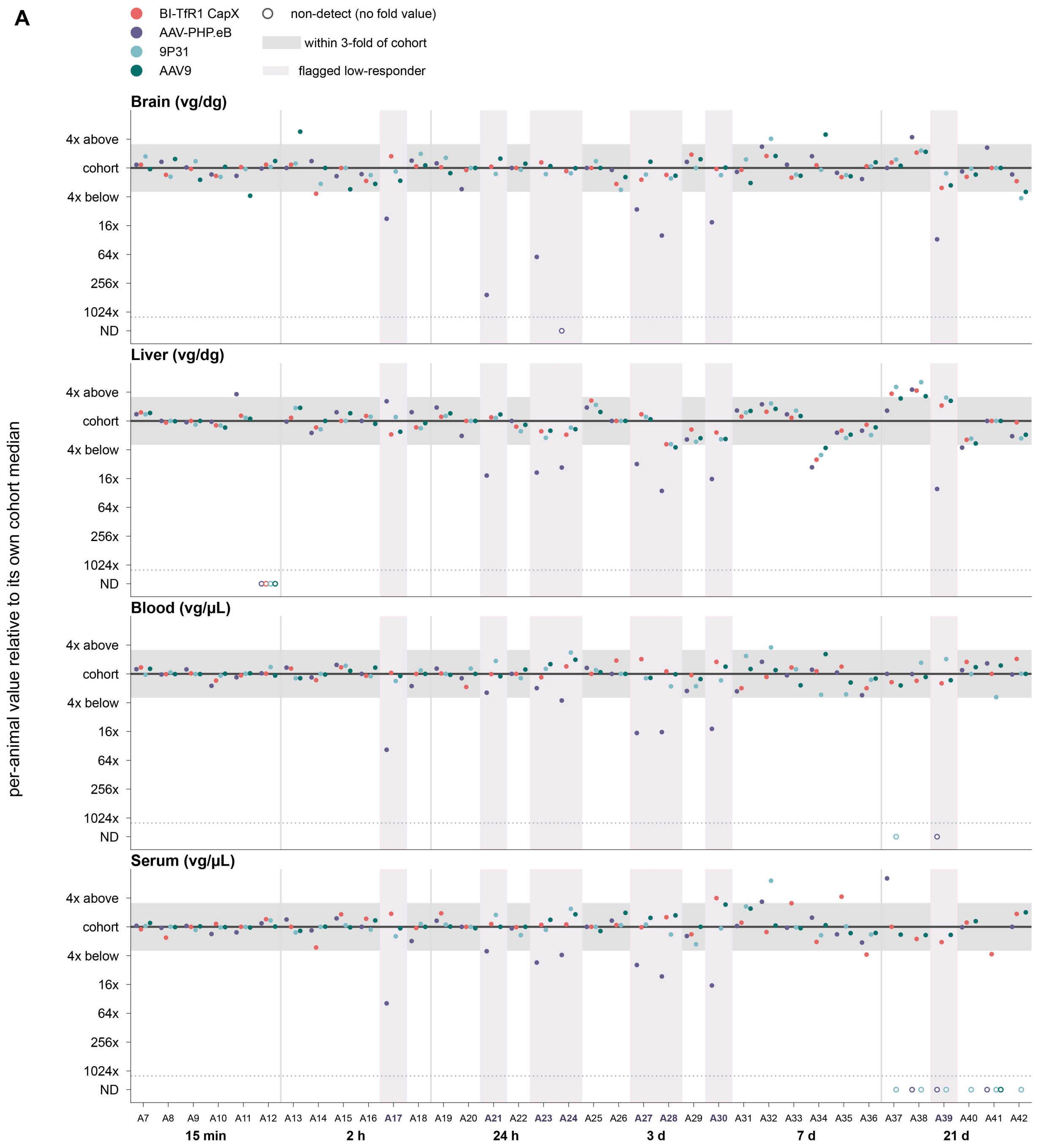


**Figure S7. Biodistribution of AAV-PHP.eB is specifically reduced or absent in a subset of animals injected with the barcode AAV pool used in Figure 4 .** Each of 36 *hTFRC* KI animals (n = 6 at each of six timepoints) received a single injection of the pool of four AAVs: AAV-PHP.eB, BI-TfR1 CapX, 9P31 and AAV9, each packaging a distinct barcoded genome. The barcoded genomes were quantified by dPCR from the same samples. Within each timepoint, the 6 animals are plotted as separate clusters labelled by animal number (A7-A42, six control animals A1-A6 are not shown). Each point is that animal's vg/dg divided by the median of the same capsid at the same timepoint, so every capsid is compared only with itself ($\log_2$ scale). The gray band shows the range within 3-fold of the per capsid-timepoint median; tinted columns flag animals as AAV-PHP.eB low responders (A17, A21, A23, A24, A27, A28, A30, A39). Open circles on the reserved row below the dotted rule (ND) are non-detected samples.

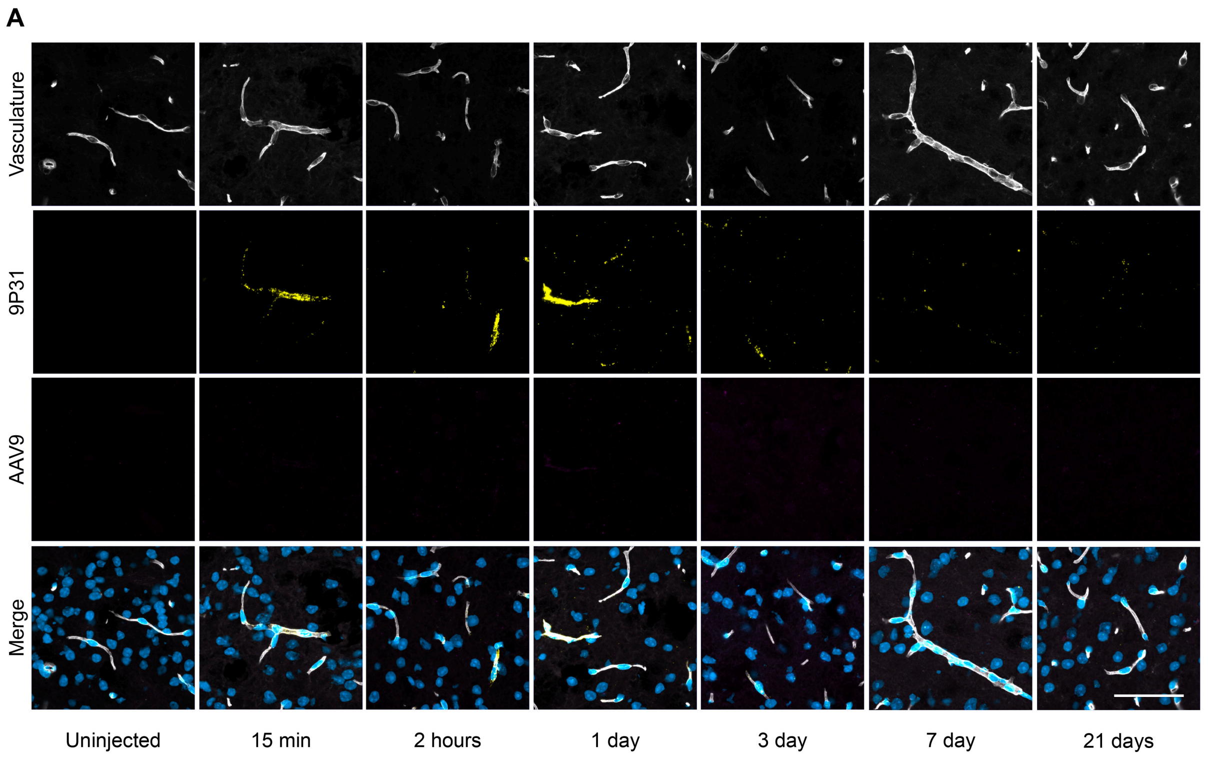


**Figure S8. BBB transport data with 9P31 and AAV9, related to Figure 4.** Representative RNAscope images from the brain at different time points post administration of the AAV pool. Images show vasculature (lycopersicon esculentum lectin, white), 9P31 genomes (yellow), AAV9 (magenta), and nuclei (DAPI, blue). Scale bar: 50 µm.

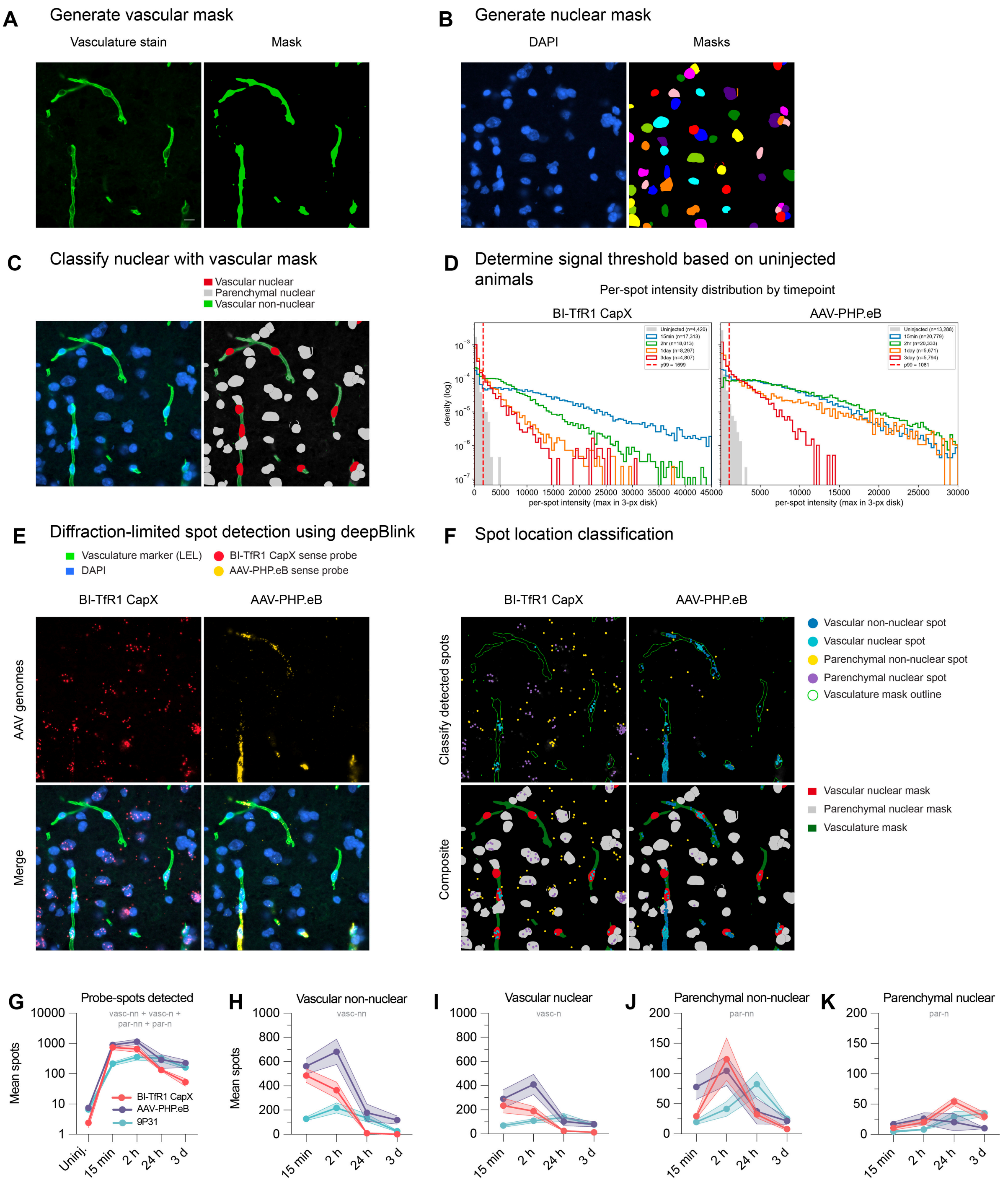


**Figure S9. Image analysis pipeline used to detect and classify RNAscope spots in Figure 4.** (**A**) Masks are generated based on the vascular marker. (**B**) Nuclear masks are generated by segmenting the DAPI+ DNA signal. (**C**) Nuclear masks are classified as vascular or parenchymal. (**D**) The threshold for calling signal from the RNAscope *in situ* hybridization sense probe against the AAV-barcoded genome (red dotted line) was set to exclude 99% of the signal in uninjected animals, and the same threshold was applied to all samples imaged with that probe under identical settings. (**E**) Using deepBlink, a neural network-based method to detect diffraction-limited spots. (**F**) Classification of spots detected into compartments. (**G**) Total spots detected using deepBlink for CapX, AAV-PHP.eB, and 9P31. (**H-K**) Mean probe spots across 3 fields of view per animal averaged in each of four compartments (vascular non-nuclear, vascular nuclear, parenchymal non-nuclear, or parenchymal nuclear). Data points represent mean with SEM; n = 6 animals per timepoint with 3 females and 3 males; unless otherwise stated (see Method details).

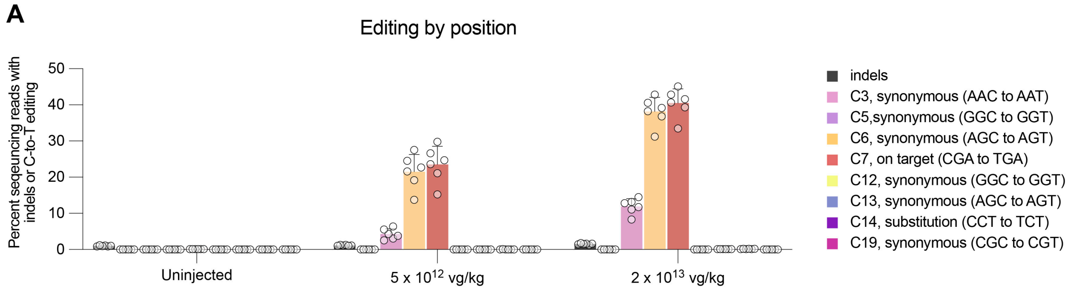


**Figure S10. Target and bystander *PRNP* CBE edits, related to Figure 5.** (**A**) Frequency of *in vivo* bystander and on-target (C7) edits. Data points represent individual transgenic mice containing *human TFRC* and human *PRNP* (see methods), (n = 6 animals per dose) and bars show the mean with SD, no statistical tests were performed.

A

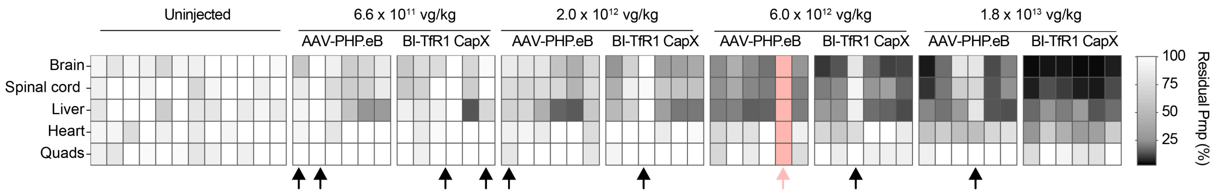


**Figure S11. Relative *Prnp* mRNA levels across organs within each animal related to Figure 6.** Dark arrows indicate animals that had no detectable viral genomes across the brain, spinal cord, liver, heart and quads, and were therefore excluded from the analysis. Pink arrow indicates an AAV-PHP.eB:Prnp-CHARM-treated animal that was found dead from an unknown cause 4 days post injection.

A

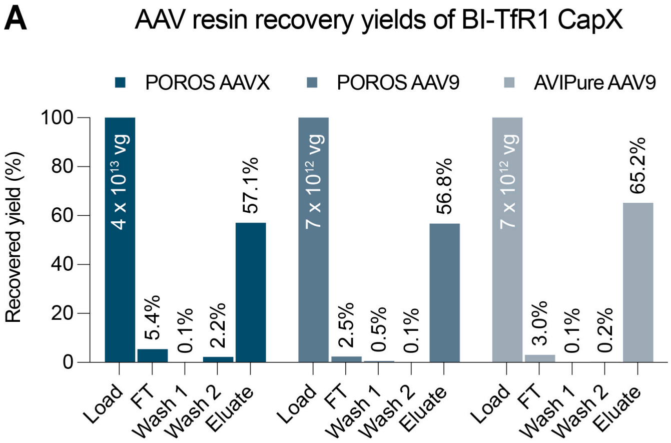


**Figure S12. CapX compatibility with commercial affinity resins.** CapX preparations were loaded onto three different commercial affinity resins as specified, and the percentage of vector genomes recovered was measured by ddPCR in the flowthrough (FT), first and second wash, and neutralized elution.
