## Supplementary material for "BI-TfR1 CapX rapidly transits the blood-brain barrier for efficient, low-dose gene delivery throughout the CNS": Data S1

Prepared March 2026 by:
*Magalie Boucher, DVM, MS, DACVP*
*Assistant Director/ Comparative Pathologist*
*Division of Comparative Medicine*

**Summary:**

The objective of this study was to evaluate the comparative toxicity profiles of the human transferrin receptor-targeting AAV vector (BI-TfR1 CapX) and the benchmark AAV9 vector in humanized *TFRC* (B-hTFR1) mice. Assessments were conducted at 3 and 21 days post-administration.

Under the conditions of this study, there were no treatment-related effects on mortality, body or organ weights, clinical pathology (hematology), or gross and microscopic morphology. Both AAV vectors were well-tolerated and showed no discernible toxicity compared to the PBS vehicle control.

**Study design:**

| | | Day 3 | | Day 21 (3-weeks) | |
|---|---|---|---|---|---|
| **Groups** | Dose | Sex | | Sex | |
| | | Males | Females | Males | Females |
| **PBS** | NA | 3 | 3 | 3 | 3 |
| **AAV9** | 1E13 vg/kg | 3 | 3 | 3 | 3 |
| **BI-TfR1 CapX** | 1E13 vg/kg | 3 | 3 | 3 | 3 |

**Tissue List:**

| Slide # | Tissue (s) |
|---|---|
| 1 | Brain (1/2 will be submitted in formalin) and half will be OCT frozen by the investigator |
| 2 | Heart and Kidneys |
| 3 | Lung and esophagus/trachea in cross section |
| 4 | Liver (left lateral, medial lobe/gall bladder) |
| 5 | Adrenals, thymus, spleen |
| 6 | Stomach |
| 7 | Duodenum/pancreas, jejunum, cecum |
| 8 | Ileum, colon, mesenteric lymph node |
| 9 | Muscle (bicep femoris) longitudinal and transverse including the sciatic nerve |
| 10 | Urinary bladder with Ovary or Testis (unilateral)/other half OCT frozen by the investigator |
| 11 | Sternum/bone marrow |

| | |
|---|---|
| 12 | Spinal column/spinal cord and DRG(cervical, thoracic, lumbar) |

**<u>Results:</u>**

**Body Weight:**

Interpretation of body weight data was impacted by two primary confounding factors: instrumental variance and baseline age differences.

Weights recorded at Day 0 utilized a different scale than those at Day 3 and 21, introducing potential instrumental bias. However, as these fluctuations were mirrored in the PBS control group, they were attributed to methodological variance rather than test-article-related toxicity.

Similarly, the small cohort size (n=3/sex/timepoint) and non-uniform ages at the start of the experiment resulted in inherent variability that precluded the detection of subtle shifts. Consequently, analysis focused on identifying broad clinical trends. As no clear difference in body weight (percent change from baseline) was identified between AAV-treated and control groups, no treatment-related effects on body weight were concluded.

**Organ weight:**

Minor variation in both absolute and relative (percent of body weight) organ weights were observed across treatment and control groups. These changes were not associated with macroscopic or microscopic findings and were not considered treatment related.

The only exception was a single female (Animal No. 212) in the AAV9 group at Day 21, which had increased absolute and relative left kidney weight. This change correlated with the gross finding of enlarged kidney at necropsy and the microscopic findings of chronic hydronephrosis and pyelonephritis histologically. Given the unilateral nature of this lesion and the known sporadic occurrence in mice (Imai, 2024), the increased kidney weight was interpreted as a spontaneous, incidental finding unrelated to AAV administration.

Reference:
Imai DM, Barthold SW, Griffey SM, Percy DH. Pathology of Laboratory Rodents and Rabbits, 5th ed. Hoboken, NJ: Wiley-Blackwell; 2024

**Hematology:**

Hematologic parameters exhibited minor fluctuations across the control and both AAV treatment groups (AAV9 and BI-TfR1 CapX). No consistent trends were observed, and variations were often isolated to individual animals. Due to the small group size (n=3) and age variability within groups, no definitive relationship to either viral vector was identified. No treatment-related effects on the hematology parameters evaluated were concluded.

**Histopathohology:**

There were no macroscopic or microscopic changes related to AAV9 or AAV (BI-TfR1 CapX) in either male or female B-hTFR1 mice. All microscopic changes observed were mostly minimal, did not worsen or lessen between Day 3 and 21 and have been reported as background and incidental findings in mice. Such findings were considered incidental and unrelated to treatment, falling within the range of expected biological variation for these mice.

References:

Pathology of the Mouse: Reference and Atlas (Maronpot RR, Boorman GA, Gaul BW, eds). Cache River Press, Vienna, IL.
International nomenclature and diagnostic criteria for lesions in rats and mice (INHAND). https://www.toxpath.org/inhand.asp
Nonneoplastic Lesion Atlas. A guide for standardizing terminology in toxicologic pathology for rodents. https://ntp.niehs.nih.gov/atlas/nnl

The raw data is located in (Table S2).
For bodyweights please refer to (Table S2, Body Weights)
For organ weights please refer to (Table S2, Organ Weight)
For complete blood count please refer to (Table S2, CBC Results)
For histopathology results please refer to (Table S2, Histopathology)
