## Supplementary material for "BI-TfR1 CapX rapidly transits the blood-brain barrier for efficient, low-dose gene delivery throughout the CNS": Data S2

### PRNP-CHARM-001

**STN: PS009720**

### Intravenous AAV delivered epigenetic editor targeted to *PRNP* to treat prion disease

**Information Package for INTERACT Meeting**

**Submitted: November 25, 2024**

**Sponsor**
**Sonia M. Vallabh PhD**
**Eric V. Minikel PhD**
**Broad Institute**
**75 Ames St 8047**
**Cambridge MA 02142**

**1. EXECUTIVE SUMMARY** **3**
1.1. Background on prion disease and the therapeutic hypothesis of PrP lowering 3
1.2. Proposed therapeutic 3
1.3. Envisioned clinical trial design 4
1.4. Questions 7
**2. LIST OF FIGURES AND TABLES** **7**
**3. PRODUCT NAME** **8**
**4. CHEMICAL NAME AND STRUCTURE** **8**
**5. PROPOSED INDICATION** **9**
**6. DOSAGE FORM AND ROUTE OF ADMINISTRATION** **9**
**7. PURPOSE AND FORMAT OF MEETING** **9**
**8. BACKGROUND ON PRNP-CHARM-001** **9**
*8.1. BACKGROUND ON PRION DISEASE AND THE THERAPEUTIC STRATEGY OF PRP LOWERING 9*
8.1.1. Prion disease is a fatal neurodegenerative disease with no existing cure caused by PrP 9
8.1.2. PrP lowering is a well-validated therapeutic hypothesis in prion disease 10
8.1.3. Neurons are the cell type of interest for PrP-lowering therapies 13
8.1.4. PrP is dispensable for healthy life 14
8.1.5. Symptomatic and pre-symptomatic patients present opposing clinical opportunities and challenges. 15
8.1.6. Key points for treatment protocols 17
8.1.7. Relevant history with CDER 18
*8.2. BACKGROUND ON TARGETED EPIGENETIC SILENCING USING CHARM 19*
8.2.1. Epigenetic editing as a therapeutic intervention for prion disease 19
8.2.2. CHARM can be used to silence PRNP 21
*8.3. BACKGROUND ON DELIVERY VIA A SYSTEMICALLY DELIVERED BLOOD-BRAIN BARRIER-CROSSING AAV CAPSID 24*
8.3.1. The BI-hTFR1v2 capsid enables dramatically improved CNS delivery compared toAAV9 25
8.3.2. The CNS tropism of BI-hTFR1v2 is specific to human TfR1 28
9. STATUS OF PRODUCT DEVELOPMENT 29
9.1. GENERAL INVESTIGATIONAL PLAN 29
9.2. CHEMISTRY, MANUFACTURING AND CONTROL (CMC) DATA 30
9.3. PHARMACOLOGY AND TOXICOLOGY INFORMATION 30
9.3.1. Off-target assessment 31
9.3.2. In vivo studies 31
9.4. SUMMARY OF PREVIOUS HUMAN STUDIES 35
9.5. SPECIFIC OBJECTIVES/ OUTCOMES EXPECTED FROM MEETING 35
**10. LIST OF QUESTIONS AND SPONSOR POSITIONS** **36**
**11. PROPOSED AGENDA** **39**
**12. SPONSOR PARTICIPANTS** **39**
**13. LIST OF ABBREVIATIONS** **39**
**14. REFERENCE LIST** **41**

### 1. SUMMARY

This meeting is to discuss an investigator-initiated effort to advance a prion protein (PrP)-lowering AAV-delivered epigenetic editor, PRNP-CHARM-001, envisioned as a one-time, permanent therapy for the treatment and prevention of prion disease.

We have endeavored to outline all key points of our meeting materials in this Executive Summary, while providing links to further detail on key topics in the main text.

#### 1.1. Background on prion disease and the therapeutic hypothesis of PrP lowering

Prion disease is a rapidly progressive neurodegenerative disease. From first symptoms, patients steeply decline into profound dementia, losing faculties over the course of weeks and typically dying within five months (8.1.1). The disease is universally fatal, and there currently exists no standard of care.

The pathogenesis of prion disease is well understood. A single protein, prion protein (PrP), encoded by the prion protein gene (*PRNP*) misfolds into a protein-only pathogen which spreads across the brain, killing neurons (8.1.1). PrP is strictly required for disease, as evidenced by decades of genetic experiments in animal models. More recently, we have shown with PrP-lowering antisense oligonucleotides (ASOs) that lowering PrP levels in the brain both delays onset and slows progression of prion disease across a range of treatment paradigms (8.1.2). While these data provide foundational proof of concept for PrP lowering, we are motivated to develop a one-time PrP-lowering gene therapy that may prove more potent, better tolerated and deliverable to the whole brain.

Both animal and human genetic data indicate that PrP reduction is well tolerated. Homozygous PrP knockout mice have no striking phenotype beyond their well-established invulnerability to prion disease (8.1.4).

#### 1.2. Proposed therapeutic

There are multiple possible paths to attempt to reduce PrP expression in the brain. As PrP is nonessential, a single treatment to permanently shut off the *PRNP* gene is attractive. To this end, our proposed therapeutic seeks to irreversibly methylate the *PRNP* promoter. It is composed of a compact zinc finger (ZF) domain targeted at the *PRNP* promoter, genetically fused to an effector domain that recruits the endogenous methyltransferase DNMT3A1 already present in the cell (8.2.1).

Together, this therapy, known as PRNP-CHARM (where CHARM stands for Coupled Histone tail for Autoinhibition Release of Methyltransferase), is of a size deliverable to the CNS in a single AAV capsid. Note that this product contains no nuclease domain; its mechanism relies on methylation-mediated silencing of the *PRNP* locus, not on DNA cleavage or editing.

Because prion disease is driven by neurons across the whole brain (8.1.3), meaningful biodistribution will depend on an AAV capsid capable of crossing the blood-brain barrier in adults following intravenous delivery. The capsid we will use for delivery is BI-hTfR1v2 , a novel AAV9-derived capsid engineered to provide efficient gene delivery to neurons and glial cells throughout the CNS (8.3.1). The capsid has been modified such that it binds the apical domain of the human transferrin receptor (hTfR1), allowing it to cross the blood brain barrier.

Due to an amino acid change present in the binding site of TfR1 in humans compared to other species, BI-hTfR1v2 does not bind the transferrin receptor in non-human primates or any other species besides human (8.3.2), therefore CNS biodistribution would not be expected. BI-hTfR1v2 is uptaken into the CNS of TfR-humanized mice.

The power of the PRNP-CHARM approach has been demonstrated both in wild-type mice and in mice engineered to carry the human *PRNP* gene.

#### 1.3. Envisioned clinical trial design

Given its high potential to be protective against prion disease, we are working to swiftly advance PRNP-CHARM-001 to testing in humans in a series of investigator-initiated trials.

The key endpoints in an initial clinical study of PRNP-CHARM-001 will be 1) safety, and 2) target engagement: measuring whether the drug lowered PrP levels in the brain. It has been shown in rats that brain PrP levels can be tracked pre- and post-treatment using cerebrospinal fluid (CSF) PrP levels as a proxy (8.1.2).

Two populations stand to benefit from a PrP-lowering therapy: 1) symptomatic prion disease patients, and 2) carriers of high penetrance *PRNP* variants that reliably cause fatal prion disease.

Treatment of presymptomatic carriers, as well as symptomatic patients, is supported by the following parameters:

- PrP is well established as the single causal protein in prion disease (8.1.1).
- PrP is expendable for healthy life (8.1.4).
- Once symptoms begin, prion disease is rapidly lethal (8.1.5); treatment at this stage cannot address irreversible neuronal loss already incurred.
- Preclinical studies indicate that presymptomatic treatment can extend healthy life, and is most effective if initiated before pathology is evident (8.1.2).
- Genotypes corresponding to a >90% lifetime risk of fatal prion disease can be identified with confidence before symptom onset (8.1.5).
- Life tables indicate the distribution of lifetime risk according to *PRNP* variant (8.1.5).
- Lowering of brain PrP, the causal molecule in prion disease, can be monitored by measuring PrP in CSF (8.1.5). This can be done in the absence of the disease process, as native PrP is present in all humans.

The symptomatic and presymptomatic populations provide distinct opportunities for data readout and for clinical benefit (8.1.5), as outlined below.

| | **Symptomatic patients** | **Presymptomatic at-risk individuals** |
|---|---|---|
| Safety | Safety assessments will be **confounded** by rapid disease progression and limited to overt toxicity and immunogenicity. | As *PRNP* mutation carriers are healthy prior to symptom onset, safety **can be reliably assessed** in detail. |
| Target engagement | Measurement of drug-associated changes in CSF PrP levels will be **confounded** by disease-associated changes in CSF PrP levels. | As CSF PrP levels are longitudinally stable presymptomatic individuals (mean CV = 10% across genotypes over up to six years, (8.1.5), drug-associated changes in CSF PrP levels **can be reliably measured**. |
| Prospect for benefit | Potential benefit is possible, but **limited** by 1) rapid disease progression, 2) the 4-week time to full effect of the drug on the protein level, based on PrP's 5-day in vivo half life, and 3) lack of basis on which to expect reversal of accumulated symptoms caused by neuronal loss. | In presymptomatic carriers the drug will have full time to reach maximal target engagement and the **prospect to extend healthy life.** |
| Risk vs. benefit | Risks of drug are **reasonable** given this population's imminently fatal diagnosis and the lack of standard of care for prion disease. | Risks of drug are **reasonable** in the context of 90%+ penetrant *PRNP* variants, given the rapid lethality of disease once it strikes. Expected permanence of therapy will be offset by specificity of the drug and the non-essentiality of the *PRNP* gene. Recruitment can bias towards older individuals closer to expected age of disease onset. |

Given their distinct opportunities and challenges, symptomatic and presymptomatic treatment are best conceived as two independent clinical paths, neither of which gates the other. We envision conducting two small (~10 patients each) independent investigator-initiated trials, as outlined below.

*Clinical study draft protocols*

| Study population | **Presymptomatic adult carriers of high risk *PRNP* variants** | **Symptomatic prion disease patients** |
|---|---|---|

| | | |
|---|---|---|
| Eligibility criteria | Positive test for high penetrance pathogenic *PRNP* variant: E200K, D178N, P102L, P105L, 5-OPRI, 6-OPRI, A117V, G198S<br>● Though age of onset cannot be predicted for a given individual, enrollment can be biased towards older individuals closer to expected age of onset based on genetic prion disease life tables.<br>● Note: As pathogenic biomarkers do not reliably predict onset of genetic prion disease, they will not determine enrollment. | Probable prion disease by CDC diagnostic criteria [CDC criteria 2021], including either:<br>● Positive spinal fluid RT-QuIC<br>● Positive *PRNP* mutation test<br>MRC Prion Disease Rating Score >13 |
| Dose schedule escalation/ study design | ● Low single dose - 2 patients<br>● Mid single dose - 2 patients<br>● High single dose - 2 patients<br>● High* single dose - 4 patients<br>Each escalating dose cohort will be gated by an IDMC determination based on interim safety data.<br>**Or highest dose deemed tolerated based on interim analyses.* | ● Low single dose - 2 patients<br>● Mid single dose - 2 patients<br>● High single dose - 2 patients<br>● High* single dose - 4 patients<br>Each escalating dose cohort will be gated by an IDMC determination based on interim safety data.<br>**Or highest dose deemed tolerated based on interim analyses.* |
| CSF collection | <u>Goal</u>: target engagement and pharmacokinetic analyses<br><u>Timepoints relative to dose</u>: at enrollment, pre-dose, 6 weeks, 3 months, 6 months, 1 year | <u>Goal</u>: target engagement and pharmacokinetic analyses<br><u>Timepoints relative to dose</u>: at enrollment, pre-dose, 6 weeks, 3 months, 6 months, 1 year |
| Primary endpoint | Safety/ adverse events | Safety/ adverse events |
| Secondary endpoints | CSF PrP concentration | ● Time to death or initiation of life-extending measures (permanent intubation or ventilation)<br>● Time to drop 10 points on MRC Prion Disease Rating Scale<br>● CSF PrP concentration |
| Exploratory endpoints | Pathological biomarkers:<br>● CSF prion titer<br>● CSF total tau levels<br>● Plasma and CSF neurofilament light chain levels | Pathological biomarkers:<br>● CSF prion titer<br>● CSF total tau levels<br>● Plasma and CSF neurofilament light chain levels |

| | | |
|---|---|---|
| Long term follow-up | ● Safety<br>● Age of onset of genetic prion disease | ● Safety |
| Data analysis method | Intention to treat | Intention to treat |

##### 1.4. Questions

The specific questions we hereby pose to the agency are:

1. **[Clinical] In light of their distinct characteristics we propose to perform parallel trials in the two patient populations that stand to benefit from a PrP-lowering therapy: 1) symptomatic prion disease patients and 2) pre-symptomatic carriers of high penetrance pathogenic *PRNP* mutations. Does the Agency concur?**
2. **[Clinical] In both symptomatic and presymptomatic trials we propose to measure cerebrospinal fluid (CSF) prion protein levels as a pharmacodynamic biomarker of drug activity in the brain. Does the Agency concur?**
3. **[Preclinical] Our novel engineered AAV9-derived viral vector, BI-hTfR1v2, crosses the blood brain barrier by binding the human transferrin receptor. Because this interaction depends on the presence of a human-specific amino acid, BI-hTfR1v2 will not achieve pharmacologically relevant biodistribution in any common preclinical species such as wild-type mice, rats, dogs, minipigs or monkeys (8.3.1). For this reason, we propose that homozygous humanized *TFRC* knock-in mice are appropriate and sufficient as the single species for definitive biodistribution and toxicology studies. Does the Agency concur?**
4. **[Preclinical] Given the lack of homology between the human *PRNP* promoter and that of mouse, the planned human ZF-CHARMs will not have on-target activity in the *TFRC* mice in which we propose to conduct our definitive biodistribution and toxicology studies (8.3.2). We acknowledge that while this study will capture off-target effects of CHARM this study is not designed to capture on-target toxicity. However prion protein is known to be non-essential and we therefore do not expect on-target toxicity from PrP lowering. In order to assess on-target effects we propose to supplement our definitive biodistribution and toxicology studies with separate biodistribution and safety information collected in a non-GLP human PrP-lowering study (9.3.2.2) that will test the human ZF-CHARM candidate in humanized *PRNP* (huPrP) mice. Does the Agency concur?**
5. **[CMC] As a Potency Assay for release criteria, we propose to use RT-qPCR-based quantification of *PRNP* RNA in human HEK293T cells. Should an alternative assay be needed, we propose luminescence-based quantification of tagged endogenous PrP in human U251MG cells. Does the Agency concur?**

As patient-scientists living with a known high risk for genetic prion disease, we (Drs. Vallabh and Minikel) are highly motivated to seek the swift development of effective and safe PrP-lowering therapeutics.

The meeting will be a success if a path forward is identified for first-in-human treatment of prion disease patients and at-risk individuals with PRNP-CHARM-001.

#### 2. LIST OF FIGURES AND TABLES

##### LIST OF FIGURES

**Figure 1**. The pathway of prion disease
**Figure 2**. Published genetic and pharmacologic proofs-of-concept validating PrP lowering as a therapeutic hypothesis in prion disease.
**Figure 3**. Treatment time point is a key determinant of efficacy of PrP-lowering therapy.
**Figure 4**: CSF PrP levels are stable in presymptomatic prion disease mutation carriers.
**Figure 5**: CHARM: Coupled Histone tail for Autoinhibition Release of Methyltransferase.
**Figure 6**: CHARM modifies Prnp methylation profile, Prnp RNA brain RNA levels, and PrP brain protein levels in vivo.
**Figure 7**: CHARM dose-responsively lowers brain PrP, and extends survival of prion-infected mice
**Figure 8**: Towards a human-targeting PRNP-CHARM candidate.
**Figure 9**: BI-hTFR1v2 capsid is a second generation TfR1-binding capsid optimized for efficient CNS transduction.
**Figure 10**: BI-hTFR1 efficiently delivers genes to neurons in the CNS of TFRC KI mice.
**Figure 11**: BI-hTFR1 more efficiently transduces and crosses human brain endothelial cell monolayers in vitro by binding the apical domain of human TfR1.
**Figure 12**: The enhanced binding and transduction phenotype of BI-hTFR1 is specific to human TfR1
**Figure 13**: Binding of the BI-hTFR1v2 capsid to the apical domain of human TfR1 depends on the *** residue that is unique to humans.

##### LIST OF TABLES

**Table 1**: Evidence that PrP is central to prion disease pathophysiology.
**Table 2**: Utility of CSF PrP measurement in different prion disease patient populations.
**Table 3**: Mouse lines proposed for use in upcoming studies.
**Table 4**: Proposed study design for capsid bridging study (non GLP).
**Table 5**: Proposed study design for PRNP-CHARM-001 potency study (non GLP).
**Table 6**: Proposed study design for PRNP-CHARM-001 GLP pharmacology and toxicology study.
**Table 7**: Parameters of symptomatic and presymptomatic treatment populations.

#### 3. PRODUCT NAME

PRNP-CHARM-001

#### 4. CHEMICAL NAME AND STRUCTURE

PRNP-CHARM-001 is a recombinant adeno-associated virus (AAV)-based gene therapy comprised of a single-stranded AAV2 genome encoding an epigenetic editor (dubbed CHARM) targeted to the *PRNP* promoter by a zinc finger (ZF) binding domain and an engineered AAV9-derived capsid BI-hTFR1v2, that has been designed to enter the human CNS via its ability to bind to human Transferrin receptor (TfR1).

**PRNP-CHARM-001 consists of the following components:**

**Payload**: a single-stranded AAV2 genome containing AAV2 ITRs flanking the following payload:

- A mammalian promoter
- A cDNA payload with the following features:
- **CHARM effector domain**. CHARM stands for Coupled Histone tail for Autoinhibition Release of Methyltransferase[1]. This effector domain is a fusion protein comprised of:
    - A DNMT3L C-terminal domain (D3L), which recruits and stabilizes the endogenous DNMT3A and DNMT3A1 DNA methyltransferase enzymes
    - A linker domain
    - An unmethylated histone H3 tail, which stimulates DNMT3A activity
- **Zinc finger (ZF) DNA-binding domain.** Targeting to the *PRNP* promoter is achieved using a zinc finger protein domain. Note that the ZF domain is solely present for DNA targeting; it does not have, and never had, either an associated nuclease or catalytic activity, and neither DNA cutting nor base editing are involved in the drug's mechanism.
- A WPRE 3' UTR element for enhanced mRNA stability
- A polyadenylation sequence to terminate transcription

**Capsid: AAV9-derived vector BI-hTFR1v2**
BI-hTFR1v2 is a second generation AAV9-derived capsid built on the BI-hTFR1 capsid[2]. BI-hTFR1 contains a 7-mer peptide insertion in loop VIII of AAV9 and was engineered to bind to human transferrin receptor (TfR1, encoded by *TFRC*). The BI-hTFR1v2 capsid has additional substitution mutations that improve transduction in the CNS and reduce biodistribution to the liver in *TFRC* KI mice.

#### 5. PROPOSED INDICATION

Prion disease.

1. Patients symptomatic with prion disease
2. Presymptomatic carriers of highly penetrant pathogenic *PRNP* variants

#### 6. DOSAGE FORM AND ROUTE OF ADMINISTRATION

Sterile suspension for one-time intravenous infusion.

#### 7. PURPOSE AND FORMAT OF MEETING

The purpose of the meeting is to receive feedback on the questions listed below, which are critical for the advancement of PRNP-CHARM-001, a novel genetically targeted therapy with a well-established mechanism of action being developed for a rapidly fatal neurodegenerative disease with no standard of care. Above all, this meeting will be a success if a feasible path forward is identified for both the symptomatic and presymptomatic patient populations.

1. **[Clinical] In light of their distinct characteristics we propose to perform parallel trials in the two patient populations that stand to benefit from a PrP-lowering therapy: 1) symptomatic prion disease patients and 2) pre-symptomatic carriers of high penetrance pathogenic *PRNP* mutations. Does the Agency concur?**
2. **[Clinical] In both symptomatic and presymptomatic trials we propose to measure cerebrospinal fluid (CSF) prion protein levels as a pharmacodynamic biomarker of drug activity in the brain. Does the Agency concur?**
3. **[Preclinical] Our novel engineered AAV9-derived viral vector, BI-hTfR1v2, crosses the blood brain barrier by binding the human transferrin receptor. Because this interaction depends on the presence of a human-specific amino acid, BI-hTfR1v2 will not achieve pharmacologically relevant biodistribution in any common preclinical species such as wild-type mice, rats, dogs, minipigs or monkeys (8.3.1). For this reason, we propose that homozygous humanized *TFRC* knock-in mice are appropriate and sufficient as the single species for definitive biodistribution and toxicology studies, and that initial human doses could be selected based on these data. Does the Agency concur?**
4. **[Preclinical] Given the lack of homology between the human *PRNP* promoter and that of mouse, the planned human ZF-CHARMs will not have on-target activity in the *TFRC* mice in which we propose to conduct our definitive biodistribution and toxicology studies (8.3.2). We acknowledge that while this study will capture off-target effects of CHARM this study is not designed to capture on-target toxicity. However prion protein is known to be non-essential and we therefore do not expect on-target toxicity from PrP lowering. In order to assess on-target effects we propose to supplement our definitive biodistribution and toxicology studies with separate biodistribution and safety information collected in a non-GLP human PrP-lowering study (9.3.2.2) that will test the human ZF-CHARM candidate in humanized *PRNP* (huPrP) mice. Does the Agency concur?**
5. **[CMC] As a Potency Assay for release criteria, we propose to use RT-qPCR-based quantification of *PRNP* RNA in human HEK293T cells. Should an alternative assay be needed, we propose luminescence-based quantification of tagged endogenous PrP in human U251MG cells. Does the Agency concur?**

### 8. BACKGROUND ON PRNP-CHARM-001

#### 8.1. BACKGROUND ON PRION DISEASE AND THE THERAPEUTIC STRATEGY OF PRP LOWERING

##### *8.1.1. Prion disease is a fatal neurodegenerative disease with no existing cure caused by PrP*

Prion disease is a fatal and incurable neurodegenerative disease caused by prion protein (PrP)[3] that kills an estimated 400-500 people in the United States every year. PrP is encoded by a chromosomal gene, *PRNP*, and is expressed in the brains of all mammals, including humans. In prion disease, PrP undergoes post-translational misfolding into a self-replicating conformer known as a prion. Prions spread exponentially throughout the brain by recruiting other molecules of PrP, eventually accumulating to a level where they kill neurons. Subtypes of prion disease known historically by various names including Creutzfeldt-Jakob disease (CJD), fatal familial insomnia (FFI), and Gerstmann-Straussler-Scheinker (GSS) disease all share this same fundamental molecular mechanism.

First symptoms of prion disease can include a mix of cognitive, motor, and autonomic dysfunction but typically converge swiftly towards a rapidly progressive dementia[4]. Activities of daily living are swiftly lost as patients progress toward akinetic mutism. A typical patient will live only five months from first symptoms. 85% of cases are sporadic, apparently due to stochastic protein misfolding events, while the other 15% are genetic, due to gain-of-function mutations in *PRNP*[5]. Acquired prion disease caused by exposure to exogenous prions, for example through dietary routes or medical procedures, has thankfully become vanishingly rare, today accounting for <1% of cases[6]. There is no single point of origin for prions in the brain; as many regions can be first affected and prions readily spread across all regions, prion disease is best conceived of as a whole-brain disease. Near definitive pre-mortem diagnosis can be achieved through the combined deployment of RT-QuIC, a clinical assay that detects prion seeds in CSF[7], and targeted sequencing of *PRNP*.

To date there is no proven therapy for prion disease and no clinical standard of care. All available data from human genetics, biochemical approaches, laboratory animal models, and observations in wild and agricultural animals that are natural hosts of prion disease, are unanimous that PrP is the cause of prion disease. The therapeutic hypothesis of PrP lowering is backed by decades of genetic experiments in animal models and more recently by preclinical proof-of-concept studies in mice using antisense oligonucleotides (ASOs) (see below)[8–16]. A PrP-lowering ASO, ION717, has now progressed to a first-in-human trial under the sponsorship of Ionis Pharmaceuticals. However, ASO therapies are intrinsically limited in terms of potency, distribution, tolerability and the need for repeat intrathecal dosing. Therefore, the development of additional PrP-lowering therapeutics is urgently needed.

##### *8.1.2. PrP lowering is a well-validated therapeutic hypothesis in prion disease*

Decades of work from multiple orthogonal approaches including human genetics, histological and biochemical approaches, and animal models, all converge on PrP as *the* molecular cause of prion disease (Table 1). Briefly, infectious prions are composed of misfolded PrP[3,17,18]; PrP deposition is pathognomonic for prion disease neuropathology[19]; prion strains are encoded in PrP conformation[20–22]; PrP genetic variants cause genetic prion disease, impact sporadic prion disease risk, affect prion disease duration and presentation, govern species barriers and affect transmission risk in humans and other animals[23]. Because all prions, regardless of strain or species, are comprised of PrP, it has long been expected that depleting PrP substrate should be a universal therapeutic for prion disease regardless of the patients' etiology (sporadic, genetic, acquired) or subtype; recent studies have provided corroborating evidence that both pharmacologic and genetic lowering of PrP increase survival across prion strains[14].

***Table 1: Evidence that PrP is central to prion disease pathophysiology**. Reproduced from Vallabh 2020.*

| **Category** | **Evidence** |
|---|---|
| Biochemical | Prions, the infectious agent in prion disease, are composed of PrP[18].<br>Prion "strains" are encoded in distinct conformations of PrP[20–22].<br>Prion infectivity can be generated *in vitro* from purified PrP[24–27] . |
| Human genetics | All genetic prion disease families possess protein-altering variants in *PRNP*[28].<br>Certain *PRNP* missense variants confer protection against prion disease[29–31]. |
| Animal genetics | PrP is required for prion propagation[8].<br>PrP is required for prion neurotoxicity[10].<br>PrP dosage and incubation time are inversely correlated[9].<br>PrP amino acid sequence governs the "species barrier"[32]. |

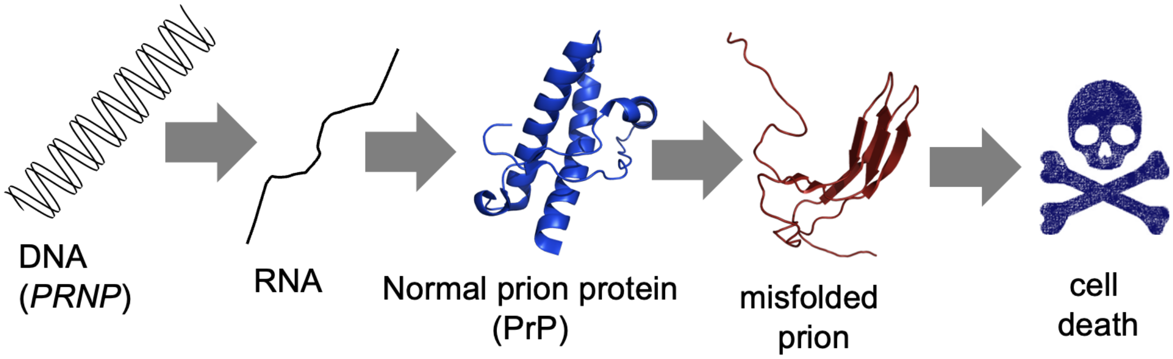


***Figure 1. The pathway of prion disease.** Adapted from Vallabh 2020.*

The advent of transgenic and knockout mice 30 years ago provided the first genetic evidence that PrP dosage governs the tempo of prion disease[9] (Figure 2A). These models are relevant because prion disease naturally afflicts not only humans but also diverse mammals including deer, elk, sheep, goats, cattle, camels, and mink. It is exceptionally well modeled in mice, where intracerebral inoculation of prions into naive wild type (WT) animals leads to fatal disease within 5–6 months, with all of the same pathological hallmarks seen in the human disease[33]. Heterozygous knockout mice survive prion infection more than twice as long as WT mice (Figure 2A), and homozygous knockouts are incapable of propagating prions or developing prion disease[8,16]. Conversely, disease is accelerated in transgenic mice overexpressing PrP[9]. This finding has been replicated in conditional genetic models, where lowering or abolishing PrP expression can respectively delay or prevent development of disease[11,12].

Benefit is directly responsive to PrP dosage. Chronic dosing with an ASO that reduces cortex *Prnp* RNA by 50%, beginning at an early treatment time point, triples survival of prion infected animals (Figure 2B). In a transient dosing paradigm, a low dose of ASO corresponding to 21% knockdown leads to a significant 15% increase in survival time, but progressively higher doses achieving deeper target engagement offer increasing survival benefit[14] (Figure 2C). ASO-mediated reduction of PrP in the brain is dose-dependently reflected in the lowering of CSF PrP in rats (Figure 2D), indicating that CSF is a relevant sampling compartment in which to monitor brain PrP levels[34].

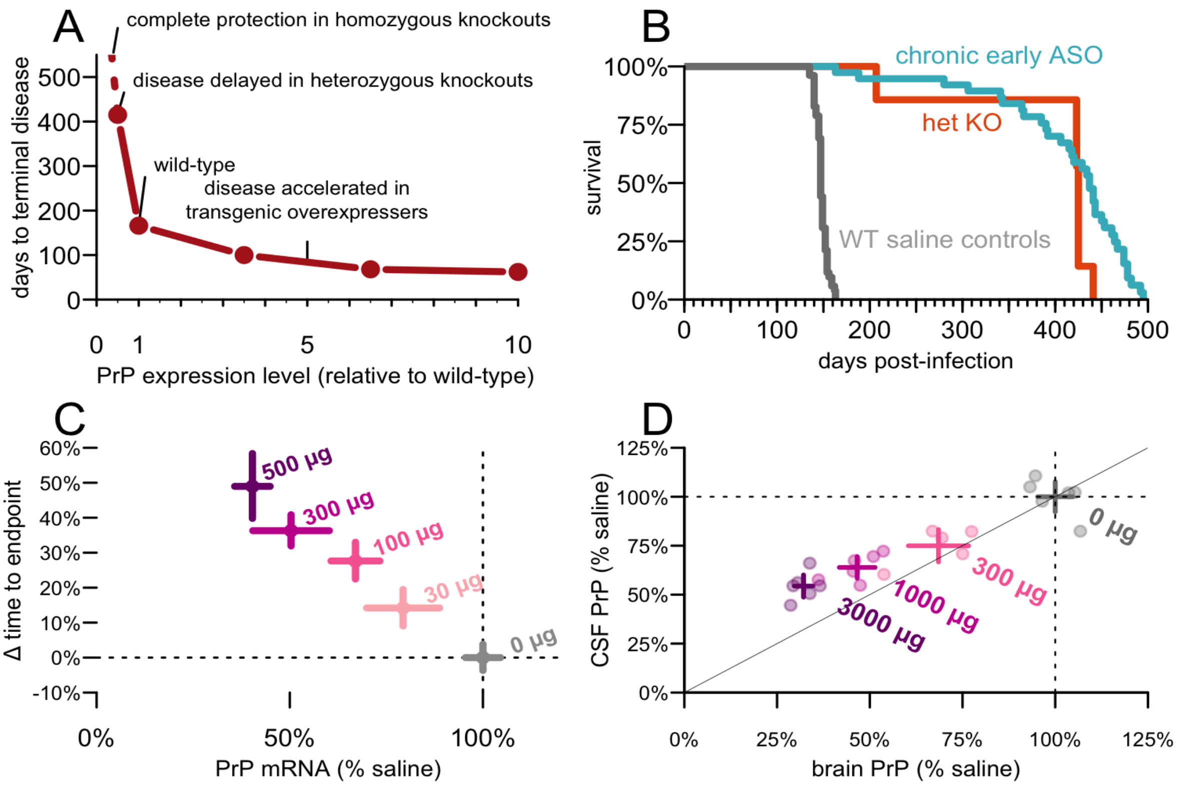


***Figure 2. Published genetic and pharmacologic proofs-of-concept validating PrP lowering as a therapeutic hypothesis in prion disease. A)****PrP expression level determines survival time following intracerebral prion inoculation in mice across wild-type, transgenic, and knockout lines.* ***B)*** *The benefit of chronic early ASO treatment on a symptomatic endpoint mirrors heterozygous knockout in mice.* ***C)*** *Following treatment with varying doses of PrP-lowering ASO, a continuous dose-response relationship emerges between brain PrP level and time to symptomatic endpoint in mice.* ***D)*** *CSF PrP mirrors brain PrP knockdown after ASO administration in rats. Panel A adapted from Fischer 1996[9] , B-C from Minikel 2020[14], D from Mortberg 2022[34].*

##### *8.1.3. Neurons are the cell type of interest for PrP-lowering therapies*

While prion disease affects all brain regions, prion toxicity is both specific to neurons and cell-autonomous, meaning that every neuron in which PrP is depleted is protected from disease. The progression of prion pathology halts following neuronally specific PrP depletion as shown in conditional knockout mice where *PRNP* is depleted in only neurons[11]. A comparison of mice restrictively expressing PrP under neuronal versus astrocytic promoters found that neuronal PrP alone supported typical prion disease, while mice expressing only astrocytic PrP failed to experience clinical symptoms following prion inoculation[35]. Though PrP knockout mouse brains neither propagate prions nor experience prion neurotoxicity following prion inoculation, a PrP-expressing graft introduced into the same knockout mouse brain will both amplify prions following prion challenge, and degenerate. The restriction to PrP-expressing tissue is precise; even knockout tissue adjacent to the graft shows no pathology[10]. These findings indicate that a therapy does not need to target every cell in the brain to rescue prion disease, but can be highly effective as long as it reduces PrP in the majority of neurons.

***PrP-lowering is beneficial at both presymptomatic and symptomatic timepoints – but the benefit differs in both degree and kind***

The most striking finding of almost one decade of work with PrP-lowering ASOs has been the importance of treatment time point to outcome.

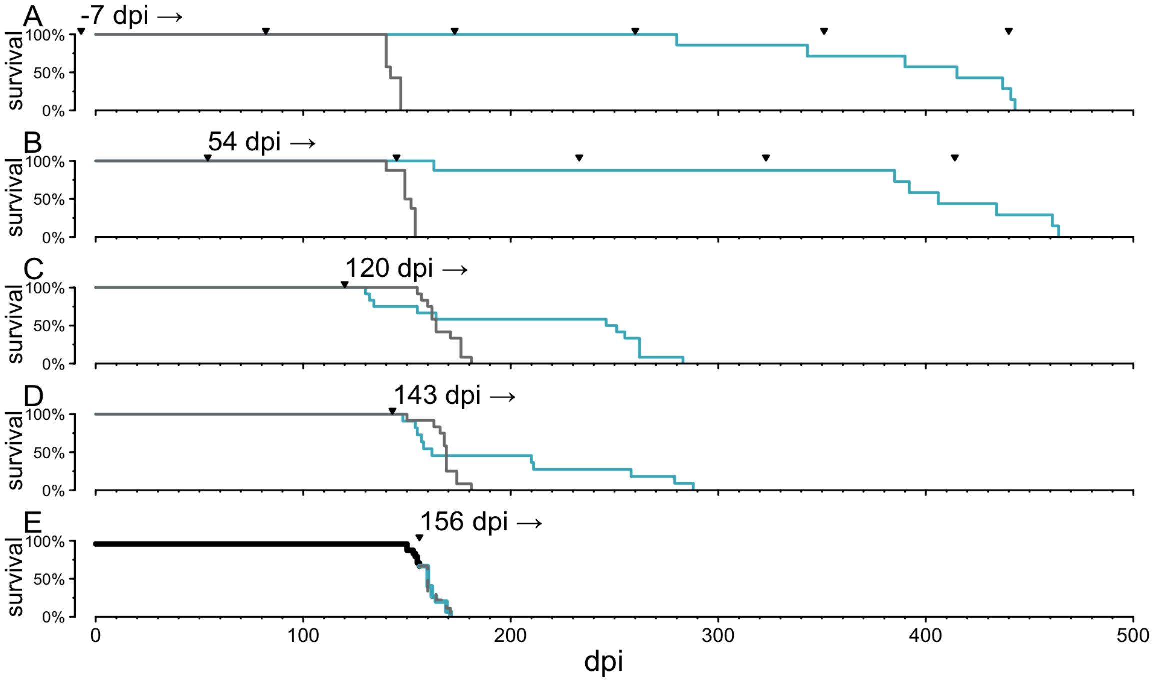


***Figure 3: Treatment time point is a key determinant of efficacy of PrP-lowering therapy.*** *dpi = days post inoculation. Adapted from Minikel 2020[14]. PrP-lowering ASO treatment conveys a different magnitude of survival benefit to prion-inoculated mice depending on the timepoint at which treatment is initiated, with earlier initiation corresponding to larger benefit and treatment at an advanced disease point failing to convey benefit. Mice received ASO treatment (cyan) or saline (gray) initiated prophylactically at 7 days before inoculation (A), just prior to detectable plasma neurofilament light chain (NfL) rise at 54 dpi (B), just prior to symptom onset in the presence of marked neuropathology at 120 dpi (C), after frank symptoms and weight loss in all*

*animals at 143 dpi (D), and after some animals have already reached endpoint (E). In the early treatment experiments (A, B), ASO was administered chronically every 90 days, and the endpoint was the presence of five prion disease symptoms. In the late treatment experiments (C–E), a single dose was administered and the endpoint was 20% weight loss. In (E) the black portion of the curve indicates survival before the dose was administered.*

Figure 3B shows that chronic administration of an ASO lowering PrP by ~50% can triple survival in prion-infected mice[14]. This result was obtained as long as therapy is initiated at any time from prior to infection up through 78 days post-infection (dpi), when animals are presymptomatic but neuropathological sequelae including astrogliosis and increase in plasma neurofilament (NfL) are emerging[14]. Notably, survival benefit in early-treated animals consisted mostly of an extension of healthy life.

In animals treated at the cusp of frank symptom onset, a single dose of ASO was still able to increase survival by 70%[14] (Figure 3C), however this timepoint does not correspond to a systematic clinical intervention timepoint in humans patients, the majority of whom are identified based on onset of frank symptoms. For the 15% of cases caused by a protein-altering *PRNP* variant, risk can be identified ahead of symptoms; however, these individuals do not reliably show even molecular signs of disease before onset of clinical symptoms, making presymptomatic treatment of carriers more analogous to the pre-pathological treatment paradigms shown in Fig 2B, 3A and 3B. Trial design considerations for this population are described in following sections.

If treatment was administered at a timepoint when all animals had measurably lost weight, the earliest consistent clinical sign of prion disease mice, it was possible to extend survival in a subset of the animals (Figure 3D), though they did not regain weight that they had lost, nor did their number of symptoms decline. Rather than extending healthy life, treatment at this symptomatic timepoint extended symptomatic decline of the animals. If treatment was administered at a timepoint corresponding to advanced disease (Figure 3E), when the first animals in the cohort had begun to reach prion endpoint, no animals benefited from treatment.

##### *8.1.4. PrP is dispensable for healthy life*

In contrast to its undisputed central role in prion disease, PrP appears dispensable for healthy life. PrP knockout mice[8], cows[36], and goats[37] are all healthy and viable and initially defied efforts to identify any knockout phenotype other than resistance to prion infection. It was eventually discovered that homozygous (but not heterozygous) PrP KO mice exhibit a myelin maintenance deficiency in peripheral nerves (but not in the CNS) leading to a mild peripheral neuropathy[38]. This is apparently due to a lack of signaling of an N-terminal PrP peptide to Adgrg6, a receptor expressed on peripheral Schwann cells but not in the CNS[39]. While this myelin phenotype is histologically evident in aged animals, it is phenotypically subtle. Identifying a difference in grip strength, rotarod performance, or hot plate response is difficult, and some aged (60 week) PrP KO mice but not others exhibit a small but significant difference from wild-type animals, apparently as a function of mouse strain[38]. The histological phenotype has been further replicated in PrP KO goats, where no outward phenotypic difference was reported[40]. We have identified older humans with naturally occurring heterozygous loss-of-function mutations in *PRNP* who appear to be healthy[5,41]. Moreover, such mutations occur in the population at exactly the frequency expected based on mutation rates; unlike the vast majority of genes in the human genome, there appears to be no purifying selection against loss-of-function variants in *PRNP*[41]. While Phase I studies should be conducted with the usual high level of care and monitoring, we have no reason to expect on-target toxicity from lowering PrP to the minimum achievable level. Consistent with this expectation, to date we have seen no evidence of an adverse phenotype associated with PrP lowering in mice receiving PrP-lowering ASO or CHARM treatment.

***8.1.5. Symptomatic and pre-symptomatic patients present opposing clinical opportunities and challenges.***

Prion disease is uniformly fatal. There are two distinct patient populations of interest: symptomatic patients and pre-symptomatic individuals at known high risk of fatal genetic prion disease.

**Symptomatic prion disease.** While all cases of prion disease trace to the misfolding of PrP, most cases appear to occur randomly, and are not presently identifiable before symptoms. Symptomatic patients, whether sporadic or genetic in etiology, typically follow a rapid progression. For the most common prion disease subtype, sporadic CJD, median survival is 5 months from first symptom[42]. During this time, cognitive and functional decline is extremely rapid: most patients lose the ability to complete all basic activities of living such as speaking, walking, eating, and using the toilet, within weeks[43]. Moreover, most patients lose considerable time just seeking a diagnosis — historically prion-specific diagnostic tests were not even ordered until ~3 months into the disease course[44] — though there is some evidence that diagnosis is improving in recent years[45]. Previous randomized trials, patients died a median of ~2 months from randomization[46,47], and many were profoundly impaired well before this time[46].

Given rapid progression, achieving clinical benefit in this population may be challenging. This is particularly true because PrP's in vivo half-life of 5 days[48,49] means that a PrP-lowering drug will require weeks to achieve full effect, during which time patients will continue to progress. As described above, we have observed in mouse studies that treatment at a symptomatic stage is less effective than treating before symptom onset (Figure 3). Certain factors such as Met/Val heterozygosity at *PRNP* codon 129 in sporadic disease, or the pathogenic P102L *PRNP* that causes genetic prion disease tend to predispose to a slightly more slowly progressive disease course[43]; to best position a symptomatic trial for success, such factors could be assessed at enrollment along with current symptom status.

**Pre-symptomatic at-risk carriers.** Roughly 15% of cases of prion disease trace to protein-coding variants in *PRNP*. Carriers of such variants can be identified years or decades in advance of symptoms, offering an opportunity to treat before the rapid symptomatic course begins. In recent years, case-control datasets have enabled penetrance estimates for *PRNP* variants to be refined, and individuals with ~90% or higher lifetime risk of fatal prion disease can now be reliably identified[5]. Three pathogenic variants – E200K, D178N and P102L – comprise 85% of high-penetrance genetic prion disease cases[5].

Clinical trials in presymptomatic at-risk individuals offer the opportunity to test PRNP-CHARM-001 not just for its ability to intercept and prolong an ongoing disease process, but for its ability to preserve healthy life. Unlike symptomatic patients, presymptomatic carriers will reliably live to see the drug to reach full target engagement, increasing the potential for benefit. Natural history data show that cross-sectionally, in most carriers available for recruitment there is no sign of a disease process underway nor that initial prion formation has occurred. Only in the presymptomatic population does a PrP-lowering drug have the opportunity not only to slow prion propagation and neurotoxicity, but to slow or prevent the initial prion formation event, again enhancing the prospect for benefit.

For the purposes of initial study endpoints, pre-symptomatic at-risk individuals provide an unconfounded background against which to assess both safety and target engagement of PRNP-CHARM-001. Characterization of presymptomatic at-risk mutation carriers shows their health status to be normal[50], allowing for more straightforward assessment of safety than in symptomatic patients, where drug-related events must be differentiated from rapidly evolving and multi-faceted symptom progression.

*CSF PrP can serve as a pharmacokinetic biomarker in the presymptomatic population*

PrP is abundant in CSF and readily quantifiable by ELISA. Longitudinal CSF sampling has shown that CSF PrP levels are stable in presymptomatic genetic prion disease carriers across genotypes (Figure 4), just as they are in healthy controls. This known stability contrasts with symptomatic patients, in whom the kinetics of PrP accumulation into plaques and impact on soluble PrP levels in CSF over time are not clear[51] (Table 2). Meanwhile rat ASO treatment studies show that reduction of prion protein in the brain is dose-dependently mirrored in CSF[34] (Figure 2D), demonstrating the relevance of CSF as a sampling compartment.

***Table 2: Utility of CSF PrP measurement in different prion disease patient populations.***

| **Disease stage** | **Etiology** | **PrP ELISA useful for target engagement?** | **Concerns** |
|---|---|---|---|
| Pre-symptomatic | Genetic | Meaningful at baseline and post-treatment | none |
| Symptomatic | Genetic | Utility not established | Declines over the symptomatic course of disease; detailed baseline trajectory not available. |
| Symptomatic | Sporadic | Utility not established | Declines over the symptomatic course of disease; detailed baseline trajectory not available. |

In addition to providing a readout of brain target engagement, CSF PrP may have additional significance in the presymptomatic population. Given that PrP is well established as the single causal molecule in prion disease and as a dose-dependent driver of the disease process, reduction in CSF PrP levels has also been proposed to be predictive of potential clinical benefit analogous to viral load in HIV/AIDS[15]. Indeed, as further described below, we have previously discussed with CDER the potential for CSF PrP lowering to serve as a surrogate endpoint in preventive genetic prion disease trials of PrP-lowering oligonucleotide drugs[15].

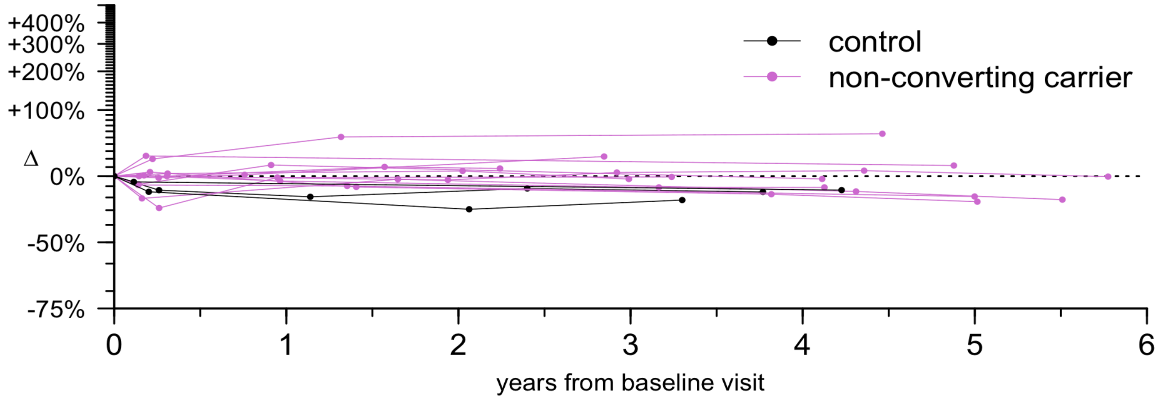


***Figure 4: CSF PrP levels are stable in presymptomatic prion disease mutation carriers.*** *Longitudinal CSF PrP levels in presymptomatic genetic prion disease mutation carriers across a range of genotypes, as well as non-mutation controls, show a mean test-retest CV of 10%, demonstrating assay reliability and stable PrP levels in presymptomatic genetic carriers. CSF was drawn roughly annually over up to six years. Adapted from Vallabh 2024.*

***8.1.6. Key points for treatment protocols***

Proposed clinical study protocols for both the symptomatic prion disease population and presymptomatic high-risk population are detailed in Sponsor Position 2 (Tables 5 and 6). Note that we do not propose gating eligibility of presymptomatic high-risk individuals on the presence of a molecular prodrome, for two reasons. First, such a prodrome is both brief and inconsistent in prion disease if detectable at all[50,52], making this recruitment strategy numerically infeasible. Second, all preclinical data from PrP-lowering therapies suggests that once prodromal signs of disease such as neuronal damage markers are detectable, the prospect for therapeutic benefit is already compromised compared to fully preventive treatment (Fig 3).

The key points of both proposals are summarized below.

*Proposal for symptomatic treatment:*

- Recruit individuals with early prion disease symptoms based on the Medical Research Council (MRC) prion disease rating scale. Note that while these individuals will all be symptomatic, their etiology will vary; patients in the symptomatic stage of both sporadic and genetic prion disease will be eligible.
- Treat with PRNP-CHARM-001.
- Primary endpoint: safety.
- Secondary endpoint: CSF PrP levels. Monitor CSF PrP levels at the following timepoints for target engagement and pharmacokinetic analyses: enrollment, pre-

dose, six weeks post-treatment, three months post-treatment, six months post-treatment, and 1 year post-treatment.

- Exploratory endpoints: pathological biomarkers. Measure CSF prion titers, CSF total tau levels, and CSF and plasma NfL levels at the same timepoints listed above.
- Follow treated individuals long term (already standard for one-time gene therapies). Continue to monitor CSF PrP to assess real-world treatment durability. While this trial is not powered for a survival or pathological endpoint, on an exploratory basis survival or time to intubation/ventilation, pathological biomarker trajectories and MRC rating scale trajectories can be compared to natural history controls.

*Proposal for pre-symptomatic treatment:*

- Recruit individuals with highly penetrant *PRNP* pathogenic variants, based on genotype and age, informed by penetrance estimates and age of onset datasets in hand.
- Treat preventively with PRNP-CHARM-001.
- Primary endpoint: safety.
- Secondary endpoint: Cerebrospinal fluid PrP levels. Monitor CSF PrP levels at the following timepoints for target engagement and pharmacokinetic analyses: enrollment, pre-dose, six weeks post-treatment, three months post-treatment, six months post-treatment, and 1 year post-treatment.
- Exploratory endpoints: pathological biomarkers. Measure CSF prion titers, CSF total tau levels, and CSF and plasma NfL levels at the same timepoints listed above. All are expected to be negative in controls, but should be monitored
- Follow treated individuals long term (already standard for one-time gene therapies). Continue to monitor CSF PrP to assess real-world treatment durability. While this trial is not powered for a survival or pathological endpoint, on an exploratory basis age of onset and pathological biomarker trajectories can be compared to genotype-matched natural history controls[53].

##### ***8.1.7. Relevant history with CDER***

We have previously interacted with CDER under PIND 141250 about PrP-lowering ASOs. Of particular note, on November 11, 2017, we met with twenty-five CDER scientists in the context of a Critical Path Innovation Meeting to discuss preventive clinical strategies for the advancement of PrP-lowering therapeutics. The notes from this meeting are attached (Appendix 1), and the white paper we submitted as prereading material has since been adapted for publication[15]. The need for preventive treatment in prion disease was recognized in this meeting, and the discussion focused on enabling steps. Feedback received at this meeting has motivated many of our efforts since, including:

- Validation that PrP lowering is protective against prion disease in additional in vivo paradigms: rats, hamsters, humanized mice infected with human prions, and a spontaneous mouse model of prion disease.
- Creation of an online patient registry (www.prionregistry.org)

- Expansion of natural history efforts to demonstrate stability of CSF PrP in individuals of various *PRNP* genotypes, over the timespan of years (Figure 4)
- Validation that treatment-mediated CNS reduction of PrP is dose-dependently reflected in CSF PrP levels (Figure 2D).

#### 8.2. BACKGROUND ON TARGETED EPIGENETIC SILENCING USING CHARM

In early 2023, our team at the Broad Institute partnered with Dr. Jonathan Weissman's team at the Whitehead Institute and Dr. Ben Deverman's team at the Broad Institute to develop an AAV-mediated, PRNP-targeting CHARM therapeutic for prion disease. We applied for and were awarded an NIH grant (NINDS 5U19NS132315) to support preclinical proof-of-concept and IND-enabling studies.

##### *8.2.1. Epigenetic editing as a therapeutic intervention for prion disease*

Epigenetic transcriptional silencing represents an attractive approach for eliminating expression of pathogenic proteins like PrP without the need to mutate the underlying DNA sequence[54–59]. Prion disease is an excellent candidate for this approach, since simply decreasing PrP expression will have a therapeutic effect[15] and the human PRNP promoter contains a large annotated CpG island to serve as a substrate for DNA methylation.

Permanent silencing can be achieved through targeted DNA methylation by the recruitment of the constitutively active catalytic domain (D3A) of the de novo DNA methyltransferase enzyme DNMT3A[60–62] along with the C-terminal domain (D3L) of its cofactor DNMT3L[63–65]. DNA methylation at cytosine-guanine dinucleotide (CpG) sites, producing 5-methyl-CpGs (5mCpGs), is mitotically inherited and contributes to transcriptional silencing directly by blocking transcription factor binding and indirectly by recruiting methyl-CpG-binding factors that induce heterochromatin[66]. The Weissman lab previously demonstrated that these domains, with the addition of a repressive KRAB domain, could be fused to a nuclease-deficient S. pyogenes Cas9 yielding a CRISPR-based editor for programmable, heritable gene silencing termed[67].

CHARM (Coupled Histone tail for Autoinhibition Release of Methyltransferase) is a more recently developed, related approach with specific advantages for treating prion disease. Two key improvements promote both tolerability and deliverability to the central nervous system, which is constrained by the 4.7 kb packaging limit of AAV[1].

***i. Use of a zinc finger DNA targeting domain.*** Zinc finger proteins (ZFPs) are ubiquitous DNA-binding proteins in eukaryotes[68] whose modular nature has enabled programming for specific genome targeting[69–73]. Previous work has demonstrated that engineered ZFPs fused to chromatin-modifying domains can successfully modulate target gene transcription in vivo[74–77]. As a DNA targeting module, ZFPs offer the advantage of their compact size which, at roughly an order of magnitude smaller than that of SpCas9, makes them suitable for delivery via AAV (Fig. 5B) ZFPs, which lack bacterial epitopes[78] also offer reduced risk of immunogenicity compared to SpCas9, of which a large proportion of the human population already has an immune memory[79–81].

***ii. Recruitment of endogenous methyltransferases.***
A further departure from the CRISPR-off model is that rather than risking potentially toxic overexpression of catalytically active DNA methyltransferase by expressing it as part of the therapeutic cargo, CHARM recruits endogenous DNMT3A1, the dominant DNA methyltransferase in somatic tissue. To circumvent the need to overexpress the D3A catalytic methyltransferase domain, CHARM incorporates the catalytically inactive D3L domain, which recruits the full-length endogenous enzyme to the target site[82,83], and an unmethylated H3 histone tail, which stimulates its activity[84–86].

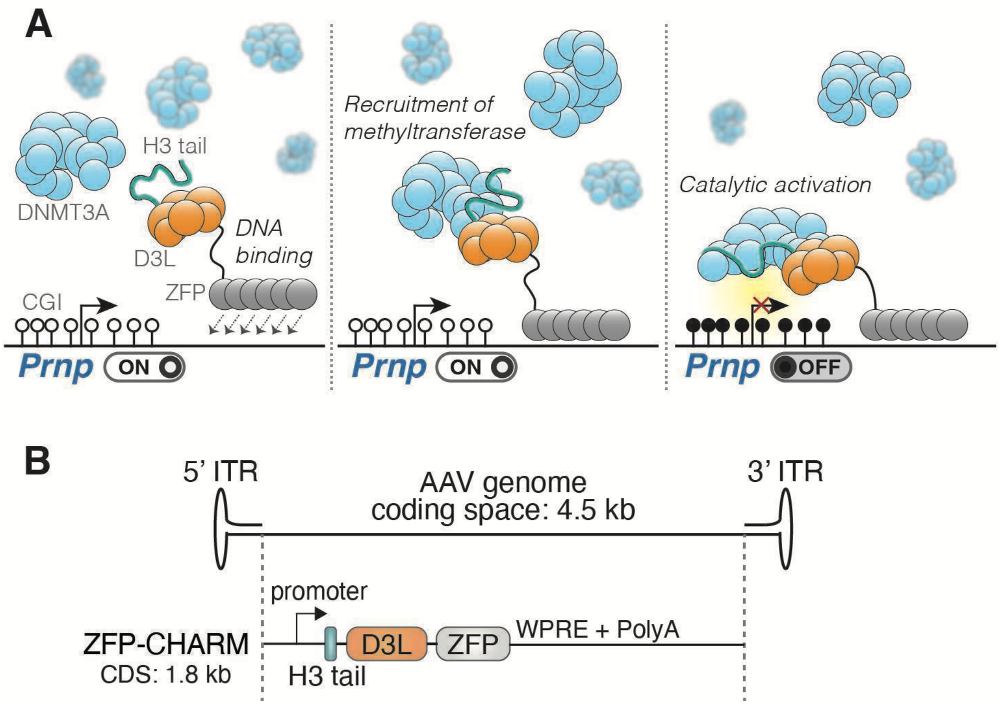


***Figure 5. CHARM: Coupled Histone tail for Autoinhibition Release of Methyltransferase.*** A) Cartoon depiction of endogenous DNMT3A recruitment and activation by the CHARM system. The D3L domain interfaces with the DNMT3A methyltransferase domain to facilitate recruitment, and the histone H3 tail binds the DNMT3A ADD domain to stimulate catalytic activity. B) Size comparison of CHARM to available coding space in a single-stranded AAV2 genome.

##### *8.2.2. CHARM can be used to silence PRNP*

To establish proof of concept, CHARM constructs targeting the mouse *Prnp* locus were tested in cells as described[1], then advanced to in vivo studies. For mouse experiments, constructs were packaged in the PHP.eB vector known to transduce mouse neurons following IV dosing. Unless otherwise specified, all mice in our in vivo experiments received 1.5e13 vg/kg CHARM delivered intravenously by retro-orbital injection, and were sacrificed 6 weeks post injection. Control animals received  no injection (n=5 per group). As prion disease is a whole brain disease,

Prnp RNA and PrP protein were quantified by RT-qPCR and ELISA, respectively, in brain whole hemisphere homogenates. Prnp RNA levels were reduced by 82 (82.1)% and PrP protein levels by 68 (68.3)%. These are, respectively, the largest treatment-mediated reductions in brain PrP ever observed in our studies or reported by any group.

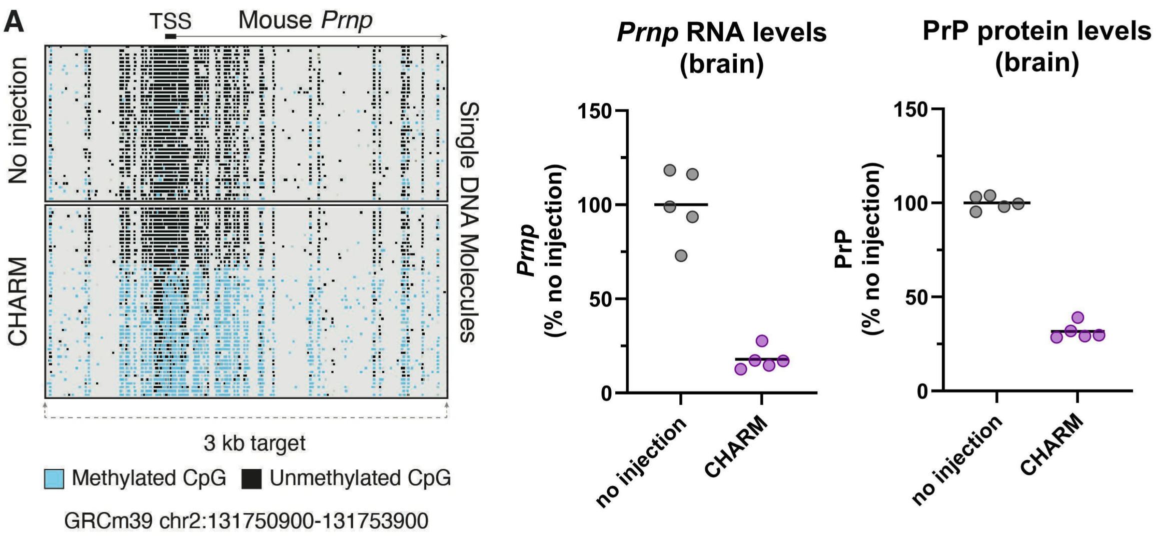


***Figure 6. CHARM modifies Prnp methylation profile, Prnp RNA brain RNA levels, and PrP brain protein levels in vivo.*** N=5 wild-type C57BL/6N mice per group were dosed with 1.5e13 vg/kg AAV-PHP.eB containing PRNP-CHARM, or received no injection at 5-8 weeks of age. A) DNA methylation was assessed by targeted nanopore long-read sequencing in genomic DNA extracted from whole brain hemispheres 6 weeks post dose. Whole brain hemispheres were analyzed for B) Prnp RNA by RT-qPCR (housekeeping gene TBP) and C) PrP protein levels by ELISA at 6 weeks post-dose.

Next, we explored multi-dose and longer term in vivo paradigms. Across a four-fold dose range, we observed dose responsive reductions in brain Prnp RNA and PrP protein (Fig 7A-B). We further observed no significant difference in target reduction at 6 weeks and 12 weeks, and 6 months post dose, consistent with the long-term activity expected of the CHARM mechanism (Fig 7C). An additional 12-month durability study is ongoing.

We also performed a survival study in the best-established model of prion disease, wild-type (C57BL/6) mice intracerebrally inoculated with the RML strain of mouse prions. These animals received no injection or CHARM (n=10 treated, n=15 controls) at 120 days post inoculation (dpi), a timepoint that corresponds to the cusp of symptom onset. Untreated animals reached prion disease terminal disease endpoint at median 168 dpi while treated animals survived to median 307 dpi, resulting in a mean extension of survival of 83%.

**Fig 7. CHARM dose-responsively lowers brain PrP, and extends survival of prion-infected mice. A,B)**

Finally, having established proof of concept by targeting mouse *Prnp*, we are in the early stages of screening human candidates for human *PRNP*. Given the lack of homology between the mouse and human *PRNP* promoter, our ZF-CHARMs are species-specific. However the CpG content of the human *PRNP* promoter is similar to that of mouse (Figure 8A-B), suggesting equal targetability a priori. Indeed, methylation profiling and potency studies in human cells (Figure 8C) have nominated top performing human PRNP-CHARM constructs with in vivo potency in *PRNP* humanized mice comparable to the mouse tool compounds described above. Studies to select a lead molecule for advancement are ongoing.

***Figure 8. Towards a human-targeting PRNP-CHARM candidate.***

#### 8.3. BACKGROUND ON DELIVERY VIA A SYSTEMICALLY DELIVERED BLOOD-BRAIN BARRIER-CROSSING AAV CAPSID

An effective prion disease therapy requires efficient PrP lowering in neurons throughout the entire CNS, ideally via a systemically delivered BBB-crossing vector that can reach all regions of the CNS via extensive capillary networks. Unfortunately, AAV9, the natural serotype commonl used for CNS gene therapies, does not efficiently cross the adult BBB and transduce neurons. Therefore PRNP-CHARM-001 will use a novel capsid, BI-hTFR1v2 that has enhanced BBB crossing activity even at low to moderate systemic doses.

##### *8.3.1. The BI-hTFR1v2 capsid enables dramatically improved CNS delivery compared toAAV9*

PRNP-CHARM-001 uses an AAV9-derived capsid BI-hTFR1v2, which was specifically designed for systemically delivered human CNS gene therapy applications. BI-hTFR1v2 is a second generation variant of the recently developed capsid BI-hTFR1[2] (Huang and Chan *et al Science* 2024) that crosses the BBB using receptor-mediated transcytosis via interactions with the human Transferrin receptor (TfR1) (Figure 9).

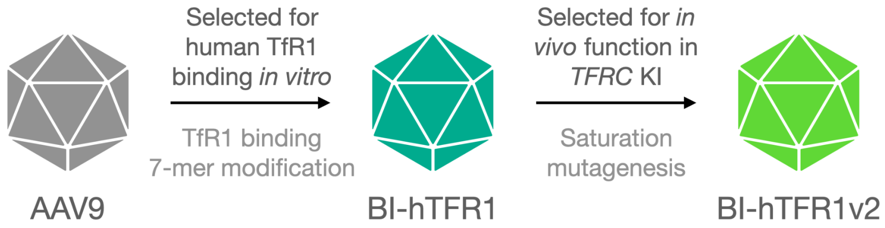


**Figure 9. BI-hTFR1v2 capsid is a second generation TfR1-binding capsid optimized for efficient CNS transduction.** BI-hTFR1 and BI-hTFR1v2 contain the same 7-mer insertion required for binding human TfR1. BI-hTFR1v2 contains additional mutations outside of the 7-mer insertion that reduce liver biodistribution and improve the CNS tropism in humanized *TFRC* knock-in (KI) mice.

<u>Key strengths of BI-hTFR1v2</u>:

(1) BI-hTFR1 and BI-hTFR1v2 are based on AAV9, the best studied natural AAV variant for systemic CNS delivery applications, which has been used successfully to treat spinal muscular atrophy (SMA) due to its ability to transduce spinal motor neurons early in life [Byrne 2019]. Compared to AAV9, the enhanced tropism of BI-hTFR1 is CNS-specific with no detected increase in targeting to the liver or other assessed organs. BI-hTFR1 showed a selective increase in biodistribution to and transduction of the brain and spinal cord (Fig. 10B) and transduced the majority of neurons across multiple brain regions in *TFRC* KI but not WT mice when delivered at a dose of $2.5 \times 10^{13}$ vg/kg (Fig. 10C–E).

(2) Our second generation BI-hTFR1v2 capsid is capable of even greater CNS neuron transduction as compared to BI-hTFR1. In adult TFRC KI mice, BI-hTFR1v2 can transduce more than 50% of $NeuN^+$ cortical neurons when IV administered at the low systemic dose of $2.5 \times 10^{12}$ vg/kg (~50 times lower than clinical precedents with AAV9). BI-hTFR1v2 also exhibits a reduced biodistribution and transduction in the liver relative to BI-hTFR1 and AAV9.

(3) BI-hTFR1 utilizes a well established, <u>human-relevant</u> mechanism-of-action to cross the BBB: the engagement of the apical domain of human TfR1 (Figure 11A and B). BI-hTFR1v2 shares an identical 7-mer targeting peptide with BI-hTFR1 and thereby engages TfR1 at the same site within the apical domain. TfR1 mediates constitutive, ligand-independent receptor-mediated transcytosis (RMT) across the CNS vasculature[87–90] and boasts a track record as a target to increase the delivery of biologics into the CNS of mice[91–93], nonhuman primates (NHPs)[94–96], and humans as investigational therapies[97] and as an approved antibody-based therapeutic for Mucopolysaccharidosis type II[98]. Protein shuttles designed to bind human TfR1 to facilitate RMT are showing evidence of CNS entry[99] and <u>efficacy in humans at doses predicted from *TFRC* KI mouse studies</u>[98,100,101].

(4) BI-hTFR1v2 binds TfR1 in the presence of its natural ligand, holo-Tf, which is critical because AAVs cannot be administered at doses capable of competing with Tf for TfR1 binding[2].

(5) BI-hTFR1 can be consistently produced and purified using commonly used affinity resins with yields comparable to AAV9[2]. BI-hTFR1v2 is likewise compatible with commonly used affinity resins. Our approach did not require the introduction or conjugation of bulky protein domains. We do not anticipate additional complexities in the manufacturing of BI-hTFR1v2 compared to AAV9, one of the top producing natural AAVs.

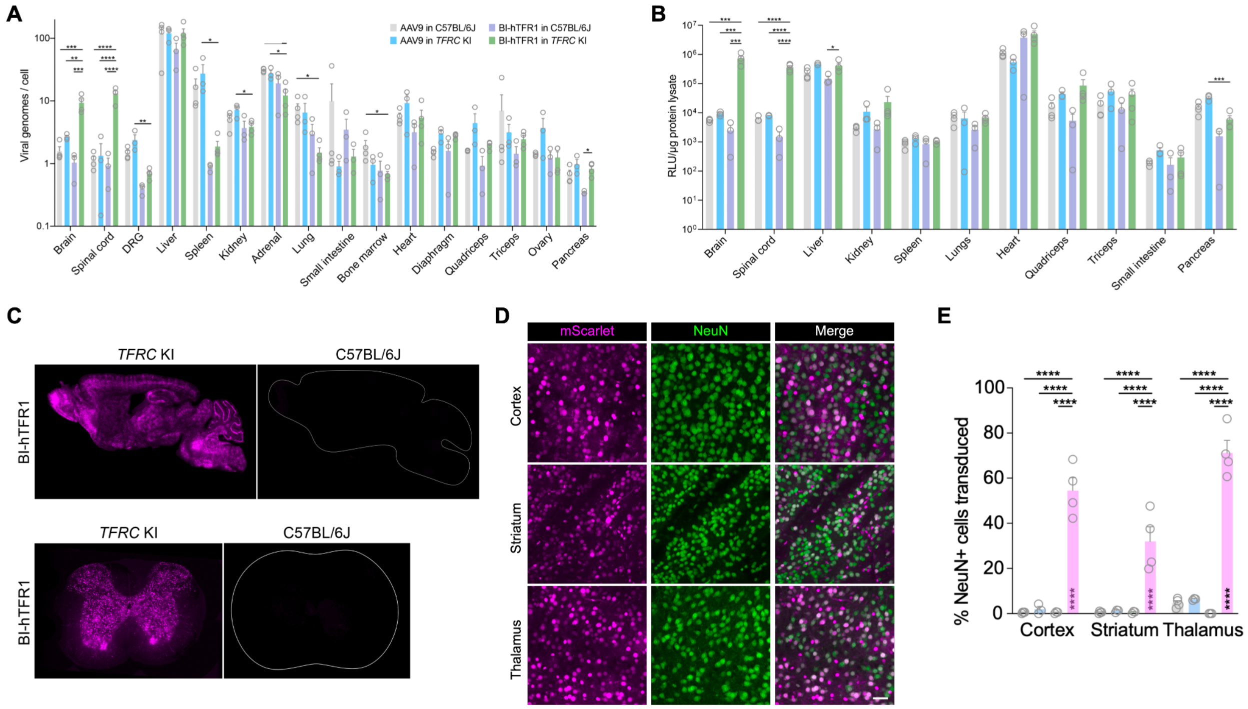


**Figure 10. BI-hTFR1 efficiently delivers genes to neurons in the CNS of *TFRC* KI mice.** In *TFRC* KI mice, mouse *Tfrc* exons 4–19 encoding the extracellular region of TfR1 have been replaced by those of human *TFRC*. (**A, B**) BI-hTFR1 or AAV9 encoding CAG-NLS-mScarlet-P2A-Luc-WPRE-SV40pA were intravenously injected into adult female C57BL/6J or *TFRC* KI mice at 5 x $10^{11}$ vg/mouse (approximately $2.5 \times 10^{13}$ vg/kg). The biodistribution reported as vector genomes per mouse genome (**A**) and transduction based on luciferase relative light units (RLUs/ug protein) in lysates (**B**) are shown at three weeks post-injection (two-way ANOVA using BI-hTFR1 in *TFRC* KI mice as the main comparison group with Bonferroni multiple comparison correction: ****, ***, **, and * indicate $p \leq 0.0001$, $\leq 0.001$, $\leq 0.01$, and $\leq 0.05$, respectively; each data point represents an individual mouse, error bars indicate ± SEM). (**C**) Representative brain and spinal cord images from the indicated groups of mice are shown. (**D**) Images show the colocalization of mScarlet reporter expression with NeuN+ neurons in the indicated brain regions. (**E**) The percentages of $NeuN^{+}$ neurons transduced is shown (two-way ANOVA using BI-hTFR1 in *TFRC* KI mice as the main comparison group with Bonferroni multiple comparison correction: **** indicates $p \leq 0.0001$; each data point represents an individual mouse, error bars indicate ± SEM).

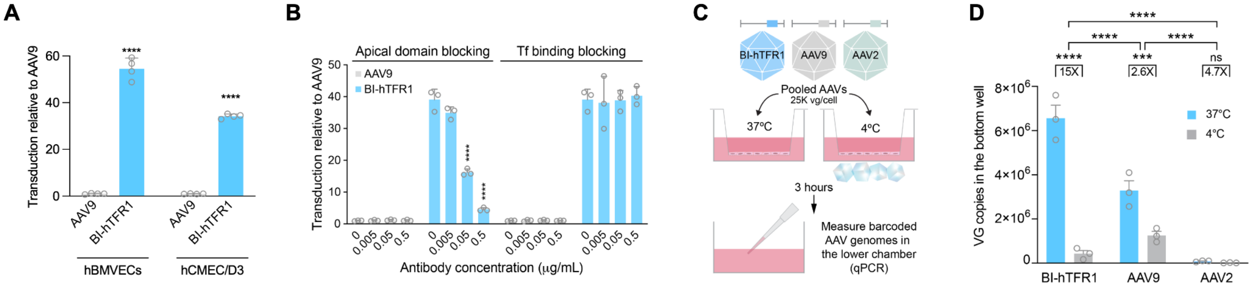


**Figure 11. BI-hTFR1 more efficiently transduces and crosses human brain endothelial cell monolayers *in vitro* by binding the apical domain of human TfR1.** (**A**) BI-TFR1 more efficiently transduces hBMVEC and hCMEC/D3 cells as compared with AAV9 (luciferase reporter assay). (**B**) Transduction (luciferase activity) of hCMEC/D3 cells incubated with 3 x $10^{8}$ vector genomes (vg)/mL of the indicated AAV and specified concentrations of OKT9 or AF2474 antibody is shown (two-way ANOVA using the no antibody control for each condition as the main comparison group with Bonferroni

multiple comparison correction: **** indicates $p \leq 0.0001$; n = 3 replicates, error bars indicate ± SEM). In (A) and (B), values are normalized to AAV9 (reported as fold change) in each cell line. (**C**) The transwell BBB model experimental design is shown. (**D**) More BI-hTFR1 is actively (37C vs 4C) transported from the apical (top) to basolateral (bottom) chamber as compared to AAV9 and AAV2. Vector genomes in the bottom chamber were quantified by qPCR (two-way ANOVA with Bonferroni multiple comparison correction: **** and *** indicate $p \leq 0.0001$ and $\leq 0.001$ respectively; n = 3 transwell replicates, error bars indicate ± SEM).

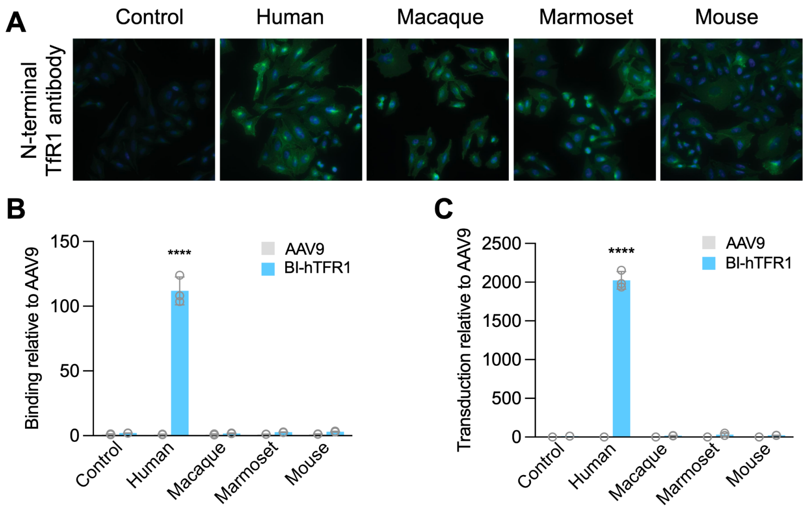


**Figure 12. The enhanced binding and transduction phenotype of BI-hTFR1 is specific to human TfR1.** (**A**) CHO cells were generated that stably express *TFRC* from the indicated species. Images show immunostaining using an anti-TfR1 antibody that recognizes a common N-terminal epitope. BI-hTFR1 associates with (**B**) and transduces (**C**) cells expressing human TfR1, but not cells expressing macaque, marmoset, or mouse TfR1.

##### *8.3.2. The CNS tropism of BI-hTFR1v2 is specific to human TfR1*

BI-hTFR1 does not exhibit enhanced transduction of cells expressing TfR1 from macaques, marmosets, or mice (Figure 12). To understand the species specificity in more detail, we mapped the human-specific amino acids required for BI-hTFR1v2 binding to TfR1. CHO cells modified to express human TfR1 were efficiently transduced by BI-hTFR1v2 (Figure 13). In contrast, in cells that express human TfR1 with *** and *** (introducing the macaque amino acids at this position), BI-hTFR1v2 exhibited no increased transduction phenotype as compared with AAV9. Similar findings were obtained when *** alone was introduced (Figure 13F). Therefore, the human specific amino acid *** is required for BI-hTFR1v2's ability to bind human TfR1. Fitting with these data, replacing *** with a *** residue in macaque TfR1 was sufficient to render cells expressing this mutant form of macaque TfR1 permissive to transduction by BI-hTFR1v2. Notably, the *** residue is only found in human TfR1 and not any other common preclinical or domesticated species (Figure 13H). Based on these data, we conclude that macaques are not a physiologically relevant species for assessing the biodistribution and toxicity of BI-hTFR1v2 and PRNP-CHARM-001.

**Figure 13. Binding of the BI-hTFR1v2 capsid to the apical domain of human TfR1 depends on the ***residue that is unique to humans.**

We expect that IV dosing of BI-hTFR1v2 in *TFRC* KI mice will be representative of IV dosing in humans based on the following: (1) The mechanism of transport of the capsid across the BBB (via binding TfR1) is conserved across species. (2) TfR1 protein levels in mouse and human microvasculature are similar. (3) Receptor mediated transcytosis across the vasculature scales with brain size because brain capillaries in both mice and humans are typically no more than 40 µm apart and most neurons are therefore within 20 µm of a capillary[102]. Together, these findings support the expectation that AAV vg/kg dosing in humanized *TFRC* KI animals will be consistent with the human vg/kg dose.

#### 9. STATUS OF PRODUCT DEVELOPMENT

PRNP-CHARM-001 is being developed as an academic, investigator-initiated project with NIH and philanthropic funding. There is no industry sponsor, and there are no plans to commercialize.

##### 9.1. GENERAL INVESTIGATIONAL PLAN

Our approach is non-allele-specific and in principle, our drug candidate will be capable of lowering PrP in any and all patients diagnosed with or at risk for prion disease. The potential patient population for this approach is therefore hundreds of patients in the U.S. and low thousands worldwide.

A proposed study protocol will be filed with the IND submission. Our draft study parameters for both a first symptomatic and first presymptomatic clinical study are outlined in the Executive Summary.

##### 9.2. CHEMISTRY, MANUFACTURING AND CONTROL (CMC) DATA

**Potency assay**

The mechanism of action for our therapeutic is reduction of PrP expression via methylation of the *PRNP* gene promoter. The transgene packaged in our vector leads to expression of the ZF-CHARM epigenetic editor, resulting in DNA methylation at the *PRNP* gene promoter and attendant decreases in *PRNP* RNA and PrP protein expression. We are contacting contract research organizations to explore implementing a *PRNP* expression-based potency assay.

As our primary strategy, we propose to quantify *PRNP* RNA levels in HEK293T by RT-qPCR. CHARM-based reductions in PrP protein dose dependently tracks with reduction in PRNP mRNA (Figure 7A, B). In addition, the PRNP promoter has been shown to be silencable by ZFP-CHARM in HEK293T cells[1] (see Figure S5). We will transduce HEK293T cells with 5 serial dilutions of the drug product based on a standard number of vector genomes per cell (vg/cell) and perform qRT-PCR to measure expression of *PRNP* RNA following AAV transduction. Importantly, HEK293T cells robustly express *PRNP* at the RNA level; they also express *TFRC* and are efficiently transduced by BI-TfR1v2. This assay will determine the correlation between AAV concentration and qPCR PrP detection in cell lysates, including comparison to the signal obtained for an 8-point standard curve based on a plasmid dilution series.

As an alternative strategy, we propose to quantify PrP expression at the protein level using a tagged U251MG human glioblastoma derived cell line in which the 11 amino acid HiBiT tag[103] has been knocked into the C-terminus of endogenous PrP. This small tag does not itself alter PrP's processing or expression. Following treatment and lysis, addition of HiBiT's complementation partner, LgBiT, to cell lysate generates a luminescent signal in proportion to the PrP present. This signal is dose-dependently decreased by PrP-lowering siRNA treatment in lockstep with PRNP RNA levels as measured by RT-qPCR (Fig. 14). Like HEK293T cells, U251MG cells express *TFRC* and are therefore expected to be transduced by BI-hTfR1v2. As above, cells will be transduced with 5 serial dilutions of the drug product based on a standard number of vector genomes per cell (vg/cell) and signal will be compared to a standard curve.

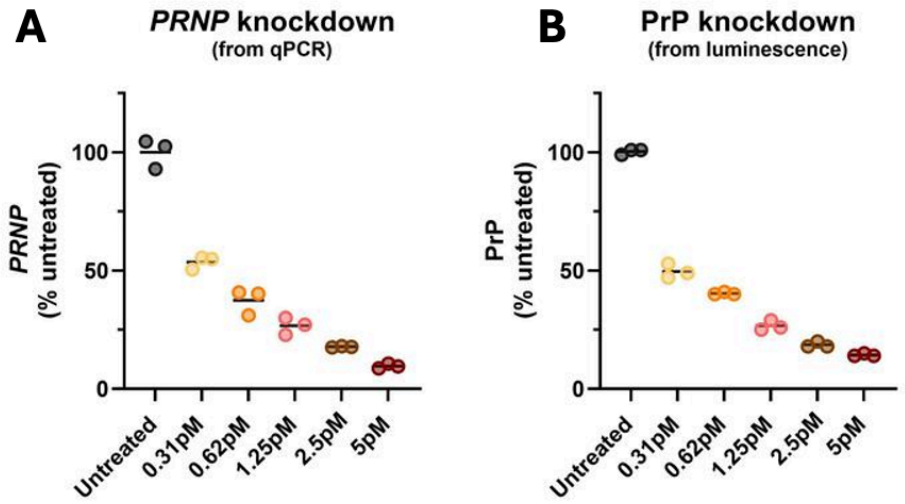


**Figure 14. Comparable dose-dependent reduction of *PRNP* RNA by RT-qPCR and PrP protein by the HiBiT luminescence assay following siRNA treatment of U251MG cells.** U251MG cells were engineered to express PrP in the endogenous locus with a C-terminal 11 amino acid HiBiT tag. Following treatment with PrP-lowering siRNA for 72 hours, cells were lysed and then subjected to A) *PRNP* RNA quantification by RT-qPCR (housekeeping gene TBP), or B) PrP quantification using the HiBiT luminescence assay, per manufacturer's instructions.

Should unanticipated challenges arise in validation of both proposed assays, a less preferred alternative assay would be measurement of the ZF-CHARM transgene via RT-qPCR. This is not our favored approach because ZF-CHARM is effective at low levels of expression, and in design of the final drug candidate we will be seeking to balance broad reach of neuronal cells with low ZF-CHARM expression in any given cell. We therefore anticipate that the transgene may be difficult to detect, even where ZF-CHARM is active.

#### 9.3. PHARMACOLOGY AND TOXICOLOGY INFORMATION

All of the nonclinical studies performed to date are described above. Below we list the additional nonclinical studies we propose to perform prior to IND filing.

##### *9.3.1. Off-target assessment*

In order to focus our analysis of potential off target sites on the human genome, off target analysis will be conducted in a panel of human cell lines confirmed to express both *PRNP* and *TFRC.* Cells will be treated with AAV-formulated PRNP-CHARM-001 and differentially expressed transcripts will be assessed by RNA-seq. AAV dosage will be chosen such that MOI approximates that expected in vivo. Timepoint will be chosen to correspond to full target suppression. If our final human candidate contains a self-silencing mechanism, off-targets will be assessed both during and after CHARM exposure.

Cell lines will be chosen to represent tissues in which both BI-hTfR1v2 uptake and CHARM expression are expected. While ongoing biodistribution and promoter studies may influence this list, based on our present understanding we conditionally nominate the following:
1) iPSC-derived human neurons
2) Human glia-derived cell line
3) Human hepatocyte-derived cell line

##### *9.3.2. In vivo studies*

We propose that the humanized *TFRC* KI mice are the pharmacologically relevant species in which to conduct GLP pharmacology, biodistribution, and toxicology studies, as the BI-hTFR1v2 capsid will be pharmacologically active only in the presence of the human TfR1. As described above, the vector will not show relevant pharmacological activity in other common preclinical species such as wild-type rodents and non-human primates (Figures 12-13).

Note that in all studies below, we will aspire to sex balance each group as feasible based on breeding of transgenic or knock-in lines.
*Definition and justification of terms*

**Mouse lines**

*Table 3: Mouse lines proposed for use in upcoming studies.*

| Abbr. | Mouse description | Appropriate capsid | Appropriate ZF-CHARM | Assays for quantification of brain PrP knockdown |
|---|---|---|---|---|
| *hTFRC* | Mice in which the humanized TfR1 extracellular domain has been knocked into the | BI-hTFR1, BI-hTFR1v2 | mouse | qPCR and ELISA |

| | | | | |
|---|---|---|---|---|
| | mouse locus (human exon 4-19 gene fragment substitution for mouse exon 4-19) [citation] | | | |
| HuPrP | A BAC transgenic mouse line expressing human *PRNP*. | PHP.eB | human | qPCR and ELISA |

**Route**
The route of delivery for all studies listed below is intravenous (IV).

**PrP RNA and protein endpoints**
Where PrP RNA and/or protein are listed as an endpoint, this refers to quantification of PrP RNA and/or protein in whole brain hemisphere homogenates, as we have reported in our ASO development work and all previous studies [see e.g. Minikel 2020[14]]. Because prion disease is a whole brain disease, we consider whole hemisphere to be the most relevant unit of assessment.

*Envisioned in vivo study designs*

**1. Capsid bridging study (non-GLP)**

Since most of our pre-clinical data were generated using PHP.eB as the capsid for delivery, we will perform a bridging study to compare delivery of our lead human ZF-CHARM construct using PHP.eB vs BI-hTFR1v2 (hereafter referred to as BI-hTFR).

This experiment will provide data informing how the PHP.eB doses compare to BI-hTFR1 doses, bridging our earlier preclinical work with PHP.eB to upcoming GLP toxicology and clinical studies, and informing dose calculations for the latter.

We will inject three different doses (low, mid, high) for each construct, plus an uninjected control. Mice will be euthanized at 6wk post-injection and brain and liver tissue will be collected.

| Group | Mouse line | N | Dose (vg/kg) | Capsid | ZF-CHARM | Takedown timepoint | Endpoints |
|---|---|---|---|---|---|---|---|
| **1** | *hTFRC KI* | 8 | vehicle control | N/A | N/A | 6 wk | - Brain PrP RNA + protein<br>- Brain and liver biodistribution |
| **2** | *hTFRC KI* | 6 | 2.00e12 | PHP.eB | mouse | 6 wk | - Brain PrP RNA + protein<br>- Brain and liver biodistribution |
| **3** | *hTFRC KI* | 6 | 1.00e13 | PHP.eB | mouse | 6 wk | - Brain PrP RNA + protein<br>- Brain and liver biodistribution |

| | | | | | | | |
|---|---|---|---|---|---|---|---|
| **4** | *hTFRC KI* | 6 | 5.00e13 | PHP.eB | mouse | 6 wk | - Brain PrP RNA + protein<br>- Brain and liver biodistribution |
| **5** | *hTFRC KI* | 6 | 2.00e12 | BI-hTfR1v2 | mouse | 6 wk | - Brain PrP RNA + protein<br>- Brain and liver biodistribution |
| **6** | *hTFRC KI* | 6 | 1.00e13 | BI-hTfR1v2 | mouse | 6 wk | - Brain PrP RNA + protein<br>- Brain and liver biodistribution |
| **7** | *TFRC KI* | 6 | 5.00e13 | BI-hTfR1v2 | mouse | 6 wk | - Brain PrP RNA + protein<br>- Brain and liver biodistribution |

*Table 4: Proposed study design for capsid bridging study (non GLP).*

**2. PRNP-CHARM-001 potency study (non-GLP)**

Potency of our human ZF-CHARM candidate will be demonstrated across a dose range in huPrP mice in whom target engagement can be assessed. Note that dose levels may shift depending on the results of the capsid bridging study detailed above.

| Group | Mouse line | N | Dose (vg/kg) | Capsid | ZF-CHARM | Takedown timepoint | Endpoints |
|---|---|---|---|---|---|---|---|
| **1** | huPrP | 8 | vehicle control | N/A | N/A | 6 wk | - Brain PrP RNA + protein<br>- Brain and liver biodistribution |
| **2** | huPrP | 6 | 2.00e12 | PHP.eB | human | 6 wk | - Brain PrP RNA + protein<br>- Brain and liver biodistribution |
| **3** | huPrP | 6 | 1.00e13 | PHP.eB | human | 6 wk | - Brain PrP RNA + protein<br>- Brain and liver biodistribution |
| **4** | huPrP | 6 | 5.00e13 | PHP.eB | human | 6 wk | - Brain PrP RNA + protein<br>- Brain and liver biodistribution |
| **5** | wt | 8 | vehicle control | N/A | N/A | 6 wk | - Brain PrP RNA + protein<br>- Brain and liver biodistribution |
| **6** | wt | 6 | 2.00e12 | PHP.eB | mouse | 6 wk | - Brain PrP RNA + protein<br>- Brain and liver biodistribution |
| **7** | wt | 6 | 1.00e13 | PHP.eB | mouse | 6 wk | - Brain PrP RNA + protein<br>- Brain and liver biodistribution |
| **8** | wt | 6 | 5.00e13 | PHP.eB | mouse | 6 wk | - Brain PrP RNA + protein<br>- Brain and liver biodistribution |

*Table 5: Proposed study design for PRNP-CHARM-001 potency study (non GLP).*

Safety assessment endpoints:
Gross morbidity and mortality
Modified Irvin screen (Week 3 and Week 6)

**3. PRNP-CHARM-001 GLP biodistribution and toxicology study**

We propose to conduct our definitive GLP pharmtox study in *TFRC* humanized mice. As our capsid has been shown to bind only the human version of the TfR receptor, it would not be pharmacologically active in any other species.

Note that dose levels may shift depending on the results of the studies detailed above.

This study is designed to assess biodistribution and safety. Target engagement of PRNP-CHARM-001 is not expected in the hTFRC mice, but we have no reason to expect on-target toxicity as *PRNP* is dispensable for healthy life (8.1.4) Meanwhile potential off target sites in the human genome are not assayable in a mouse study but will be assessed in a panel of human cell lines (9.3.1).

At two takedown timepoints – 1 and 3 months – groups of animals from each dose cohort will undergo full necropsy for a standard battery of safety and toxicology endpoints as described in further detail below.

Species: Humanized *TFRC* KI mice
Test Article: Clinical candidate (BI-hTfR1v2 with human ZF-CHARM construct)
Route of administration: Tail vein IV
Dosing frequency: Once
Age: Sexually mature (> 8 weeks)
Number of animals: 80 (~40/sex)

| Group | Mouse line | N | Dose (vg/kg) | Capsid | ZF-CHARM | Takedown timepoint | Endpoints |
|---|---|---|---|---|---|---|---|
| **1** | *hTFRC* | 10 | Vehicle control | N/A | N/A | 1 month | Described below |
| **2** | *hTFRC* | 10 | Minimum anticipated efficacious dose | BI-hTfR1v2 | human | 1 month | Described below |
| **3** | *hTFRC* | 10 | Maximum anticipated efficacious dose | BI-hTfR1v2 | human | 1 month | Described below |
| **4** | *hTFRC* | 10 | 2-5x maximum anticipated efficacious dose | BI-hTfR1v2 | human | 1 month | Described below |
| **5** | *hTFRC* | 10 | Vehicle control | N/A | N/A | 3 months | Described below |
| **6** | *hTFRC* | 10 | Minimum anticipated efficacious dose | BI-hTfR1v2 | human | 3 months | Described below |

| | | | | | | | |
|---|---|---|---|---|---|---|---|
| **7** | *hTFRC* | 10 | Maximum anticipated efficacious dose | BI-hTfR1v2 | human | 3 months | Described below |
| **8** | *hTFRC* | 10 | 2-5x maximum anticipated efficacious dose | BI-hTfR1v2 | human | 3 months | Described below |

*Table 6: Proposed study design for PRNP-CHARM-001 GLP pharmacology and toxicology study.*

Observations and examinations:
- General health: Clinical observations, body weight, and food consumption
- Neurological examinations: Cage side neurological observations (modified Irwin screen) weekly through Day 28 and monthly through Day 91
- Ophthalmology (28- and 91-day cohorts only): prior to dosing and prior to necropsy
- Clinical Pathology: Hematology and Serum Chemistry once at the time of necropsy

Necropsy and Histopathology
- Gross examination
- Organ weights: Full tissue list
- Microscopic examination: To be done by a board-certified Pathologists. Will include a full tissue list for all animals in the high dose group. Read down evaluation of target organs identified in the high dose for tissues from the mid- and low dose groups.

Biodistribution and persistence:
- The following tissues will be collected and evaluated for vector DNA expression by PCR from animals receiving the maximum anticipated efficacious dose (mid-dose group): brain (3 regions), spinal cord (3 regions), gonads, heart, liver, kidney

##### 9.4. SUMMARY OF PREVIOUS HUMAN STUDIES

None.

##### 9.5. SPECIFIC OBJECTIVES/ OUTCOMES EXPECTED FROM MEETING

Concurrence on high-level clinical plan focused on symptomatic patients.
Concurrence on high-level clinical plan for high-risk pre-symptomatic individuals.
Concurrence on nonclinical studies required for this clinical plan.

#### 10. LIST OF QUESTIONS AND SPONSOR POSITIONS

This section provides the questions posed by SPONSOR to the FDA. For ease of review, the questions are provided in **bold font,** and SPONSOR'S position regarding each question is provided in normal font below the question.

**1. [Clinical] We propose to perform parallel trials in symptomatic prion disease patients and pre-symptomatic carriers of high penetrance pathogenic *PRNP* mutations. Does the Agency concur?**

Section V.A (Study Population) of the FDA's Guidance for Industry, Human Gene Therapy Products Incorporating Human Genome Editing (01-2024) recognizes the tradeoffs inherent in treating patients earlier versus later in disease course, stating "Subjects with severe or advanced disease may be more willing to accept the potential risks of an investigational human GE product. However, these subjects may be predisposed to experiencing more AEs [adverse events] or be receiving concomitant treatments, which could make the safety or effectiveness data difficult to interpret. Therefore, in some instances, subjects with less advanced or more moderate disease may be appropriate for inclusion in first-in-human clinical studies."

As described above, symptomatic prion disease patients and presymptomatic high-risk individuals are distinct populations with distinct challenges, opportunities for benefit and prospects for data collection. We aim to honor these complex tradeoffs by performing parallel small N first-in-human studies in both populations.

Symptomatic prion disease patients decline swiftly (median survival time of five months), suffering irreversible neuronal loss. We therefore do not expect to see a signal of efficacy in these patients, especially as the drug is expected to take four weeks to reach full effect at the protein level, during which patients will continue to decline. We further accept that our ability to assess safety in these patients will be limited to acute toxicity and acute immune response. Finally, because CSF PrP declines as a function of the disease course, our biomarker will not provide a reliable readout of target engagement in this population.

By contrast, high-risk genetic carriers, though they do not reliably show prodromal molecular change in advance of symptom onset, face near certainty of fatal disease. Treatment of this population will allow full time for PrP protein reduction and reliable collection of both detailed safety and target engagement data. To manage risk of therapy in this population, enrollment can be biased toward older individuals nearer to expected age of onset for their genotype; toxicology studies will of course also have been performed in humanized mice.

| | **Symptomatic patients** | **Presymptomatic at-risk** |
|---|---|---|
| **Prospect for benefit** | Efficacy not expected. | Prospect to extend healthy life. |
| **Safety** | Acute toxicity and immune response. | Can be assessed in detail. |
| **Target engagement** | Confounded by disease related changes in CSF PrP. | Can be reliably measured. |
| **Prospect for near-term post-mortem tissue analysis** | Yes | No |

| | | |
|---|---|---|
| **Risk vs. benefit** | Reasonable given imminently fatal prognosis. | Reasonable given 90%+ lifetime risk. |

*Table 7. Parameters of symptomatic and presymptomatic treatment populations.*

In summary, the symptomatic and presymptomatic populations are both positioned to benefit from PRNP-CHARM-001, and are sufficiently distinct to merit independent first-in-human studies.

**2. [Clinical] In both symptomatic and presymptomatic trials we propose to measure cerebrospinal fluid (CSF) prion protein levels as a pharmacodynamic biomarker of drug activity in the brain.**

The molecular aim of PRNP-CHARM-001 is to reduce PrP levels in the brain. PrP is the sole essential molecule in prion disease as it serves as substrate for the formation of pathogenic prions; brain PrP dosage is known to control prion formation, replication and neurotoxicity.

Animal data show that CSF PrP levels dose-dependently reflect PrP knockdown in the brain (Fig 2D). In humans, CSF PrP is readily measurable by ELISA and stable for years in healthy individuals (Fig 5; mean CV = 10% over up to six annual measurements), including in high-risk presymptomatic individuals harboring pathogenic *PRNP* variants.

Given the above, we propose that CSF PrP levels will enable clinical assessment of whether the PRNP-CHARM-001 is having its intended molecular effect. Our clinical protocols (Executive Summary) for both symptomatic prion disease patients and high-risk presymptomatic individuals incorporate CSF PrP measurements at enrollment, baseline, and periodically throughout the clinical protocol. In symptomatic patients, CSF PrP data will be collected although disease-related decline in this analyte has been observed absent treatment, and may complicate interpretation. In presymptomatic individuals, drug-dependent changes in CSF PrP levels should be readily accessible given the stable longitudinal baseline in CSF PrP levels observed across genotypes. Thus, while we will measure CSF PrP in both populations, the presymptomatic population offers the stronger opportunity to collect these key data.

**3. [Preclinical] Our novel engineered AAV9-derived viral vector, BI-hTfR1v2, crosses the blood brain barrier by binding the human transferrin receptor. Because this interaction depends on the presence of a human-specific amino acid, BI-hTfR1v2 will not achieve pharmacologically relevant biodistribution in any common preclinical species such as wild-type mice, rats, dogs, minipigs or monkeys (8.3.1). For this reason, we propose that homozygous humanized *TFRC* knock-in mice are appropriate and sufficient as the single species for definitive biodistribution and toxicology studies, and that initial human doses could be selected based on these data. Does the Agency concur?**

In accordance with Section III.B.2 (Animal Species Selection) of the FDA's Guidance for Industry, Preclinical Assessment of investigational Cellular and Gene Therapy Products (11-2013), we have evaluated available animal species and have concluded that because the BI-hTFR1v2 capsid will not distribute to the brain of any other species, humanized *TFRC* KI mice are the appropriate and sole pharmacologically relevant species for definitive biodistribution and toxicology studies of PRNP-CHARM-001.

The BI-hTfR1v2 capsid crosses the blood-brain barrier by binding human TfR1. BI-hTfR1v2 binding to TfR1 requires a single human-specific amino acid, ***. *** is uniquely found in human TfR1 and is absent both in macaque (Figure 12) and every other NHP and domesticated species analyzed (Figure 13). Therefore, the vector will not distribute in these species in a pharmacologically relevant way.

However, BI-hTfR1v2 does efficiently cross the blood-brain barrier in mice in which the relevant portion of TfR1 has been humanized; BI-hTfR1v2 is compatible with both humanized exon 4-19 (humanized extracellular domain) and human apical domain models. Similar humanized *TFRC* KI mice have been successfully used in preclinical studies for TfR1 binding investigational drug products currently in clinical trials (e.g., NCT06075537 and NCT05594992).

Because they support CNS-wide biodistribution and transduction of neurons, which are critical requirements of our therapy, it is our position that the only pharmacologically relevant animal model for evaluating PRNP-CHARM-001 is humanized *TFRC* KI mice.

**4. [Preclinical] Given the lack of homology between the human *PRNP* promoter and that of mouse, the planned human ZF-CHARMs will not have on-target activity in the *TFRC* mice in which we propose to conduct our definitive biodistribution and toxicology studies (8.3.2). We acknowledge that while this study will capture off-target effects of CHARM this study is not designed to capture on-target toxicity. However prion protein is known to be non-essential and we therefore do not expect on-target toxicity from PrP lowering. In order to assess on-target effects we propose to supplement our definitive biodistribution and toxicology studies with separate biodistribution and safety information collected in a non-GLP human PrP-lowering study (9.3.2.2) that will test the human ZF-CHARM candidate in humanized *PRNP* (huPrP) mice. Does the Agency concur?**

As described in Sponsor Position #3, *TFRC* humanized mice are the only preclinical model in which BI-hTfR1v2 will have pharmacologically relevant biodistribution, and we have therefore proposed to use this model for definitive biodistribution and toxicology studies. Given lack of homology between the mouse and human PRNP promoter, we expect that our final human *PRNP*-targeting ZF-CHARM drug candidate will lack homology to the mouse *Prnp* promoter and therefore will not have on-target activity in the humanized *TFRC* mice. As described in Section 8.1.4, PrP is known to be non-essential; knockout data from multiple PrP knockout mouse lines, PrP knockout goats and cows all suggest that total lack of PrP is tolerated. In addition, human genetic data show that loss-of-function *PRNP* alleles arise at the expected rate in the general population, suggesting that natural selection does not act against chance loss of PrP.

For these reasons, it is not expected that reduction of PrP by CHARM, or by any other mechanism, will generate on-target safety concerns directly related to PrP-lowering. Our definitive biodistribution and toxicology study in *TFRC* mice will capture off-target and vector-related toxicity. In addition, we propose to supplement these data by conducting a non-GLP study, using PHP.eB as a surrogate capsid, with the human *PRNP*-targeting CHARM drug candidate in humanized *PRNP* (huPrP) mice (PRNP-CHARM-001 potency study (non-GLP)). This non-GLP study is designed to include safety endpoints. Together, we believe these studies will provide on-target pharmacodynamic impact as well as biodistribution for the clinical capsid.

**5. [CMC] As a Potency Assay for release criteria, we propose to use RT-qPCR-based quantification of *PRNP* RNA in human HEK293T cells. Should an alternative assay be needed, we propose luminescence-based quantification of tagged endogenous PrP in U251MG cells. Does the Agency concur?**

Section V.B.2 (Approaches to Potency Assay Selection and Design) of the FDA's Draft Guidance for Industry, Potency Assurance for Cellular and Gene Therapy Products (12-2023) states that "release testing for a gene therapy vector that expresses a transgene should generally include a potency assay that quantitates transgene mRNA or protein in transduced cells." As described above, we propose to assess potency of our drug product 1) by quantifying *PRNP* RNA levels by RT-qPCR in HEK293T cells, or 2) by quantifying PrP protein levels in U251MG cells by leveraging a HiBiT tag installed at the endogenous protein's C terminus.

Strategy 1: We will transduce HEK293T cells with 5 serial dilutions of the drug product based on a standard number of vector genomes per cell (vg/cell) and perform qRT-PCR to measure expression of prion protein at the RNA level following AAV transduction. Importantly, HEK293T cells robustly express *PRNP* at the RNA level; they also express *TFRC* so should be efficiently transduced by the TfR1 binding AAVs. This assay will determine the correlation between AAV concentration and qPCR PrP detection in cell lysates, including comparison to the signal obtained for an 8 point plasmid dilution series.

Strategy 2: We will use U251MG cells in which endogenous PrP has been modified with an 11 amino acid HiBiT tag at its C terminus. We will transduce these cells with 5 serial dilutions of the drug product based on a standard number of vector genomes per cell (vg/cell) and perform the HiBiT complementation assay to measure expression of prion protein at the protein level following after AAV transduction. U251MG cells express *TFRC* and are efficiently transduced by the TfR1 binding AAVs. This assay will determine the correlation between AAV concentration and HiBiT-based PrP detection in cell lysates, including comparison to the signal obtained for an 8 point plasmid dilution series. Preliminary siRNA treatment data (Fig. 14) indicate that *PRNP* expression in U251MG cells will support a dynamic range sufficient to distinguish batch-to-batch differences in knockdown.

As CHARM is active at low expression levels, we believe that quantification of our target transcript will provide a more robust assay than quantification of transgene expression.

We are requesting feedback on the appropriateness of this proposed assay at the INTERACT stage, instead of waiting for Pre-IND, because assay development and validation require significant lead time and earlier input from the Agency will ensure that these steps do not delay manufacturing.

#### 11. PROPOSED AGENDA

- Introduce participants (5 minutes)
- Sonia Vallabh: brief introduction to this project and our patient community (5 minutes)
- Discussion of questions to the agency and any other input from agency (40 minutes)
- Summary and action items (10 minutes)

#### 12. SPONSOR PARTICIPANTS

Sonia Vallabh, PhD — Broad Institute
Eric Minikel, PhD — Broad Institute
Ben Deverman, PhD — Broad Institute
Jonathan Weissman, PhD — Whitehead Institute
Eric Lander, PhD — Broad Institute
Alissa Coffey, PhD — Broad Institute
Fiona Serack, PhD — Broad Institute
Ken Chan, PhD — Broad Institute
Gregg Cappon, PhD — consultant
Tim Lavaute – NIH
Chris Boshoff – NIH
P.J. Brooks – NIH

#### 13. LIST OF ABBREVIATIONS

| Abbreviation | Definition |
|---|---|
| ASO | antisense oligonucleotide |
| AAV | Adeno-associated virus |
| BAC | Bacterial artifical chromosome |
| BI-hTFR1 | First generation AAV capsid engineered to bind human TfR1 |
| Bi-hTFRv2 | Improved, second generation AAV capsid engineered to bind human TfR1 |
| cDNA | complementary DNA |
| CHARM | Coupled Histone tail for Autoinhibition Release of Methyltransferase |
| CJD | Cruetzfeldt-Jakob diseaes |
| CNS | Central nervous system |
| CpG | cytosine-guanine dinucleotide |
| CSF | cerebrospinal fluid |
| D3A | DNA methyltransferase 3 alpha active catalytic domain |

| | |
|---|---|
| DNMT3A | DNA methyltransferase 3 alpha protein |
| D3L | DNA methyltransferase 3 like C-terminal domain |
| DNMT3L | DNA methyltransferase 3 like |
| DPI | Days post-inoculation |
| ELISA | Enzyme-Linked Immunosorbent Assay |
| FFI | fatal familial insomnia |
| GSS | Gerstmann-Straussler-Scheinker disease |
| hSyn | Human Synapsin promoter fragment |
| huPrP | A BAC transgenic mouse line expressing human PRNP |
| IC | intracerebral |
| ICV | Intracerebroventricular |
| IDMC | Independent Data Monitoring Committee |
| ITR | Inverted Terminal Repeat |
| IV | Intravenous |
| KI | Knock in |
| Luc | Firefly Luciferase |
| mRNA | messenger RNA |
| MRC | Medical Research Council |
| NfL | neurofilament light chain |
| NLS | Nuclear localization signal |
| NGS | Next-generation sequencing |
| NHP | Non human primate |
| PHP.eB | AAV-PHP.eB a surrogate mouse BBB crossing capsid |
| polyA | Poly adenylation sequence |
| PRNP | human gene encoding the Prion protein |
| Prnp | mouse gene encoding the Prion protein |
| PrP | Prion protein |
| RO | retroorbital |
| RLU | Relative light units |
| RT-QuIC | real-time quaking induced conversion |
| RT-qPCR | Real-time quantitative polymerase chain reaction |
| ssAAV | single stranded AAV genome derived from AAV2 |
| SV40pA | Poly adenylation sequence from SV40 |
| Tf | Transferrin |
| TfR1 | Transferrin Receptor 1 protein |
| TFRC | human gene encoding Transferrin Receptor 1 |

| | |
|---|---|
| Tfrc | mouse gene encoding Transferrin Receptor 1 |
| UTR | Untranslated region |
| WPRE | Woodchuck Hepatitis Virus Posttranscriptional Regulatory Element |
| ZF | Zinc finger |
| ZFP | Zinc finger protein |

49. Fornasiero EF, Mandad S, Wildhagen H, Alevra M, Rammner B, Keihani S, Opazo F, Urban I, Ischebeck T, Sakib MS, Fard MK, Kirli K, Centeno TP, Vidal RO, Rahman RU, Benito E, Fischer A, Dennerlein S, Rehling P, Feussner I, Bonn S, Simons M, Urlaub H, Rizzoli SO. Precisely measured

protein lifetimes in the mouse brain reveal differences across tissues and subcellular fractions. Nat Commun. Nature Publishing Group; 2018 Oct 12;9(1):4230.

74. Zeitler B, Froelich S, Marlen K, Shivak DA, Yu Q, Li D, Pearl JR, Miller JC, Zhang L, Paschon DE, Hinkley SJ, Ankoudinova I, Lam S, Guschin D, Kopan L, Cherone JM, Nguyen HOB, Qiao G, Ataei Y, Mendel MC, Amora R, Surosky R, Laganiere J, Vu BJ, Narayanan A, Sedaghat Y, Tillack K, Thiede C, Gärtner A, Kwak S, Bard J, Mrzljak L, Park L, Heikkinen T, Lehtimäki KK, Svedberg MM,

Häggkvist J, Tari L, Tóth M, Varrone A, Halldin C, Kudwa AE, Ramboz S, Day M, Kondapalli J, Surmeier DJ, Urnov FD, Gregory PD, Rebar EJ, Muñoz-Sanjuán I, Zhang HS. Allele-selective transcriptional repression of mutant HTT for the treatment of Huntington's disease. Nat Med. 2019 Jul;25(7):1131–1142. PMID: 31263285

92. Ullman JC, Arguello A, Getz JA, Bhalla A, Mahon CS, Wang J, Giese T, Bedard C, Kim DJ, Blumenfeld JR, Liang N, Ravi R, Nugent AA, Davis SS, Ha C, Duque J, Tran HL, Wells RC, Lianoglou S, Daryani VM, Kwan W, Solanoy H, Nguyen H, Earr T, Dugas JC, Tuck MD, Harvey JL, Reyzer ML, Caprioli RM, Hall S, Poda S, Sanchez PE, Dennis MS, Gunasekaran K, Srivastava A, Sandmann T, Henne KR, Thorne RG, Di Paolo G, Astarita G, Diaz D, Silverman AP, Watts RJ, Sweeney ZK, Kariolis MS, Henry AG. Brain delivery and activity of a lysosomal enzyme using a blood-brain barrier transport vehicle in mice. Sci Transl Med. American Association for the Advancement of Science; 2020 May 27;12(545):eaay1163.

93. Hammond SM, Abendroth F, Goli L, Stoodley J, Burrell M, Thom G, Gurrell I, Ahlskog N, Gait MJ, Wood MJA, Webster CI. Antibody-oligonucleotide conjugate achieves CNS delivery in animal models for spinal muscular atrophy. JCI Insight [Internet]. American Society for Clinical Investigation; 2022 Dec 22 [cited 2024 Nov 11];7(24). Available from: https://insight.jci.org/articles/view/154142 PMID: 0

94. Kariolis MS, Wells RC, Getz JA, Kwan W, Mahon CS, Tong R, Kim DJ, Srivastava A, Bedard C, Henne KR, Giese T, Assimon VA, Chen X, Zhang Y, Solanoy H, Jenkins K, Sanchez PE, Kane L, Miyamoto T, Chew KS, Pizzo ME, Liang N, Calvert MEK, DeVos SL, Baskaran S, Hall S, Sweeney ZK, Thorne RG, Watts RJ, Dennis MS, Silverman AP, Zuchero YJY. Brain delivery of therapeutic proteins using an Fc fragment blood-brain barrier transport vehicle in mice and monkeys. Sci Transl Med. American Association for the Advancement of Science; 2020 May 27;12(545):eaay1359.

95. Rué L, Jaspers T, Degors IMS, Noppen S, Schols D, De Strooper B, Dewilde M. Novel Human/Non-Human Primate Cross-Reactive Anti-Transferrin Receptor Nanobodies for Brain Delivery of Biologics. Pharmaceutics. 2023 Jun 16;15(6):1748. PMCID: PMC10300862

96. Yu YJ, Atwal JK, Zhang Y, Tong RK, Wildsmith KR, Tan C, Bien-Ly N, Hersom M, Maloney JA, Meilandt WJ, Bumbaca D, Gadkar K, Hoyte K, Luk W, Lu Y, Ernst JA, Scearce-Levie K, Couch JA, Dennis MS, Watts RJ. Therapeutic bispecific antibodies cross the blood-brain barrier in nonhuman

primates. Sci Transl Med. American Association for the Advancement of Science; 2014 Nov 5;6(261):261ra154-261ra154.
